# Nyx: a flexible framework for sleep scoring across species, lifespan and modalities

**DOI:** 10.64898/2026.07.24.740558

**Authors:** Letizia Signorelli, Solomiia Korchynska, Cantin Ortiz, Alessandro Arena, Sophia Wilhelm, Charlotte Boccara

## Abstract

Sleep is a key factor in almost every physiological process, yet scoring tools lag behind innovations in basic to clinical research. New automated algorithms are trained on narrow datasets, typically healthy adult mouse or human, and fail to generalise, leaving labs to fall back on slow, laborious, and often poorly reproducible manual scoring. Here we introduce Nyx, an unsupervised framework built on principal component analysis that flexibly clusters epochs into sleep states, akin to how spike-sorting algorithms cluster single-neuron activity. Validated on about 8,000 hours spanning 19 independent datasets, it generalises across species, lifespan, recording modalities, and altered sleep architectures, matching human inter-scorer agreement. Beyond this, Nyx opens new ground for modern sleep research: classifying NREM subtypes in rodents, revealing states that cut across canonical boundaries in non-mammalian species, and enabling real-time, closed-loop manipulation. Distributed as an open-source package with a graphical user interface, Nyx offers an accessible, explainable framework built to scale with the pace of innovation itself.

## Main

Sleep is a key factor in nearly every major biological process, and mounting evidence links its disruption to long-term harm to brain health. Yet the way sleep is measured remains largely unchanged since Rechtschaffen and Kales first codified epoch-by-epoch scoring rules in 1968^1^, a legacy that has constrained the field for decades. Manual scoring by trained experts remains the standard, but it is time-consuming and inherently subjective. Inter-scorer variability is well documented in both rodent and human recordings^2,3^, with the largest disagreements concentrated around stage transitions and at contested stages such as non-rapid eye movement stage 1 (N1) and rapid eye movement (REM) sleep^4^. As interest grows and recording tools multiply, sleep measurement extends into labs without specialist training, and data accumulates faster than it can be scored. Many researchers then reach for simple, unreliable proxies – a theta-to-delta ratio from a single channel, for instance^5^ – or fall back on manual scoring itself.

A steady stream of automated sleep scoring algorithms has been developed^6^ to address this scalability and reliability problem^3,7–12^, yet none are currently creating a consensus within the community due to several limitations. Most are supervised classifiers trained on expert-labelled data — such as Somnotate^3^ for rodents and YASA^9^ for humans —, and while they reduce annotation burden, they inherit the conventions and biases of their reference labels, generalise poorly outside the contexts in which they were trained, and function as black boxes that offer little insight into stage assignments^3,8–11^. Unsupervised approaches, on the other hand, are sparser and species-specific: FASTER uses density-based and hidden Markov clustering on EEG/EMG features in mice^13,14^, and the recent AISleep applies UMAP and kernel density estimation to single-channel EEG in humans^15^. A problem consistent across both supervised and unsupervised approaches is that they have been built and validated within a single species or recording context. These limitations matter increasingly as sleep research expands across diverse species, developmental stages, health status and recording modalities (Fig. 1c). No current algorithm is designed to operate jointly across this full space, leaving both cross-context generalisation and the discovery of novel sleep structures out of reach.

**Fig. 1:**
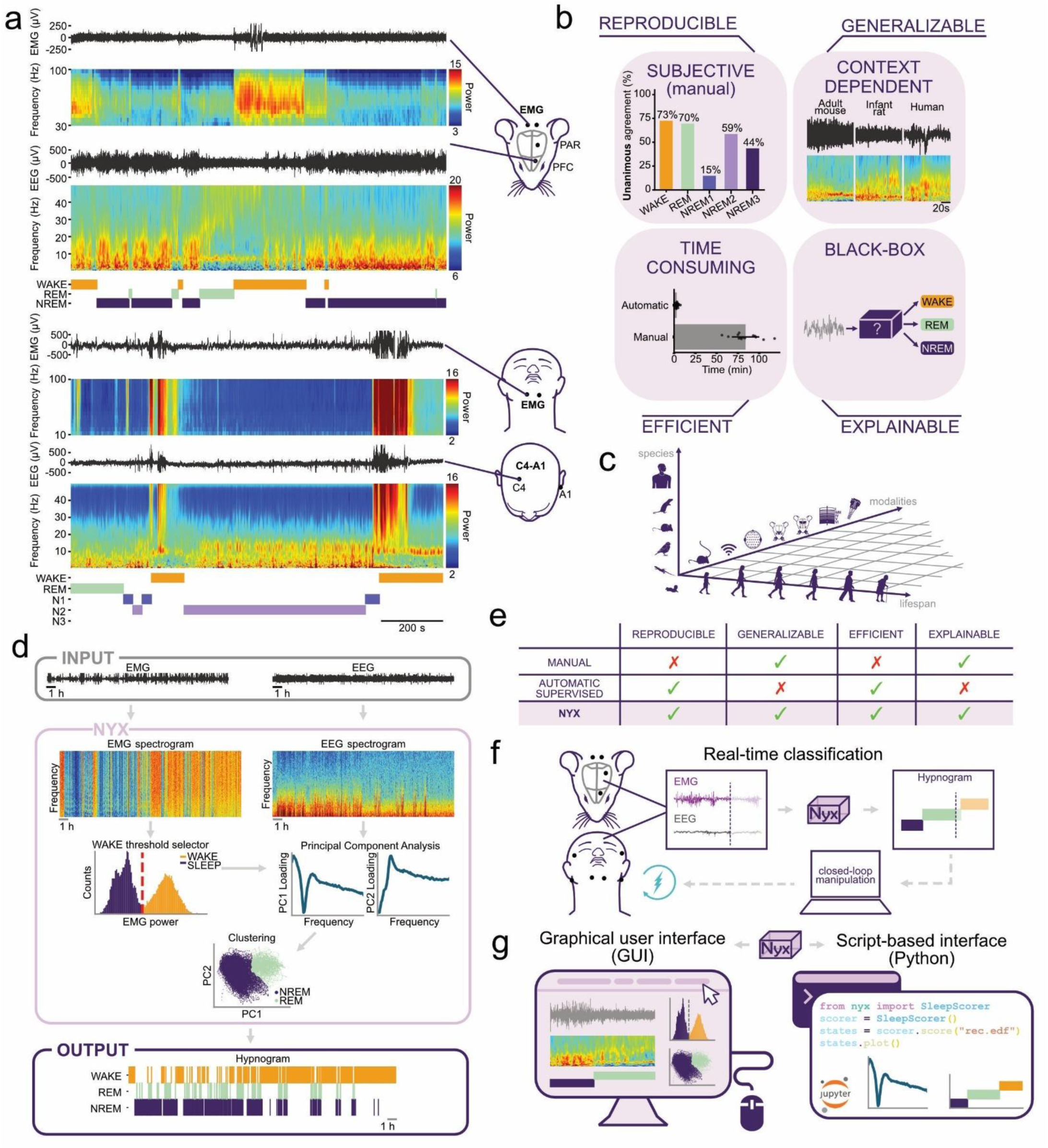
Designing Nyx for reproducible, generalisable, explainable, and efficient sleep scoring **a,** Representative scoring of a rat and a human recording. From top: raw EMG trace, EMG time–frequency spectrogram, raw EEG trace, EEG time–frequency spectrogram, and Nyx (automated) hypnogram, illustrating the sleep-stage sequence assigned by the algorithm. WAKE in orange, REM in green, NREM in purple. **b,** The four challenges of current sleep scoring approaches (inner labels) and the corresponding properties a unified framework should satisfy (outer labels). Subjective: even five experienced scorers reach unanimous agreement on only 73% of WAKE, 70% of REM, and as little as 15% of N1 epochs in human polysomnography (DOD-H and DOD-O; Fig. S2, S3). Context-dependent: signal features vary markedly across species, ages, and recording modalities (example traces and spectrograms from adult mouse, infant rat, and human). Time-consuming: manual scoring takes ∼80 min per 8-h recording vs minutes for automated methods (in house comparison based on 5 scorers). Black-box: supervised methods provide no insight into how epochs are assigned to stages. A unified framework should jointly satisfy four properties: reproducibility, generalisability, efficiency, and explainability. **c,** The three-dimensional space across which Nyx enables sleep research to operate: species (humans, rodents, birds, reptiles), developmental stages (early postnatal to advanced age), and recording modalities (tethered, wireless, EEG-cap; surface and intracortical). **d,** Nyx pipeline. From a single EEG channel (or equivalent neural signal) and an EMG (or equivalent motor-activity signal), per-epoch spectral features are extracted, projected into a low-dimensional space via PCA, and assigned stages by unsupervised clustering (K-means, Gaussian mixture models, or HDBSCAN). No pre-labelled training data are required. **e,** Comparison of manual scoring, supervised automated methods, and Nyx against the four properties defined in b. Nyx is the only framework to satisfy all four simultaneously. **f,** Real-time scoring extension: incoming epochs are projected onto a PCA space computed on an earlier baseline session, allowing online stage assignment without re-fitting. **g,** Distribution: Nyx is released as a Python package with both a code notebook and a graphical user interface (GUI), supporting use by researchers without a dedicated computational infrastructure.

Addressing these challenges calls for a sleep scoring framework that is at once reproducible, generalisable, and efficient, while remaining explainable enough for researchers to trust its output (Fig. 1b). Here we introduce Nyx, an unsupervised framework designed to satisfy these four properties jointly (Fig. 1d-e). Nyx extracts spectral features from neural and motor-activity signals, projects them into a low-dimensional space via Principal Component Analysis (PCA) and assigns stages by flexible clustering approaches. Conceptually akin to how spike-sorting algorithms (e.g., Kilosort^16^) recover their units from the recording itself without external labels, Nyx treats sleep states the same way: structure that emerges from the data, not a category learned from examples, nor one imposed by rigid predefined rules. Distributed as both an open-source Python package and a graphical interface, Nyx is accessible to researchers without dedicated computational support (Fig. 1g). We validate this framework on close to 8,000 hours of recordings across 19 independent datasets, spanning species, developmental stages, recording modalities, and altered sleep architecture, and show that it recovers conventional sleep states with agreement comparable to inter-scorer reliability. This generalisability alone addresses a limitation no existing method currently overcomes. Beyond this, Nyx further supports analyses that are out of reach for most current alternatives. Specifically, it can be used for real-time stage assignment for closed-loop applications and classifies subtypes and sleep-state structure in non-conventional species that cuts across canonical boundaries (Fig. 1f,g).

## Results

### Designing Nyx for reproducible, generalisable, and explainable sleep scoring

Nyx is built on a thoughtfully simple pipeline: spectral features, a low-dimensional projection, and flexible clustering (Fig. 1d). It requires as input only a single EEG channel (or an equivalent neural activity signal) and an EMG channel, or an equivalent motor-activity signal such as accelerometer or video pose-estimation data. From each epoch it extracts spectral features, projects them into a low-dimensional space via Principal Component Analysis (PCA), and assigns stages by unsupervised clustering using K-means, Gaussian mixture models, or HDBSCAN. Both the spectral features driving the projection and the power spectra of the resulting clusters are directly inspectable (Fig. S1), making every stage assignment explainable. Equally, no a priori assumptions about sleep architecture or forbidden transitions between stages are imposed: epochs group by their position in feature space, and stage identities are assigned only once the clusters have emerged. This flexibility is by design: tuneable key parameters, supported by built-in selection metrics, allow Nyx to be adapted to a wide range of recording contexts and to explore stage structures beyond canonical mammalian architecture. Representative outputs in a rat and a human recording illustrate the resulting stage sequences (Fig. 1a). When compared with manual and supervised approaches across the four properties of reproducibility, generalisability, efficiency, and explainability (Fig. 1b), Nyx is the only framework to satisfy all of them simultaneously (Fig. 1e). The same framework extends to real-time scoring by projecting incoming epochs onto a baseline PCA space computed on an earlier recording session (Fig. 1f), and is distributed as a Python package with both a code notebook and a graphical user interface (GUI) for researchers without dedicated computational support (Fig. 1g).

### Nyx achieves reliable sleep scoring across rodent datasets and conditions

To validate Nyx on standard rodent EEG, we compared its outputs against expert manual annotation across 10 independent rodent datasets covering 179 recordings from 103 rats and 76 mice (Table S1). Representative traces, hypnograms, and stage-coloured projections in PCA space illustrate the close correspondence between manual and automated scoring, with discrepancies concentrated at cluster boundaries (Fig. 2a).

**Fig. 2:**
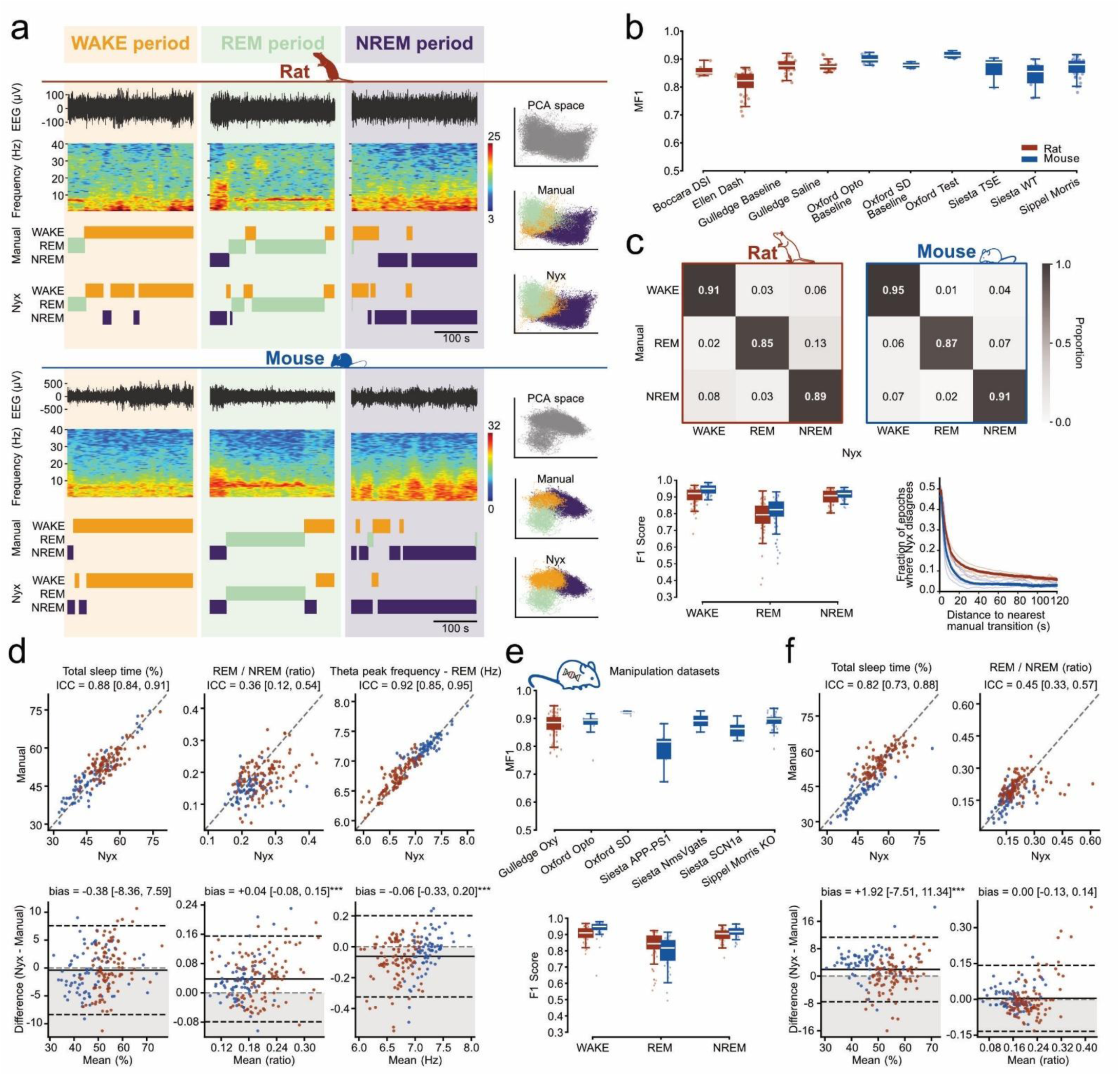
Nyx achieves reliable sleep scoring across rodent datasets and conditions **a,** Representative scoring of one rat and one mouse recording. From top: raw EEG trace, time–frequency spectrogram, manual and Nyx (automated) hypnograms; Right: PC-space projection of all epochs (top) coloured by manual (centre) and Nyx (bottom) labels. **b,** Per-recording macro F1 across the ten baseline rodent datasets (n = 179 recordings from 103 rats and 76 mice; full dataset descriptions in Table S1). **c,** Epoch-level structure of agreement under the 3-stage scheme. Top: Confusion matrices pooled across recordings per-species. Bottom left: per-stage F1 distributions across recordings, split by species, showing REM as the most challenging stage in both rats and mice. Bottom right: fraction of disagreed epochs as a function of distance from the nearest manual stage transition, peaking at ∼50% at the boundary and decaying to a low plateau (< 10% within 14 s in mice and 39 s in rats). Per-species inter-scorer reference in Fig. S2. **d,** Agreement on summary sleep variables across all baseline recordings. Scatter (top) and Bland– Altman (bottom) pairs for total sleep time, REM/NREM ratio, and theta peak frequency in REM, (Nyx on *x,* manual on *y*; bias convention: Nyx − manual). Grey shading marks the negative half of the *y*-axis to highlight the zero line. ICC(A,1) values inset. **e,** Top: Per-recording macro F1 across the seven manipulation datasets (n = 171 recordings; genetic, pharmacological, optogenetic, and behavioural manipulations; Table S1). Bottom: per-stage F1 distributions across recordings, split by species, same as c. **f,** Agreement on summary sleep variables across manipulation recordings, same layout as d. Boxplots show median, IQR, and 1.5 × IQR whiskers. Rats are shown in red and mice in blue throughout.

We first asked whether Nyx delivers consistent performance across the diversity of rodent datasets included. Per-recording macro F1 was consistently high across datasets (median range 0.84–0.93; Fig. 2b, Table S3), with no systematic effect of laboratory and only one modest outlier^17,18^ indicating that Nyx generalises robustly across research groups and recording configurations without dataset-specific tuning. We next examined whether this dataset-level agreement was matched at the epoch level, where agreement was high in both species (Cohen’s κ = 0.82 in rats, 0.85 in mice; Table S2). Per-stage F1 was high for WAKE and NREM (> 0.89 in both species), with REM slightly lower (F1 ≈ 0.78–0.80; Fig. 2C, Fig. S2A).

REM recall consistently exceeded REM precision, indicating that Nyx tends to assign more epochs to REM than the manual scorer. REM transitions, however, are also notoriously ambiguous to score, suggesting the manual reference itself may be less reliable in this specific window. Mismatches were not randomly distributed in time: disagreement peaked at ∼50% at the transition boundary itself and decayed rapidly, falling below 10% within 14 s in mice and 39 s in rats before stabilising at a low plateau (Fig. 2c). This pattern mirrors the structure of inter-scorer disagreement (Fig. S2), indicating that residual errors concentrate at the same inherently ambiguous epochs that limit human-human agreement rather than reflecting a systematic bias of the algorithm. Crucially, even compared against an established supervised benchmark (Somnotate; Brodersen et al., 2024^3^) trained on its original mouse dataset and applied without retraining, Nyx matched or exceeded per-animal macro F1 (Wilcoxon signed-rank; ***p < 0.001; Fig. S4B, Table S4).

Because sleep scoring is the starting point for nearly all downstream analyses, we next asked whether the sleep variables most commonly reported in the field could be reliably derived from Nyx outputs. Total sleep time, theta peak frequency in REM, and delta power in NREM all showed good to excellent absolute agreement with negligible bias (Fig. 2d, Table S5). REM/NREM ratio showed lower absolute agreement (ICC = 0.36 [0.12, 0.54]), reflecting a small but consistent positive bias (+0.04 ± 0.06) inherited from the epoch-level REM overscoring identified above. Applying REM post-hoc rules (see Supplementary Methods) reduced this bias to +0.01 ± 0.06 (Fig. S6a–d). Microarousals, by contrast, were consistently undercounted (bias −2.17 events/hour; Fig. S4b, Table S5). This reflects the default epoch settings used, which were optimised for stage scoring rather than microarousal detection: brief WAKE bouts blend into the surrounding stage’s features within the EMG epoch window, and would be recoverable with a shorter EMG epoch and smoothing — a direction we return to in the Discussion.

To confirm that automated scoring captures biologically meaningful experimental effects without compromising performance, we additionally analysed 7 datasets spanning 171 recordings under genetic, pharmacological, optogenetic, and behavioural manipulations (Table S1). Per-recording macro F1 was broadly comparable to baseline (Fig. 2e), with one expected exception: the Siesta APP-PS1 dataset³, an Alzheimer’s model with abnormal EEG architecture, showed slightly lower and more variable performance (median MF1 = 0.82). Epoch-level agreement and per-stage F1 distributions across the remaining manipulations matched the baseline pattern, with REM remaining the most variable stage (Fig. 2e, Table S5). At the sleep-variable level, total sleep time acquired a very small positive bias (+1.92%) but retained good reliability; the baseline REM/NREM overestimation resolved (bias 0.004, p = 0.135); and theta peak frequency and NREM delta power remained in excellent agreement (Fig. 2f, S4c, Table S5). Microarousals were the only metric to degrade further (bias −6.40 events/hour, ICC = 0.11), reflecting the same default epoch-setting constraint amplified under altered sleep architecture.

Together, these results show that Nyx delivers reliable rodent sleep scoring across laboratories, species, and manipulations — matching or outperforming established benchmarks — while supporting the downstream sleep variables most commonly used in the field.

### Nyx generalises to human sleep scoring from coarse to fine AASM resolution

To validate Nyx on human polysomnography, we compared its outputs against expert manual annotation across five datasets — ANPHY, DOD-H, DOD-O, MESA (overnight recordings, ∼8 h), and SleePing (daytime naps, ∼90 min) — encompassing both healthy and clinical (DOD-O, Obstructive Sleep Apnea suspicion) populations (Table S1). Recordings with poor signal quality were excluded (see Methods). Following Sigurdardottir et al.⁴, Nyx was evaluated under 3-, 4-, and 5-stage schemes defined by the American Academy of Sleep Medicine (AASM) scoring manual^19^, spanning from the coarsest resolution — matching rodent scoring — to the full AASM granularity. Representative outputs for one recording illustrate the close correspondence between manual and automated scoring (Fig. 3a): WAKE, REM, N2 and N3 clusters partition cleanly in PCA space, while N1 epochs sit dispersed at the interfaces between WAKE, REM and N2, further motivating the multi-scheme comparison below.

**Fig. 3:**
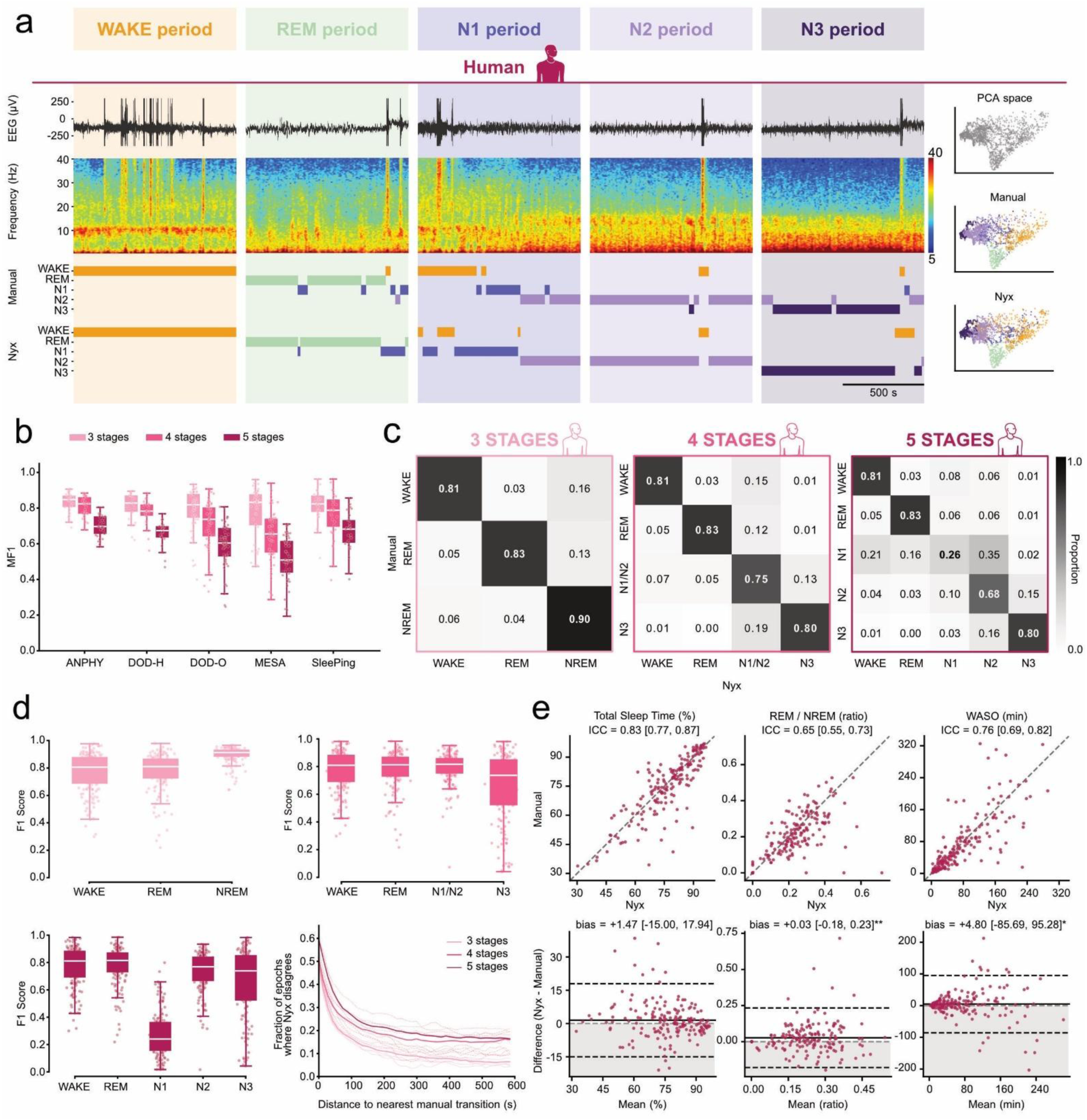
Nyx generalises to human sleep scoring from coarse to fine AASM resolution **a,** Representative epochs from a single polysomnography recording. Left: Examples of raw EEG and time– frequency spectrogram for each stage, arranged left to right as WAKE, REM, N1, N2, N3, with the corresponding manual and Nyx labels shown beneath. Right: PC-space projection of all epochs from the recording (top), coloured by manual (centre) and Nyx (bottom) labels under the 5-stage scheme. WAKE, REM, N2, and N3 partition cleanly; N1 epochs sit dispersed at the interfaces. **b,** Per-recording macro F1 across the five datasets (ANPHY, DOD-H, DOD-O, MESA, SleePing) under the 3-, 4-, and 5-stage AASM schemes. Recording counts per dataset in Table S1. Stage schemes shown in light-to-dark pink for 3-, 4-, and 5-stage respectively. **c,** Confusion matrices pooled across recordings under the 3-, 4-, and 5-stage AASM schemes, showing preserved diagonal mass at 3 and 4 stages and the N1-specific collapse at 5 stages (F1 = 0.28), with confusion concentrated against N2. **d,** Per-stage F1 pooled across datasets under each scheme, showing preserved performance for all stages from 3 to 4 stages, and the N1-specific collapse at 5 stages (F1 = 0.27). Bottom-right: fraction of disagreed epochs as a function of distance from the nearest manual stage transition, showing the steep fall and plateau characteristic of transition-localised errors. **e,** Agreement on summary sleep metrics across all recordings. Scatter (top) and Bland–Altman (bottom) pairs for total sleep time, REM/NREM ratio, and WASO (Nyx on x, manual on y; bias convention: Nyx − manual). Grey shading marks the negative half of the *y*-axis to highlight the zero line. ICC(A,1) values inset. Boxplots show median, IQR, and 1.5 × IQR whiskers.

We first asked whether Nyx provides reliable human scoring at the dataset level. Per-recording macro F1 was consistent across datasets within each scheme but dropped progressively with finer stage resolution (Fig. 3b; Table S6): from 0.82 at 3 stages, to 0.74 at 4 stages, and 0.62 at 5 stages. Unsurprisingly, the drop between 4 and 5 stages was driven almost entirely by the introduction of N1 as a separate class – mirroring the poor interrater agreement for N1 detection. MESA was the most affected dataset at 5 stages, likely reflecting a combination of lower signal quality, and a more clinically heterogeneous population, as also documented in benchmarking studies of this dataset⁵.

We next examined stage-level structure within each scheme. At 3 stages, agreement was high across all stages (per-stage F1 in the 0.77–0.90 range; Fig. 3c–d; Table S6). As in rodents, REM recall exceeded REM precision, again reflecting a tendency to assign more epochs to REM than the manual scorer. Merging N1 with N2 into a combined light-sleep class (4 stages) preserved this per-stage performance; a sharp drop emerged only when N1 was separated as its own class (5 stages), where its F1 collapsed to 0.27 (Fig. 3c–d). All other stages remained well above this level across schemes, confirming N1 as a categorical outlier potentially encompassing a mix of substates. As in rodents, disagreement localised to stage transitions, with the fraction of disagreed epochs falling steeply with distance from the nearest transition before plateauing (Fig. 3d, bottom-right). Against YASA (Vallat & Walker, 2021^9^), a widely used supervised human sleep-scoring tool, Nyx scored higher on ANPHY while YASA scored higher on SleePing (both p < 0.001, Wilcoxon signed-rank). Pooled across datasets, however, the difference was not significant (p = 0.89). The pattern held across 3-, 4-, and 5-stage schemes (Fig. S5a; Table S8). Beyond stage-level agreement, we asked whether the summary sleep metrics commonly reported in human sleep research could be reliably recovered from Nyx outputs. Total sleep time, Wake After Sleep Onset (WASO) and REM/NREM ratio were reproduced with quite good reliability (ICC = 0.65–0.83; Fig. 3e, Table S9), with WASO and REM/NREM ratio carrying small positive bias (+4.8 min and +0.03 respectively). Spectral features were near-perfectly recovered (sigma power in N2, delta power in N3, theta power in REM; ICC ≥ 0.999, |bias| < 0.7 dB; Fig. S5, Table S9). The poorest agreement was for REM latency (ICC = 0.28), reflecting its dependence on identifying both the first sleep epoch and the first REM epoch. Because Nyx assigns stages purely from cluster structure, without hard-coded rules for sleep onset or REM transitions, REM latency is particularly sensitive to small scoring differences at the beginning of the recording. This is driven by the high similarity of REM and WAKE signals – a boundary on which manual scorers themselves frequently disagree, partly due to rigid transition rules (e.g., forbidding direct WAKE-to-REM) that don’t always hold true. REM post-hoc rules (Fig. S6e–h) reduced this bias (from −35.1 min to −21.8 min under the strongest rule; p = 0.005).

Together, these results show that Nyx delivers reliable human sleep scoring across overnight as well as daytime nap recordings, in both healthy and clinical adult populations, and the 3-and 4-stage AASM schemes routinely used in sleep research, with N1 reflecting the inherent ambiguity of a stage on which human scorers themselves disagree (Fig.S3 ^2^). Remarkably — despite receiving no dataset-specific training — Nyx performed on par with a leading supervised classifier developed specifically for human data (YASA^9^).

### A single framework across three dimensions: species, lifespan, and recording configuration

A central challenge for automated sleep scoring is generalisation: across species, developmental stages, and recording configurations, sleep architecture varies substantially, yet most algorithms are tied to the contexts in which they were trained. Nyx’s unsupervised design relaxes this constraint, since the algorithm adapts to the structure of each recording rather than relying on fixed stage definitions and scoring predefined rules. We set out to test Nyx flexibility across three orthogonal axes of variation (Fig. 4a): the rodent and human lifespan, additional species, and recording configurations spanning cortical regions, recording depths (intracortical and hippocampal), EEG derivations, and EMG-free setups.

**Fig. 4:**
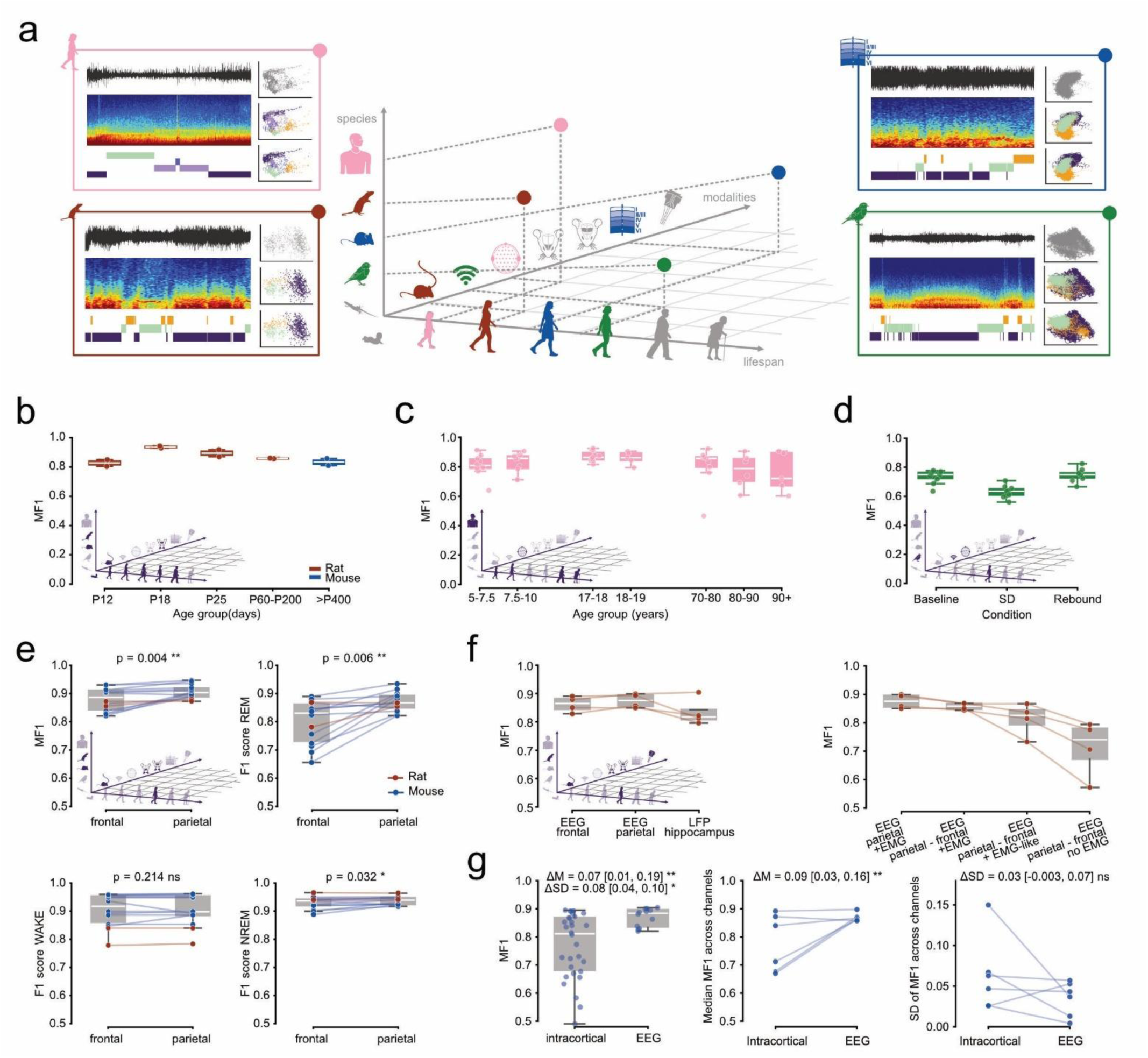
Nyx generalises across species, lifespan, and recording configuration **a,** Schematic of the three orthogonal axes tested: lifespan (rodent and human), species (jackdaw), and recording configuration (tethered, wireless, and EEG-cap setups; frontal and parietal derivations; surface and intracortical/tetrode depths) with example datasets that have been tested for this publication. **b,** Macro F1 across the rodent lifespan at P12, P18, P25, adult rats, and ageing mice (total n = 8, n = 2 individuals per group). **c,** Macro F1 across the human lifespan under the 3-stage scheme, pooled across CHAT, CCSHS, and MESA (n = 60 recordings across 7 age bins, 5–7.5 y to 90+ y). 4-and 5-stage equivalents in Fig. S7A. **d,** Macro F1 in jackdaws across a sleep-deprivation protocol (baseline, sleep deprivation, rebound; n = 9 birds, 9 recordings per condition)^21^. Per-stage F1 and hemisphere comparison in Fig. S7B. **e,** Paired comparison of frontal vs parietal EEG derivations (n = 14 pairs: 12 mouse baseline recordings from the Oxford dataset and 2 adult rats from in-house tethered recordings). Slopegraph of macro F1 with paired bootstrap testing. **f,** Effect of recording configuration in n = 4 adult rats with simultaneous frontal EEG, parietal EEG, and hippocampal LFP. Left: single-channel scoring across the three derivations (frontal, parietal, hippocampal), each paired with the recorded EMG. Right: parietal (reference) and parietal–frontal bipolar derivations, with the parietal–frontal derivation paired with the recorded EMG, with an EMG-like surrogate derived from LFP/ECoG (see Methods), or with no EMG. Confusion matrices in Fig. S7c. **g,** Surface EEG vs multi-channel intracortical scoring in n = 12 mice from the Oxford sleep-deprivation dataset with paired surface EEG (2 channels) and intracortical probes (5 channels each). Left: distribution of channel-level F1 pooled across recordings (intracortical n = 30, EEG n = 12). Annotations give the pooled/marginal between-modality differences in median (ΔM) and standard deviation (ΔSD) of F1, with 95% bootstrap CIs in brackets and two-sided bootstrap significance (∗ p < 0.05, ∗∗ p < 0.01). Centre and right: within-recording median F1 standard deviation of F1 across channels, per recording. Each line links the two modalities for one recording. Statistics are cluster bootstraps over recordings (10,000 iterations; see Methods). Boxplots show median, IQR, and 1.5 × IQR whiskers; slopegraphs connect paired recordings. Paired bootstrap p-values are Holm-corrected within panel e.

We first assessed whether Nyx’s performance was preserved across the rodent lifespan, from early postnatal development to aging (Fig. 4b; Table S10). Macro F1 remained high at every age sampled (range 0.83–0.94 across 8 animals spanning five age groups: P12, P18, P25, adult rats, and ageing mice; n = 2 per age), indicating that the unsupervised approach adapts to age-related changes in sleep architecture. Nyx clearly outperformed Somnotate across the lifespan cohort, where the pre-trained supervised algorithm was applied outside its training distribution (median MF1 0.86 vs 0.69; Nyx > Somnotate in all 8 animals; Fig. S7a, Table S11). We next applied Nyx to human polysomnographic recordings spanning childhood (5–7.5 y) to advanced age (90+ y), pooled across the CHAT, CCSHS, and MESA datasets (n = 60 recordings across 7 age bins; Table S10). At the 3-stage resolution, macro F1 remained high across the lifespan (mean 0.82 ± 0.09, range 0.76–0.87 across age bins), with no systematic decline from childhood to advanced age (Fig. 4c). Under the 4-stage scheme, performance was likewise preserved across age groups except for an age-specific drop in N3 in adults ≥ 70 y, consistent with known age-related attenuation of slow-wave activity leading to reduced spectral distinction and increased ambiguity of the N2/N3 boundary^20^.The same N3 pattern carried over to the 5-stage scheme (Fig. S7b), where N1 remained the only stage with uniformly low F1 across all age groups, as in the main validation. Together, these results show that Nyx unsupervised approach uniquely generalises across the rodent and human lifespan, capturing age-related changes in sleep architecture across mammals.

A more challenging test of generalisation is non-mammalian sleep, where stages are functionally analogous to mammalian NREM and REM but supported by very different cortical anatomy^21^. To our knowledge, no automatic sleep scoring algorithm has yet been explicitly tested in this setting. We applied Nyx to jackdaw recordings spanning an entire sleep-deprivation protocol, from baseline to recovery (Fig. 4D; n = 9 birds, 9 recordings per condition). Macro F1 was preserved between matched undisturbed conditions (0.73 ± 0.04 at baseline; 0.75 ± 0.04 during recovery; Fig. 4d; Table S10). Performance was further robust to hemisphere choice, with left, right, and combined-channel analyses yielding near-identical macro F1 (0.70 ± 0.07). Performance dropped during sleep deprivation (MF1 = 0.63 ± 0.05), but this reduction was driven entirely by REM (Fig. S7c), whose very low base rate under sleep deprivation amplifies the F1 impact of even a handful of misclassifications — overall accuracy was in fact higher during sleep deprivation (0.91) than at baseline (0.85). These lower F1 scores should nonetheless be interpreted with caution: REM is particularly difficult to identify in this species, potentially introducing imprecision into the reference manual scoring itself. We return to this point later, where we examine whether unsupervised stage definitions tailored to non-mammalian sleep architectures can recover finer structure than the mammalian three-stage scheme permits.

Researchers face a wide range of experimental constraints — cortical region of interest, electrode depth, or available equipment, among others — that often prevent recording under the ideal conditions required by standard scoring pipelines. However, this is rarely considered when building scoring algorithms. We therefore asked which recording choices matter for Nyx in practice. Specifically, we tested whether scoring quality depends on the cortical region, the depth of the electrode, the choice of derivation, or the availability of an EMG channel, and how much each choice impacts Nyx accuracy. We first compared frontal and parietal EEG derivations using a paired design (n = 14 pairs; Methods). Twelve pairs came from baseline mouse recordings in the Oxford sleep-deprivation dataset; the remaining two pairs came from adult rats with simultaneous frontal and parietal EEGs from our in-house tethered recordings. Both derivations supported high-quality scoring (median MF1 = 0.89 vs 0.90; Fig. 4e; Table S12), with an expected small but consistent unsurprising advantage for parietal (p = 0.004 on macro F1). The advantage was driven primarily by REM, where the median F1 difference reached 0.04 (frontal 0.83 vs parietal 0.87; p = 0.006). NREM F1 also showed a significant paired difference (p = 0.032) but in absolute terms the gap was negligible (0.938 vs 0.941). This pattern is consistent with the stronger posterior visibility of theta rhythms that anchor REM identification, and indicates that both derivations are usable for Nyx scoring, with parietal providing marginally better separation of the REM cluster. Beyond cortical region, sleep recordings may vary in the depth of the electrode itself. We tested whether Nyx accommodates this variation across hippocampal LFP and intracortical channels. In four adult rats from our in-house dataset with simultaneous frontal, parietal, and hippocampal recordings, all three single-channel configurations supported high-quality scoring (MF1 = 0.86, 0.88, 0.84 respectively; Fig. 4F, left; Table S13), with hippocampal recordings showing a small reduction concentrated on REM F1 (0.78 vs 0.83– 0.86 cortically). In 12 mice from the Oxford sleep-deprivation dataset with simultaneous surface EEG and multi-channel intracortical probes (Fig. 4g; Methods), Nyx supported sleep scoring across the probe channels (median MF1 = 0.81). Performance was substantially more variable across intracortical recordings than surface EEG (Table S14). On average, intracortical scoring fell modestly short of paired surface EEG (paired ΔMF1 = 0.07), consistent with this channel-level heterogeneity. These results suggest that although parietal cortical recordings provide the most consistent performance, Nyx remains adaptable to diverse recording configurations. As a further test of robustness to non-standard telemetric recording conditions, we applied Nyx to a recording from a wild rat (Fig. S7e) and recovered high-quality scoring (MF1 = 0.95).

A more consequential choice seems to be whether an EMG channel is available (Fig. 4f, right; Fig. S7d). The same in-house adult rats were re-scored using a parietal–frontal derivation, in three configurations differing in EMG availability. With a separate EMG channel, the derivation matched the single-channel performance (MF1 = 0.86). Replacing the EMG with an EMG-like^22^ (see Methods) signal derived from EEG and LFP channels produced a small drop concentrated in WAKE (MF1 = 0.81). Nyx thus applies to recordings from our laboratory without dedicated muscle electrodes — a common constraint in research groups where sleep is a secondary variable — provided surrogate electrophysiological signals can reconstruct an EMG-like signal. Removing EMG entirely, with no attempt to compensate for it with an alternative such as accelerometer or EMG-like signals, produced a larger and more variable drop (MF1 = 0.71), driven by a collapse of WAKE F1 (0.47 ± 0.33) and corresponding to a WAKE/REM confusion (Fig. S7d). The pattern identifies EMG as the most consequential of the recording choices tested; electrode placement instead can be varied at limited cost across surface region, depth, and intracortical channels when using Nyx.

Together, these analyses establish that Nyx generalises very well across the three axes that vary most in sleep research: developmental stage, species, and recording configuration. Within recording configurations, changes in electrode placement have a limited cost on accuracy, while EMG availability (or suitable alternatives) emerges as a key factor that affects scoring quality.

### Nyx extends to real-time scoring and closed-loop applications

Since a growing number of applications require scoring sleep as it happens, we developed Nyx-RT, a real-time implementation. This extension builds a model from a baseline sleep recording that encapsulates the wake-sleep threshold, the PCA projection, and the cluster mapping (Fig. 5a, see Methods for details). We use it to score an ongoing recording while dynamically tracking how the decoded state (a single point in PC space) moves from one cluster (e.g. NREM) to another (e.g., REM), for example in Video S1. We revisit this continuous view of sleep-state dynamics, and what it reveals beyond discrete stage labels, in the last section.

**Fig. 5:**
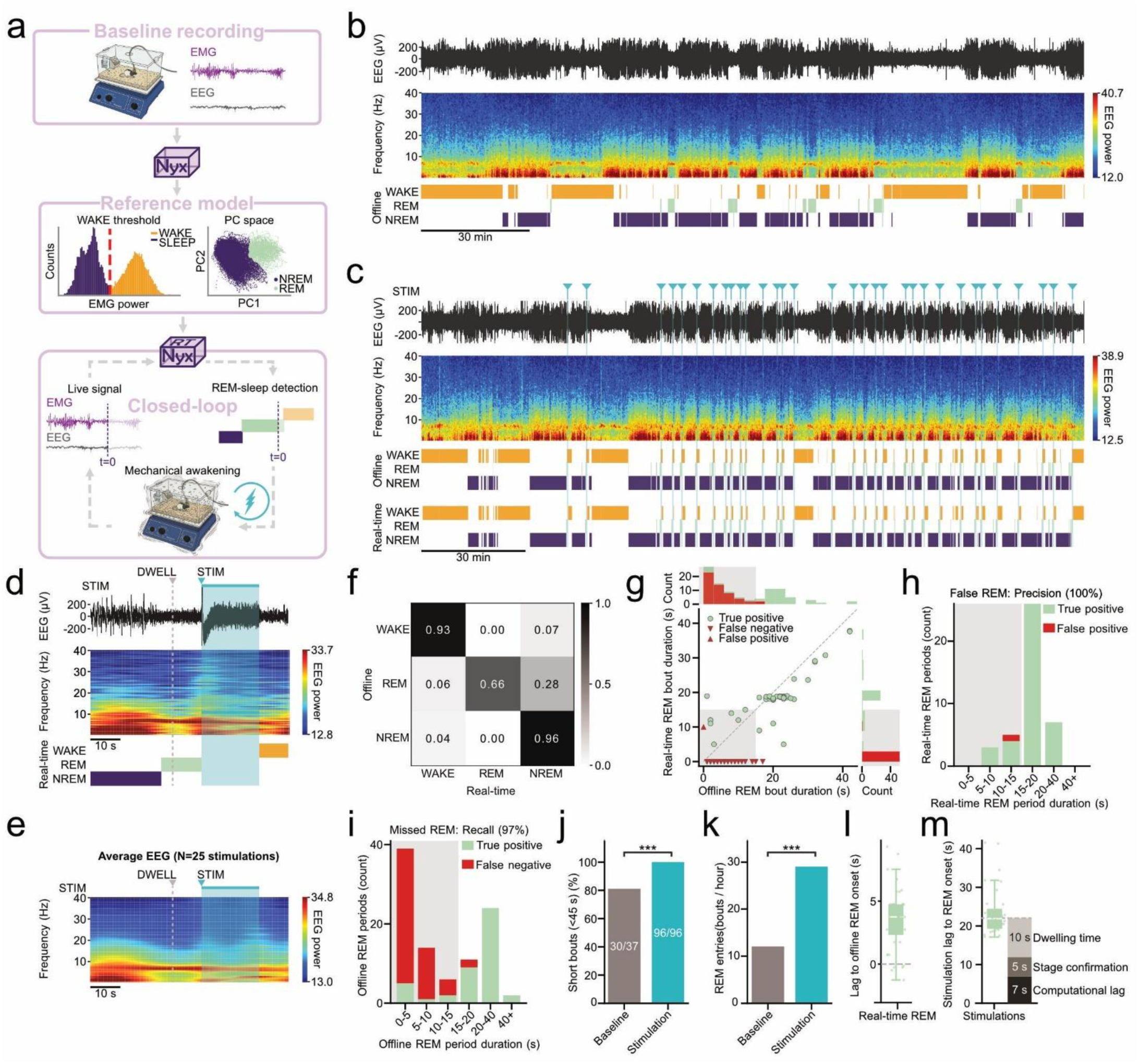
Nyx extends to real-time scoring for closed-loop applications **a,** Schematic of the workflow required to use Nyx for real-time sleep scoring and closed-loop applications. **b,** Nyx scoring of a baseline recording, showing sustained REM bouts in the absence of manipulation. **c,** Scoring of a REM-targeted manipulation recording using offline Nyx (top) and real-time Nyx (bottom), demonstrating that real-time scoring reproduces the offline result while enabling the closed-loop intervention. Shaker activations are marked with blue triangles and shaded areas. **d,** Example scoring from the real-time algorithm around a stimulation. The algorithm requires a 5 s stage confirmation window following REM onset, after which a 10 s dwell period elapses before stimulation delivery. **e,** Average of 25 EEG spectrograms aligned on stimulation onset; 7 stimulations with extended dwell period excluded. Before stimulation, NREM slow-wave activity fades as REM theta oscillations emerge. **f,** Confusion matrix normalised by row, displaying per-class recall. Real-time scoring is compared against offline Nyx scoring used as ground truth. Per-class recall is 93% WAKE / 96% NREM / 66% REM, with the largest errors confined to short REM bouts (see g–i). **g,** Matched real-time vs offline REM bouts duration. A real-time and an offline REM bout were considered matched if their onsets fell within ±10 s of each other. Red triangles correspond to periods without match (upwards: false positive, downwards: false negative). The grey shaded area marks bouts below the stimulation-triggering threshold (< 15 s). Unmatched bouts are almost entirely short (< 15 s), i.e. below the intended detection range. **h-i,** Precision (**h**; N=32) and recall (**i;** N=36) of real-time REM detection. Bouts < 15 s (grey shaded area) were excluded from accuracy metrics as they fall below the minimum duration reachable by the stimulation protocol (5 s persistence + 10 s dwell) and therefore cannot be targets for closed-loop intervention. **j,** Proportion of short REM bouts (< 45 s) in the baseline and stimulation conditions. Short REM bouts are enriched under stimulation (Fisher’s exact test, p=8.28×10^-5^), consistent with premature REM termination. **k,** REM entry frequency in the baseline and stimulation conditions (baseline: 12.0 bouts/h; stimulation: 29.1 bouts/h), more than doubling under stimulation (Poisson means test, p=1.39×10^-6^), consistent with increased REM pressure. **l,** Lag between real-time and offline-scored REM bout onsets, for true positive detections only. Mean ± sem: 3.6 ± 0.4 s (N=40 bouts). The real-time detection lags offline scoring due to the one-sided smoothing kernel **m,** Distribution of latency between offline REM onset and stimulation delivery. The expected latency from the real-time implementation (22 s: 5 s confirmation + 10 s dwell + 7s computation) is shown as a stacked bar on the right for reference. Mean ± sem: 23.6 ± 1.1 s. N = 32 stimulations. The measured latency closely matches the expected value, consistent with the known sources of delay.

A main application of Nyx-RT is that it enables closed-loop manipulations in which a stimulus (e.g., mechanical or optogenetic) is automatically delivered on entry into a target stage^23^. To validate our implementation, we ran a proof-of-concept experiment. First, we recorded a 3-hour baseline sleep session (Fig. 5b). On the following day, we used Nyx-RT to awaken the animal upon REM sleep entry via the activation of an orbital shaker (Fig. 5c,^24^). The manipulation session lasted about 3 hours, similar to the baseline period. This design let us evaluate both how accurately Nyx-RT detects REM sleep in real time and how effectively that detection could translate into REM disruption.

To reduce false-positive stage transitions, we added two safeguards: we confirmed a stage transition only after 5 consecutive seconds in the new stage, and triggered stimulation only after a further, adjustable dwell period (10 s here). These values were chosen conservatively for this proof of concept rather than optimised; shorter persistence and dwell periods would likely suffice, and could yield greater total REM suppression. EEG activity around a single (Fig. 5d) and averaged (Fig. 5e) stimulations showed the shaker switching on following REM entry within the expected latency, and reliably waking the animal. The underlying state change was visible directly in the signal: slow-wave activity, the hallmark of NREM, faded approximately 20 s before stimulation, just as theta oscillations, the hallmark of REM, emerged.

Overall, the closed-loop algorithm shows strong agreement with offline scoring (Fig. 5f, MF1 = 0.88). For closed-loop applications specifically, the critical performance metric is the reliable detection of REM bouts long enough to trigger stimulation. We therefore measured the accuracy to detect REM bouts longer than 15 s (5 s persistence and 10 s dwell; Fig. 5g), where Nyx-RT achieved perfect precision (100%; Fig. 5h) and near-perfect recall (97%; Fig. 5i). This detection performance translated into effective REM disruption: while multiple long REM bouts (> 45 s) could be identified during the baseline recording, none were left during stimulation (Fig. 5j). Concurrently, the frequency of REM entries was significantly higher (Fig. 5k), consistent with increasing homeostatic REM pressure.

To characterise how precisely detection translated into stimulation timing, we quantified the latency between REM sleep onset and stimulation delivery. A computational lag of ∼ 7 s arises from buffering and processing delay (3.5 s), along with an offset between real-time and offline REM entry caused by the one-sided smoothing kernel (3.5 s, Fig. 5l). Combined with the 5 s persistence criterion and the 10 s dwell period, this yields a theoretical latency of 22 s between REM sleep entry and stimulation, consistent with our measured values (Fig. 5m). This latency could be reduced further in two ways. The 7 s computational lag could be partially mitigated through further optimisation of windowing and smoothing parameters. The persistence criterion and dwell period, by contrast, offer more direct control: lowering or abolishing either would substantially cut total lag, though at the cost of increased off-target stimulation risk.

Taken together, these results show that Nyx can be readily adapted for real-time sleep scoring, enabling long-term automated closed-loop sleep manipulations and live tracking of the current decoded brain-state as it moves between cluster boundaries in the PC space.

### Nyx reveals sub-stage and species-specific state structure beyond conventional scoring schemes

Having established that Nyx reliably reproduces conventional sleep scoring, we next asked whether its unsupervised design could reveal structure that conventional scoring schemes do not capture — from finer temporal resolution within established stages to entirely new state definitions in species where no scoring convention exists.

Conventional epoch durations (4 s in rodents, 30 s in humans) reflect manual-scoring practicalities rather than an intrinsic timescale of the sleep–wake signal. Repeating our analysis across a range of epoch lengths in both species, Nyx’s classification agreement and cluster stability remained consistent throughout (Fig. S8, Table S15), indicating that conventional epoch durations can be shortened — recovering finer temporal resolution for transitions and short-lived events — without compromising accuracy or the underlying cluster structure.

Pushing this logic further, we asked whether abandoning epoch-binned representations altogether could reveal sub-epoch structure within established stages, namely N2 and N3. As a proof-of-concept, we applied Nyx to an adult rat recording using a wavelet scalogram backend, yielding a per-sample time–frequency representation that we then reduced with PCA as in the standard pipeline. Restricting the analysis to NREM epochs, three clusters emerged: a delta-dominant cluster (N3), a cluster combining delta with elevated sigma-band (10-16 Hz) activity (N2), and a transitional cluster (TR) which, in PC space, geometrically bridged the NREM block to the WAKE and REM clusters (Fig. 6a). The mean cluster PSDs confirmed these spectral signatures and validated the stage assignments, extending the standard 3-stage scheme to a 5-stage hypnogram with biologically interpretable NREM substages. Examining the finer temporal structure, the smoothed wavelet scalogram separated substages substantially better than the standard short-time Fourier spectrogram, with the sigma-band difference between N2 and N3 reflecting short-lived discrete events rather than sustained power differences (Fig. 6b). Returning to the broadband EEG trace, each of these events was independently captured by a standard spindle detection algorithm, and spindle density was approximately 9 times higher in N2 than in N3 epochs (8.4 vs 0.9 events/min). Nyx’s N2 cluster therefore captures genuine spindle-rich epochs, with its defining sigma-band signature corresponding to discrete, independently detectable events. Regarding the transitional cluster, we refrained from naming it N1, as its mean PSD reflected a heterogeneous mix of transitions that, in our opinion, do not constitute a single defined stage. All in all, this demonstrates that Nyx can resolve NREM substages in rodents, a capability rarely explored to date.

**Fig. 6:**
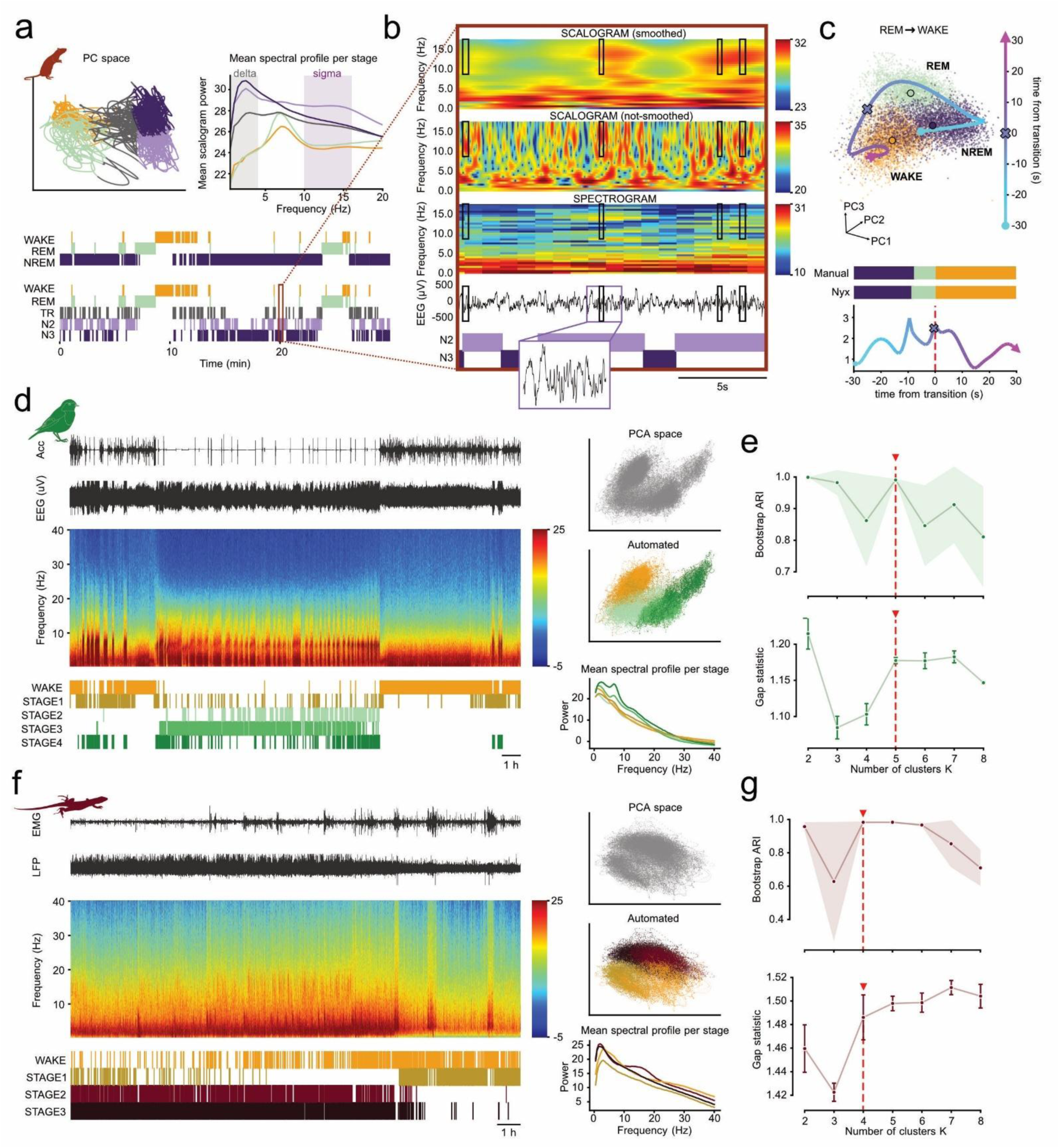
Nyx reveals sub-stage and species-specific state structure beyond conventional scoring schemes **a–b,** NREM substage analysis in an adult rat recording. **a,** Top left: PC space projection of the full recording, with NREM epochs coloured by substage assignment (N2, N3, TR); the TR cluster occupies an intermediate position between the NREM block and the WAKE/REM clusters. Top right: mean power spectral density per stage, showing the delta-dominant N3 signature and the delta plus sigma-band N2 signature. Bottom: 3-stage and 5-stage hypnograms, the latter extending the scheme with N2 and N3 substages plus a transitional cluster (TR). **b,** 20-s illustration of substage scoring. Top to bottom: smoothed wavelet scalogram (used for substage clustering); the same wavelet scalogram unsmoothed, showing short-lived events; standard short-time Fourier spectrogram for comparison (2 s bin); EEG trace. Black rectangles mark detected spindle events. The smoothed wavelet representation resolves discrete short-lived sigma events that are blurred in the spectrogram. Spindle density in the example recording: 8.4 events/min in N2 (n = 61 events) vs 0.9 events/min in N3 (n = 8 events). **c,** A single representative NREM→REM→WAKE transition from one recording in the PC space. Top: epoch cloud in PC1–PC2–PC3 coloured by state (WAKE orange, NREM purple, REM green), with the three state centroids labelled and the ±30 s trajectory overlaid, colour-coded by time from the transition (cyan→magenta, colorbar); “•” marks the trajectory start, “X” the transition (t = 0), and the arrowhead the end. Middle: Manual and Nyx hypnograms over the same window. Bottom: distance from each epoch to its assigned state centroid (Mahalanobis, a.u.) on the same time-from-transition colour scale. The state travels continuously between centroids while the label switches at a single instant. **d–g,** Unsupervised state discovery in non-mammalian recordings. **d,** Jackdaw (EEG + accelerometer)^21^ example traces, spectrogram, hypnogram, PC space coloured by cluster assignment, and mean cluster PSDs for the selected K = 5 partition (Wake + Stages 1–4). **e,** Cluster-number selection criteria: bootstrap ARI (mean ± SD across 190 pairs from 20 × 80% subsamples) and gap statistic (mean ± SE across 50 reference datasets) for K = 2–8. Bootstrap ARI shows bimodal stability peaks at K = 2 and K = 5, with reduced stability at intermediate K; the gap statistic plateaus from K = 5 onwards. Selected K = 5 marked with a dashed line and red triangle. **f,** Tegu lizard example, same layout as (D) with LFP data obtained from an electrode located in the DVR, for the selected K = 4 partition (Wake + Stages 1–3). **g,** Bootstrap ARI shows nested stability with peaks at K = 2 and K = 4 separated by a collapse at K = 3; the gap statistic plateaus from K = 4 onwards. Selected K = 4 marked with a dashed line and red triangle. Cluster labels (Stages 1–N) are generic and deliberately not mapped onto mammalian stage nomenclature. All bootstrap analyses used 20 × 80% subsamples without replacement with fixed per-draw seeds; the gap statistic used B = 50 uniform reference datasets drawn from the feature-space hbounding box. Detailed statistics in Supplementary Table S16.

Because Nyx assigns each epoch a position in PC space rather than only a discrete label, the same output can be used to track how sleep states evolve over time. Fig. 6c shows representative NREM-to-REM-to-WAKE transitions as trajectories through this space, with epochs moving progressively between clusters rather than switching abruptly at a single boundary. A comparable continuous trajectory has recently been reported for NREM-to-REM transitions in brainstem population activity^25^. With Nyx, the position of each epoch within this space also carries information beyond its assigned label: epochs near a cluster centre reflect confidently classified, stable states, while epochs near the periphery sit close to a cluster boundary and reflect states in transition (Fig. S9a-c, Video S2). This distinction can further separate two qualitatively different transition patterns: recordings where epochs settle quickly into one cluster and remain there, reflecting consolidated sleep, versus periods where epochs repeatedly cross back and forth between adjacent clusters before stabilising — a signature consistent with fragmented or unstable sleep rather than a single, well-defined transition (Fig. S9d-f, Video S3). This tracking is not limited to offline analysis: as shown above (Video S1), the same trajectories can be followed live during a recording using Nyx-RT.

To assess whether Nyx can recover state structure in recordings where no a priori scoring scheme applies, we explore two phylogenetically distant species: one jackdaw (Fig. 6d) and one tegu lizard (Fig. 6f). For the jackdaw, this also let us revisit the limitation noted in Figure 4 — where REM-like state separation was constrained by the reference scoring — by letting the clustering run unconstrained. In both cases, the number of clusters was selected empirically using bootstrap stability and gap statistic. To avoid over-interpretation, clusters were labelled generically as WAKE and Stages 1–N rather than mapped onto mammalian stage nomenclature.

For the jackdaw, bootstrap adjusted Rand index (ARI) peaked sharply at K = 2 (0.999) and again at K = 5 (0.991), with intermediate K values markedly less stable (Fig. 6E). This bimodal stability profile is diagnostic of nested cluster structure: the K = 2 partition captures the coarse wake/sleep dichotomy, while K = 5 resolves the finer state architecture within it. The gap statistic supported this finer partition, plateauing from K = 5 onwards. We therefore selected K = 5, yielding WAKE and Stages 1–4 with distinct spectral signatures in their mean cluster PSDs. For the lizard, bootstrap ARI showed a nested stability pattern: K = 2 was highly stable (0.957; the trivial wake/sleep split), K = 3 dropped and stability recovered fully at K = 4 (0.983), persisting through K = 6 before degrading again (Fig. 6g). The gap statistic plateaued from K = 4 onwards, with subsequent increments comparable to their standard errors. Both criteria therefore converged on K = 4 as the finest stable partition beyond the wake/sleep dichotomy, yielding WAKE and Stages 1–3 with distinct spectral signatures in their mean cluster PSDs. We emphasise that these are illustrative single-recording demonstrations of the potential of Nyx rather than species-level claims.

Together, these exploratory analyses illustrate that Nyx’s data-driven architecture supports state discovery at two complementary scales: refining within established stages when sub-epoch temporal resolution is available and recovering principled state partitions in recordings where no a priori scoring convention applies. In both regimes, the pipeline’s outputs remain anchored to interpretable spectral signatures and convergent stability criteria, providing a framework for extending sleep research beyond the species and conventions for which it was historically developed.

## Discussion

Sleep scoring algorithms have historically been built and validated within narrow contexts — a single species, a fixed age range (adults), one recording modality. As a result, the range of data they could score was limited in its nature. Generalisation and comparison across contexts remained out of reach, and sleep structures outside those original contexts were likely to go undiscovered. Nyx was designed to close this gap, providing a framework that is reproducible, generalisable, efficient, and explainable. Unlike supervised classifiers, which inherit the conventions of their reference labels, Nyx takes an unsupervised approach. It uses dimensionality reduction to directly reveal the underlying state geometry of each recording: epochs are projected into a low-dimensional PCA space, where they cluster according to the structure present in the data itself, free of pre-defined conventions. Because the geometrical distribution of the labels is directly visible, users can inspect them and re-cluster or adjust boundaries through the graphic user interface (GUI), keeping labelling transparent, inspectable and adaptable.

Nyx was thoroughly validated across 19 datasets spanning species, lifespan, conditions and populations. Not only does it perform on par with algorithms built specifically for a recording setup in a given species (e.g., Somnotate, YASA), but it also generalises across datasets independently of epoch length. Previous pioneering attempts at unsupervised sleep scoring were limited in both scope and accessibility, preventing their widespread use. FASTER and FASTER2 were confined to mice and AISleep to humans, and all relied on predefined frequency features. Nyx addresses these limitations at once, learning its spectral representation directly from the data, matching human inter-scorer reliability under all tested conditions, while remaining usable by researchers regardless of programming background. In rodents, agreement with manual scoring is preserved across pharmacological, genetic, and experimental manipulations, with residual errors concentrated at state transitions where inter-scorer disagreement is itself highest. In both healthy and clinical human recordings, performance is likewise high, with N1 the main exception — the stage on which expert scorers themselves rarely converge (∼15–20%), placing an inherent ceiling on what any classifier can achieve. Users should be aware that Nyx (like many automatic scorers) tends to overestimate REM-to-NREM ratio and mislabel microarousals. However, we explicitly designed workarounds addressing these limitations: post-hoc rules can be applied as filters for downstream analyses, while epoch length and smoothing are tuneable, allowing shorter windows to recover microarousal sensitivity when needed.

While Nyx performance holds up remarkably well under conditions that typically expose the limits of automated scoring, benchmarking against manual scorers should be interpreted with caution. Indeed, manual scoring is itself a variable, subjective process built on arbitrary conventions, and therefore not a neutral ground truth — supervised classifiers are effectively evaluated against the very conventions they were trained on, inflating their apparent accuracy while underestimating that of unsupervised ones. Furthermore, rules such as the prohibition of direct WAKE-to-REM transitions can bias scoring in rare yet clinically meaningful situations such as REM sleep behaviour disorder, an early biomarker of Parkinson’s disease. One solution could be to move away altogether from discrete stages in favour of continuous oscillatory measures^26^ — yet this may introduce reproducibility challenges. Nyx offers a middle ground: it preserves comparability while remaining flexible enough to capture what rigid conventions miss. This flexibility translates into analyses previously out of reach. Unconstrained by fixed scoring conventions, it allows deviations from the canonical architecture, enabling comparative analyses across developmental stages and species that supervised pipelines cannot support. Operating directly on signal structure, Nyx also extends naturally to the discovery of finer-grained substages and novel sleep patterns, demonstrated here as proofs of concept and amenable to systematic extension.

Nyx’s real-time implementation (Nyx-RT) extends this further, letting us track the decoded brain-state live as it moves through PC space, thus offering a direct window into transition dynamics, while also enabling real-time scoring for closed-loop experiments. This live view points to a more fundamental shift in how sleep is conceptualised in our base framework: as a continuous dynamical process rather than a sequence of discrete stages^27^. The low-dimensional embeddings produced by Nyx provide a natural substrate for analyses that treat sleep as a trajectory through state space, rather than only as a series of categorical epoch labels^25,28^ — enabling future in-depth analyses of sleep transitions and sleep architecture consolidation.

As an open-source project Nyx is designed to evolve with the community that uses it. Its modular interface and flexibility support adoption across diverse experimental and clinical contexts. As the very definition of sleep is increasingly called into question, Nyx is built to adapt and keep pace with a field whose scope is bound to expand beyond rigid conventions. Beyond a practical tool for sleep staging, it offers a framework for revealing how sleep states are organised across brains, species and recording modalities.

## Methods

### Datasets

To evaluate Nyx across a broad range of species, ages, recording systems, and physiological and pathological conditions, we considered 19 datasets comprising 662 recordings from 380 animals/subjects, spanning rodents (10 datasets, 374 recordings), humans (7 datasets, 259 recordings), jackdaws (1 dataset, 27 recordings), and 1 lizard and 1 wild rat (1 dataset, 2 recording; Table S1). These included in-house rodent recordings acquired with tethered and telemetry systems (see “In-house recordings” alongside publicly available (or provided by collaborators) rodent, human, and avian datasets. For the largest human cohorts (MESA^8,9^, CCSHS^9,10^, CHAT^9,11^), representative subsamples were drawn by proportional stratified random sampling; the procedure is detailed in Methods (“Dataset subsampling”). Manual scores accompanying each dataset served as ground truth, except where stated.

#### Dataset subsampling

For the largest human cohorts, representative subsamples were drawn by proportional stratified random sampling. In each case, strata were defined by the cross-classification of the demographic variables specified below; within each non-empty stratum, the number of recordings to draw was set proportional to that stratum’s share of the eligible pool, with a minimum of one recording per non-empty stratum. Recordings were then drawn uniformly at random, without replacement, using a fixed random seed for reproducibility. This procedure preserves the demographic composition of the source cohort in the subsample.

From the 2237 polysomnography recordings available in the MESA^29,30^ dataset, we selected 60 recordings using this procedure. Strata were defined by gender (female, male), self-reported race/ethnicity (White, Chinese American, Black/African-American, Hispanic), and age group in 10-year intervals (50-59, 60-69, 70-79, 80-89, 90+), yielding 39 non-empty strata.

For the older-adult subset (MESA, ≥70 years), 32 previously analysed recordings were excluded from the 1064 participants aged 70 years or older, leaving 1032 eligible recordings, from which we selected 25. Strata were defined by gender, self-reported race/ethnicity (as above), and age group (70-79, 80–89, 90+), yielding 23 non-empty strata. 4 of the drawn recordings were not available in the public EDF release and were each replaced by re-drawing one recording from the same stratum.

For adolescents, we selected 20 recordings from the 517 polysomnography recordings in the CCSHS^30,31^ dataset. Because the age range spanned less than four years, strata were defined by gender and race only (White, Black/African-American, Other), yielding 6 non-empty strata.

For children, we selected 20 recordings from the 453 polysomnography recordings in the CHAT^30,32^ dataset. Because the age range spanned less than five years, strata were defined by gender and race only (American Indian / Native Alaskan, Asian, Native Hawaiian / Other Pacific Islander, Black / African American, White / Caucasian, Multiracial, Other, Unknown), yielding 10 non-empty strata.

#### Channel selection

For the main validation, one EEG and one EMG channel per recording were used. For rodent datasets, the EEG channel was the parietal electrode where identified; the only EEG channel in single-channel datasets; or, where channels were labelled only generically (e.g. EEG1/EEG2) without confirmed positions, a consistent fixed channel of unconfirmed anatomical location. For human datasets, the C4-M1 (or C4-TP9) derivation was used, with C3-M2 as a fallback. For jackdaw recordings, we selected the same anatomically corresponding left-and right-hemisphere electrode pair across all birds.

#### Ground-truth scoring preprocessing

Manual scores were used as ground truth for evaluating the algorithm. For rodents, micro-arousals, sleep movements, and transitional or artifact-containing epochs were merged into the corresponding WAKE, NREM, or REM state. Because the algorithm scores at a fixed epoch length under its standard parameters (1 s for rodents, 15 s for humans), the ground truth scoring was resampled to the algorithm’s epoch length before comparison, that is each manually scored epoch was subdivided into the corresponding number of shorter epochs.

### In-house recordings

#### Animals

This study used recording data from 10 male Long-Evans rats (*Rattus norvegicus*), ranging in age from postnatal day 8 (P8) to 6 months and weighing no less than 25 g at the youngest recorded age. All animals were housed in groups of 3-6 animals in transparent Plexiglas cages (46 × 40 × 40 cm) in a temperature-and humidity-controlled vivarium, on a 12-h light/12-h dark cycle (lights on at 7 a.m.). Following surgery, post-weaning rats were housed individually when pre-weaning pups (before P21) remained housed with their mother and littermates. All rats were maintained on this schedule throughout the study, with sleep recordings conducted during the light phase. Food and water were available *ad libitum* at all times. All experiments were performed in accordance with the Norwegian Animal Welfare Act and the European Convention for the Protection of Vertebrate Animals used for Experimental and Other Scientific Purposes.

#### Surgical procedures and electrode implantation

All procedures were performed in accordance with institutional guidelines and relevant regulatory licenses, in male Long-Evans rats from postnatal day 8 to adulthood. Animals were anaesthetised with isoflurane (4% induction, 1.5% maintenance) in a stereotaxic frame (World Precision Instruments, Hertfordshire, UK), and received subcutaneous meloxicam (Metacam, 2 mg/kg) and buprenorphine (Temgesic, 0.5–1.0 mg/kg) for perioperative analgesia. Topical lidocaine (Xylocaine, 2%) and chlorhexidine antiseptic (Pyrisept) were applied to the surgical site. Core temperature, heart rate, and blood oxygenation were monitored continuously via a MouseStat system (Kent Scientific, CT, USA) with a feedback-controlled heating pad, and anaesthetic depth was assessed by respiratory rate and hind-paw withdrawal reflex.

For tethered implants, all three tetrodes were implanted into the CA1 subfield of the dorsal hippocampus, while two stainless steel wires were placed epidurally over the contralateral prefrontal (PFC) and parietal (PAR) cortices for electrocorticographic (ECoG) recording, and the remaining two were bilaterally inserted into the dorsal neck muscles for EMG recording. A jeweller’s screw connected to the drive ground pin served as the ground electrode, and the assembly was secured with dental cement (Palaxpress; Kulzer GmbH, Germany). For wireless implants, the transmitter body was placed subcutaneously via a midline dorsal incision, with EEG leads affixed within bilateral parietal craniotomy holes and encapsulated in dental cement, and EMG leads sutured bilaterally into the nuchal musculature. All incisions were closed with non-absorbable sutures and topical antiseptic was applied. Postoperative monitoring continued for seven days, with analgesic coverage maintained for the first 48 hours.

#### Electrophysiological recordings

Electrophysiological signals were acquired using a combination of wireless telemetry (PhysioTel HD-X02; Data Sciences International [DSI], MN, USA) and tethered *in vivo* electrophysiology recording systems (Axona Ltd., St. Albans, UK). Subsets of these recordings were selected for inclusion in the present study. Specifically, telemetric data were obtained from three adult rats (seven recordings total; 2, 3, and 2 recordings per animal, respectively), while tethered data were obtained from six rats (10 recordings total; comprising six recordings during developmental stages and four from adulthood). For wireless recordings, the mouse-specific HD-X02 transmitter was deployed in rat pups due to its small size and light weight. This transmitter featured a biopotential channel bandwidth of 0.5-80 Hz and a nominal hardware sampling rate of 300 Hz. Telemetric data were acquired and digitized at 1kHz using Ponemah software (DSI). Because this transmitter is optimized for smaller rodent species, the telemetry signal was prone to data dropout during periods of vigorous locomotion; consequently, we restricted our telemetry dataset to seven high-quality recordings (approximately 7 hours each) with minimal signal loss, during which the animals were predominantly sleeping. Tethered recordings were acquired using a DacqUSB system (Axona Ltd., St. Albans, UK) at a raw, broadband sampling rate of 48 kHz. These continuous recordings ranged in duration from 15 minutes to 1 hour, spanning three distinct developmental stages and adulthood (Supplementary Table 1).

All raw electrophysiological data were preprocessed using the SpikeInterface ^27^ (version 0.101.2 for DSI telemetry data; version 0.102.3 for Axona tethered data). Axona recordings were first converted from their native broadband format to the Neurodata Without Borders ^28^ (NWB 2.0) standard using a custom Python script. For both systems, signals were bandpass filtered (0.01-90 Hz) and downsampled to 200 Hz. To minimize movement artifacts in the tethered (Axona) recordings, the electroencephalogram (EEG) channel used for sleep scoring was locally referenced as parietal-minus-prefrontal (PAR-PFC). For telemetric (DSI) recordings, periods of signal dropout were automatically identified and zero-filled using the blank-saturation routine in SpikeInterface (absolute threshold = 1000, bidirectional, fill value = 0); these blanked segments were preserved in the continuous signal and subsequently flagged for downstream handling during sleep scoring.

Manual sleep scoring was performed by visualizing the preprocessed EEG and EMG channels simultaneously with their corresponding power spectrograms using a custom graphical user interface built on the ephyviewer framework (https://ephyviewer.readthedocs.io/en/latest/overview.html). Three distinct states were classified according to standard criteria: active wakefulness (WAKE; characterized by high-amplitude EMG power), non-rapid eye movement sleep (NREM; characterized by low EMG power accompanied by prominent slow delta oscillations 0.5-4 Hz), and rapid eye movement sleep (REM; characterized by low EMG power accompanied by sustained, regular theta activity 5-9 Hz). For recordings scored by multiple investigators, a majority-consensus hypnogram was established as the ground truth, wherein each epoch was assigned the state selected by the majority of scorers; epochs resulting in a tie were classified as ‘DISAGREE’ and excluded from subsequent analysis. The in-house telemetric (DSI) recordings were manually scored by five investigators of varying expertise levels. In-house tethered (Axona) recordings were scored by a single expert investigator using identical criteria, which were adjusted appropriately for early developmental stages during which mature hippocampal theta rhythms had not yet fully emerged.

### Sleep scoring algorithm

We developed Nyx, a semi-automated sleep scoring framework requiring only a single EEG channel and an EMG channel (or an equivalent motor-activity signal, such as accelerometer or video pose-estimation data) as input. Nyx applies PCA for dimensionality reduction of the feature space and then supports flexible clustering, with a choice of algorithms (K-means, Gaussian mixture models, HDBSCAN; see Supplementary Information for details), to infer the distributions of the different sleep stages. The overall workflow is illustrated in Fig. 1d.

#### Spectral feature extraction

Raw EMG and EEG signals were processed independently to compute short-time Fourier transform (STFT) power spectral density (PSD) estimates. For each signal, a spectrogram was computed using overlapping Hanning windows (50% overlap) with constant detrending and density scaling, retaining the frequency band most informative for each signal type: 30-100 Hz for EMG, to isolate muscle activity while excluding low-frequency movement and EEG contamination^14,33,34^, and 0.5-40 Hz for EEG, to exclude line noise (50/60 Hz) and low-frequency movement artifacts^3^ (rodent values; see Supplementary Methods for species-specific parameters). The broader preprocessing band (0.01–90 Hz) was retained upstream to preserve other frequency bands for analyses beyond the present study; the 0.5–40 Hz range was used specifically for sleep-stage feature extraction.

Window length and smoothing were set finer than the temporal resolution of sleep-stage transitions, and chosen so that manual scores could be divided into unit epochs without interpolation: 1 s for rodents (accommodating the 4, 5, and 10 s manual epochs across datasets) and 15 s for humans (half the standard 30 s epoch, for finer resolution). These parameters are adjustable; the effect of epoch length was evaluated on one mouse dataset (see “Epoch-length sensitivity analysis”, Supplementary methods).

Spectral power was converted to decibels (10·log₁₀) and normalized to remove inter-recording amplitude variation. For human recordings, each frequency bin was z-scored independently across time to enhance separation between the larger number of stages; for rodent recordings, the global mean spectrum was subtracted across time, as the rodent stages were sufficiently distinct that z-scoring was unnecessary. The two normalization strategies yielded equivalent clustering results.

#### EMG-based arousal state classification

We used EMG power to separate recordings into broad WAKE and SLEEP periods. A band-limited EMG power trace was obtained by summing the normalised log-power across the EMG frequency band at each time point:

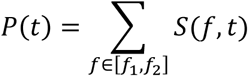

where [*f*_1_, *f*_2_] is the EMG frequency band and *S*(*f*, *t*) is the normalized log-power spectrogram. *P*(*t*) was subsequently smoothed with a uniform moving-average filter and min-max scaled to [0,1] within each recording. In rodent recordings, which typically contain long, uninterrupted wake and sleep bouts and a pronounced bimodal EMG power distribution, an amplitude threshold was applied to the trace to generate an initial binary WAKE/SLEEP hypnogram, with an additional lower bound to identify signal-absent artifact epochs (NOSIGNAL). Throughout the pipeline, NOSIGNAL aggregates epochs flagged as signal-absent at any stage (epochs below the EMG lower bound, and feature outliers identified during clustering) which are retained as a single label until the final hypnogram. The threshold was set at the trough of the bimodal EMG power distribution and could be manually adjusted. A minimum-duration constraint (default 4 s) was applied to suppress spurious brief transitions.

This step is optional. For short or ambiguous recordings, all epochs can instead be passed directly to the EEG clustering step with a uniform SLEEP label, and wakefulness is then recovered during clustering. This applies to human recordings, short rodent recordings, and recordings lacking a clear bimodal distribution (typically due to poor EMG quality or electrode malfunction): in these cases, the EMG power trace was still computed, but used as an additional feature in the EEG clustering step rather than for thresholding.

#### Dimensionality reduction and feature construction

EEG spectrograms were dimensionality-reduced using principal component analysis (PCA), computed on epochs classified as SLEEP in the preceding step; all subsequent feature construction and clustering operated on this SLEEP subset. Restricting the analysis to sleep epochs focuses the decomposition on within-sleep spectral variation (i.e. the difference between NREM and REM) rather than the larger WAKE-SLEEP contrast, which is particularly pronounced in long rodent recordings. The number of components retained was a user-defined parameter (default 5-6), chosen on the basis of explained variance and inspection of the frequency-domain component weights. A feature matrix was then constructed per epoch by combining the selected principal component scores with EMG power where applicable (see “EMG-based arousal state classification”), the latter interpolated onto the EEG spectrogram time grid. Features were standardised to zero mean and unit variance; epochs with any feature value exceeding an adjustable z-score threshold (default 6 standard deviations) were additionally labelled NOSIGNAL and excluded from clustering.

#### Flexible clustering and stage assignment

Sleep-stage classification was performed by clustering the PCA (and EMG, where applicable) feature matrix. Three algorithms were supported and could be selected per recording: K-means, Gaussian mixture models (GMM), and HDBSCAN (see Supplementary Methods). The number of clusters was a user-defined parameter, typically set slightly above the number of expected sleep stages to allow fine-grained initial clustering before merging. Following clustering, the experimenter inspected (i) the frequency profiles of the PCA loadings, (ii) 2D scatter plots of the first two principal components (PC1 vs PC2), with points coloured by cluster and, separately, by higher-component scores (e.g. PC3, PC4) to visualise structure beyond the plotted plane, (iii) per-cluster mean ± 95% CI power spectral density profiles, and (iv) cluster-level EMG statistics, and assigned each cluster to a sleep-stage label (e.g. WAKE, NREM, REM); clusters assigned the same label were thereby merged. This semi-automated curation step is analogous to manual cluster curation in spike sorting: the algorithm proposes a partition, and the expert confirms or adjusts it on the basis of interpretable features. Accuracy and confidence nonetheless improve with practice, and we recommend a supervised familiarisation phase on recordings with available ground-truth labels before independent scoring. Cluster-to-stage mappings were saved to the recording’s parameter file to enable exact reruns without further user input.

#### Human specific hierarchical scoring

Human recordings commonly require discrimination of five sleep stages (WAKE, N1, N2, N3, REM) whose spectral signatures overlap more than those in rodents due to the sub-classification of NREM stages. To improve stage separation, the human pipeline applies three sequential clustering steps, each operating on epochs belonging to a single broad category identified in the previous step. Unlike the single-pass rodent pipeline, in which PCA is fit once on the SLEEP epochs, the human pipeline re-fits PCA within each branch so that the decomposition is tuned to the spectral variation relevant at that level.

**Step 1: Global WAKE/SLEEP refinement.** PCA is run on all epochs, and the resulting PCA + EMG features are clustered (typically 2-3 clusters) to determine the binary WAKE/SLEEP boundary.

**Step 2: WAKE sub-classification.** PCA is recomputed on epochs labelled WAKE in Step 1. Clustering of these epochs separates consolidated WAKE (predominant high frequency EEG, including 8-12 Hz alpha oscillations) from N1 (predominantly low-amplitude 4-7 Hz), the transitional stage that in humans occurs near sleep onset.

**Step 3: SLEEP sub-classification.** PCA is recomputed on the SLEEP epochs from Step 1, and clustering (typically 4-5 clusters) separates N2 (sleep spindles (12–15 Hz) and K-complexes), N3 (high-amplitude slow-wave activity, <4 Hz), and REM (low-amplitude mixed-frequency EEG with muscle atonia). As in the rodent pipeline, clusters assigned to the same label are merged; in addition, for human scoring a principal-component threshold can optionally split a single cluster into two sub-stages in a post-clustering refinement step.

The final five-stage hypnogram was assembled hierarchically from the three clustering outputs: Step 1 defines the WAKE/SLEEP skeleton, which is then subdivided into WAKE and N1 by Step 2 and into N2, N3, and REM by Step 3.

#### Birds specific scoring setup

Because no EMG was acquired in the jackdaw dataset^21^, one of the single accelerometer channels was used in place of EMG as the movement-related input to the EMG-based arousal-state classifier described above. Nyx was applied independently to the left and right hemisphere channels, yielding two per-hemisphere hypnograms per recording. A combined bilateral hypnogram was then derived from the two per-hemisphere hypnograms following the rule hierarchy of van Hasselt et al. (2025): an epoch was labelled WAKE if either hemisphere scored WAKE; otherwise NREM if either hemisphere scored NREM; otherwise, REM if both hemispheres agreed on REM. This precedence reflects the asymmetric organization of avian sleep, in which WAKE and NREM episodes can occur unihemispherically while REM is typically bilateral and brief.

#### Real-time implementation

For validating the real-time sleep scoring application, we implanted a Long Evans rat with two EMG and three ECoG electrodes. A reference model was built from a baseline 3 h sleep session recorded on day D (animal age: P34). First, the session was scored with the same spectrogram and PCA pipeline approach used for conventional offline scoring. Then, several components of the scoring pipeline were frozen to serve as a reference for real-time scoring: the PCA basis, the min-max and standardisation scalers, the baseline PC projections with their REM/NREM labels, and the EMG wake threshold. The animal then underwent REM sleep manipulation sessions on the afternoons of days D and D+1, and a NREM sleep manipulation session on the morning of day D+2.

In contrast to offline scoring, the real-time system relied on the frozen reference model. A single EMG and a single ECoG from the parietal cortex (selected based on best SNR) were recorded at 20 kHz using Intan RHX and streamed towards a Python program using the built-in TCP feature. The signal was downsampled to 500 Hz and bins were classified every second. Prior to classification, EMG band power and EEG spectral features were smoothed using a Gaussian kernel (σ = 4 s). In offline scoring this kernel is applied symmetrically, weighting past and future bins equally. In real-time, only past bins are available; the kernel is therefore one-sided, introducing a temporal prediction lag of approximately 6 s. Following smoothing, power was first computed within 4 s wide EMG bins (75% overlap) and compared against the frozen EMG wake threshold from the baseline session. If the animal was labelled as asleep, the smoothed EEG features were projected into the pre-computed PC space and assigned REM or NREM by k-nearest-neighbour vote (k = 5) against the pre-labelled baseline cloud, rather than by re-clustering.

Because classification relies only on past data, stage transitions are inherently less reliable than in offline scoring. To circumvent this limitation, we added a persistence criterion: when a change of state was detected, it was first flagged as ‘uncertain’. Only after it had persisted for at least 5 s was it committed and confirmed. If the state change reverted during these 5 s, the tentative state change was annulled. This introduced a detection latency with no offline counterpart. Importantly, any transient state shorter than 5 s cannot be detected with this criterion.

The real-time scoring algorithm was used to drive a closed-loop stimulation after a certain duration, the ‘dwelling time’, had been spent in a specific stage. If during the dwelling period, the algorithm detected a potential change of stage — even unconfirmed — the timer would reset to its initial value. During this experiment, dwelling time was set at 10 s, and the closed-loop stimulation involved activating an orbital shaker, on which the cage was positioned, to awaken the animal via mechanical stimulation. The shaker was activated for 20 s, ramping up from 0 to 160 RPM during the first 10 s, after which it reached a constant RPM.

### Implementation and availability

#### Validation and analysis

##### Performance evaluation

Automatically scored hypnograms were compared with manual ground-truth labels using epoch-by-epoch agreement metrics: overall accuracy, Cohen’s κ, macro F1 score, and per-class precision, recall, and F1 score (see Supplementary Methods). Agreement was computed at the algorithm’s scoring resolution (1 s for rodents, 15 s for humans), to which the manual scores had been resampled (see Comparison to manual scoring). Epochs labelled DISAGREE or NOSIGNAL were excluded from all agreement metrics. Confusion matrices were expressed as percentages. For human recordings, performance was evaluated at three levels of granularity^35^: five-stage (WAKE, N1, N2, N3, REM), four-stage (merging N1 and N2), and three-stage (WAKE, NREM, REM, matching the rodent staging).

##### Sleep-variables comparison

To assess whether the automatic scorer (Nyx) retrieves a physiologically meaningful characterization of each recording, we derived a set of standard sleep variables from the manual and automated hypnograms and compared the two annotations on a per-recording basis. The following variables were computed:

- Total sleep time (TST), expressed as the percentage of the recording occupied by REM and NREM epochs.
- REM/NREM ratio, defined as the ratio of REM time to NREM time within each recording.
- Microarousals, defined as brief WAKE bouts of less than 20 s embedded within sleep, expressed as a rate per hour of total recording.
- Theta (5-9 Hz) peak frequency during REM, and computed from the EEG spectrogram in epochs scored as REM.
- Delta (1-4 Hz) integrated power during NREM, computed from the EEG spectrogram in epochs scored as NREM.

Spectral features were obtained as described above (see Spectral feature extraction) and averaged across the recording within each stage-specific epoch. Integrated power was computed within the band of interest and taking the within-band frequency of maximum power as the peak frequency. Power values are reported in dB. For each variable, manual and automated values were computed from the same underlying EEG signal, differing only in the hypnogram used to select epochs.

For human recordings, an analogous set of variables was derived from the manual and automated hypnograms:

- Total sleep time (TST), expressed as the percentage of the recording occupied by REM and NREMs epochs.
- REM/NREM ratio, defined as the ratio of REM time to NREM time within each recording.
- Sleep onset latency, defined as the time from recording onset to the first SLEEP epoch, in minutes.
- Wake after sleep onset (WASO), defined as the total WAKE time after sleep onset, in minutes.
- REM latency, defined as the time from sleep onset to the first REM epoch, in minutes.
- Sigma (12-16 Hz) integrated power during N2, computed from the EEG spectrogram in epochs scored as N2.
- Delta (0.5-4 Hz) integrated power during N3, computed from the EEG spectrogram in epochs scored as N3.
- Theta (5-9 Hz) integrated power during REM, computed from the EEG spectrogram in epochs scored as REM.

Spectral features were computed as for rodents and reported in dB. Human sleep variable comparisons were restricted to the primary analysis set, excluding recordings flagged as *disagree* (see Performance evaluation).

##### Recording configuration analysis

To assess whether Nyx’s performance depends on the specific electrophysiological signal used as input, we ran a set of within-recording comparisons across cortical regions, LFP depths, re-referencing schemes, and EMG configurations. Two datasets contributed. The Oxford mouse benchmark (test and sleep-deprivation sub-datasets) provided 12 mice implanted with EEG screws over frontal and parietal cortex, and 6 mice implanted with a 16-channel laminar probe (NeuroNexus A1x16-3mm-100-703, 100 µm spacing) from which we retained the same 5 LFP channels per animal, spanning the cortical layers. The in-house tethered rat recordings (n = 2 animals, 4 recordings) provided simultaneous prefrontal EEG, parietal EEG, and hippocampal CA1 LFP. For each comparison, every recording was scored independently with each derivation used as the single-channel input to the standard Nyx pipeline, and the resulting hypnograms were compared against the manual scoring.

###### Frontal vs parietal derivation

To test whether automatic-vs-manual scoring agreement differed between frontal and parietal derivations, we used a single paired design pooling all the animals. For the in-house rat recordings, we averaged that animal’s sessions to a single animal-level mean to avoid pseudoreplication. This yielded n = 14 paired units (12 mice + 2 rats). For each metric (accuracy, Cohen’s κ, macro-F1, and per-stage F1 for WAKE, NREM, REM) we compared frontal and parietal values with a two-sided Wilcoxon signed-rank test. Effect sizes are reported as the matched-pairs rank-biserial correlation (positive values indicate frontal > parietal). To control the family-wise error rate across the six metrics, *p*-values were adjusted with the Holm–Bonferroni procedure; adjusted *p* < 0.05 was considered significant.

###### Hippocampal-derivation comparison

In the two in-house adult rats (n = 4 recordings) we compared automatic-vs-manual sleep-scoring agreement across three single intracranial derivations: frontal EEG (prefrontal-cortex ECoG), parietal EEG (parietal-cortex ECoG) and hippocampal LFP (CA1). For each recording we computed accuracy, Cohen’s κ, macro-averaged F1, and macro-averaged precision and recall against the manual scoring, where macro metrics weight the three vigilance states (WAKE, NREM, REM) equally. Because only four recordings from two animals were available, this comparison is descriptive and no inferential statistics were computed across derivations.

###### EMG configuration and bipolar re-referencing

To assess Nyx’s sensitivity to the EMG input, we compared, in the same 2 rats, three configurations of the bipolar parietal–frontal derivation: paired with the recorded EMG, paired with a surrogate EMG computed from the intracranial signals (EMG-like), and without any EMG channel (no EMG). The single parietal ECoG (PAR) was included as reference to also assess the effect of bipolar re-referencing. The EMG-like signal was derived from the brain signals following Schomburg et al. 2014^22^. LFP and ECoG signals were loaded at 1 kHz and band-pass filtered to 275–600 Hz (4th-order Butterworth, zero-phase). The 12 LFP and 2 ECoG channels were pooled (14 channels total), and the signal was divided into consecutive non-overlapping 100 ms windows. For each window, the Pearson correlation matrix across the 14 channels was computed and the mean of its off-diagonal (lower-triangular) entries taken, yielding a single EMG score per window that quantifies inter-channel coupling in the high-frequency band. The resulting time series was min–max normalised to [0, 1] and used in place of the recorded EMG power for the wake/sleep classification step. Macro-F1 is shown per configuration as for the single-derivation comparison (boxplots with one line per recording on a common y-axis), alongside the row-normalised confusion matrices (manual vs Nyx WAKE/NREM/REM) for each configuration. As above, this comparison is descriptive given the small number of recordings.

###### Intracortical vs surface-EEG scoring performance

For the six sleep-deprivation recordings scored from both intracortical and surface signals (6 mice), we compared automatic-vs-manual agreement (macro F1) across the two modalities. Each recording contributed five intracortical channels and two surface EEG derivations (frontal and parietal). We characterised the two modalities along two axes: the level of performance (per-recording median F1 across channels) and its variability (per-recording standard deviation of F1 across channels. Two complementary cluster bootstraps were run, each with 10,000 iterations and a fixed seed (NumPy default_rng(42)), resampling recordings with replacement as the clustering unit. A marginal bootstrap pooled all channel-level F1 values within each modality and compared the pooled median (ΔM) and pooled SD (ΔSD) between modalities; a paired bootstrap computed the per-recording modality difference and bootstrapped its mean across recordings. Effects were considered significant when the 95th-percentile confidence interval excluded zero; two-sided p-values were derived from the bootstrap tail fraction.

##### Rodent substages sleep scoring

One cortical EEG (ECoG) recording from an in-house adult rat was used for the example analysis in Fig. 6a-b. The recording was downsampled to 200 Hz and analysed over a 30-min window (0–1800 s). Two complementary time–frequency representations were computed from the same EEG trace. A short-time Fourier spectrogram was computed with scipy.signal.spectrogram (2-s Hann windows, 50% overlap, power spectral density scaling, constant detrend), converted to decibels and mean-normalised, yielding ∼1-s time bins over 0.5–40 Hz. A wavelet scalogram was computed using a complex Morlet transform (ephyviewer implementation; central frequency f0 = 1 Hz) over 0.5–40 Hz at 0.5-Hz resolution, at the native sampling resolution. Each wavelet’s coefficients were amplitude-weighted by the inverse of its scale (scale⁻¹). Because scale is inversely proportional to frequency, this weighting is equivalent to scaling power in proportion to frequency, which compensates for the lower amplitudes of high-frequency wavelets. The resulting scalogram was converted to decibels. For the staging analyses both representations were Gaussian-smoothed in time (σ = 1 s); for the single-bout visualisation (Panel b) the scalogram was recomputed without temporal smoothing to preserve the native time resolution of discrete events. NREM epochs were defined from the main Nyx scoring run on this recording (see Section “Sleep scoring algorithm”) with the smoothed scalogram backend and used as the input mask for the substage analysis. PCA was fit on the smoothed scalogram restricted to NREM epochs, and a Gaussian mixture model (3 components) was fit on the principal components individually explaining >5% of variance (the first three), after standardising features and removing outliers (|z| > 6). The three resulting clusters were labelled as transition (TR), N3 (delta-dominant) and N2 (sigma/spindle-band enriched). For visualisation, substages were also projected into the PC space of a PCA fit on the entire recording. For the single-bout illustration, spindle events were detected on the broadband EEG within NREM by band-pass filtering the signal at 10–16 Hz (zero-phase 4th-order Butterworth), taking the Hilbert amplitude envelope, smoothing it (100-ms moving average) and thresholding at mean + 2 SD computed over all NREM samples; suprathreshold segments lasting 0.2–3 s were retained. Analyses were performed in Python using NumPy, SciPy and scikit-learn.

##### Sleep state exploration in non-mammalian species (jackdaw and lizard)

To illustrate Nyx’s applicability beyond established mammalian sleep architectures, we applied the pipeline to one jackdaw^21^ (EEG + accelerometer) and one tegu lizard recording (LFP(DVRA1) + EMG). Signals were preprocessed and feature-extracted following the standard rodent pipeline. Features comprised the EEG power spectrogram (0.5–40 Hz) combined with accelerometer activity for the jackdaw and EMG power for the lizard (5-80 Hz), z-scored and reduced via PCA; the first 3 principal components, each explaining >2% of the variance, were retained as the clustering feature space.

Because no a priori stage scheme exists for these species comparable to mammalian conventions, the number of clusters K was determined empirically. Candidate values K = 2– 8 were each fit with a Gaussian mixture model (fixed random seed) on the standardised PCA-derived feature space, with the principal-component count held fixed so that only K varied. Cluster-number support was assessed with two complementary, ground-truth-independent criteria. First, bootstrap stability: at each K we drew 20 random 80% subsamples (without replacement, fixed per-draw seeds), re-clustered each, and computed the adjusted Rand index between every pair of clusterings on their shared epochs (190 pairs), summarised as mean ± SD. Second, the gap statistic ^36^ compared the within-cluster dispersion of the data against 50 uniform reference datasets drawn from the feature-space bounding box, selecting K according to the standard one-standard-error rule. The bootstrap-stability peak and the gap-statistic recommendation converged on the reported K for each recording. We emphasise that this is a per-recording methodological demonstration; systematic cross-recording validation across the full species samples was beyond the scope of these exploratory examples.

##### Statistical analysis

Agreement between manual and Nyx biological variables estimation was quantified using three complementary measures. Intraclass correlation coefficients (ICC) were computed as ICC(A,1) -two-way mixed-effects, absolute-agreement, single measures (equivalent to Shrout & Fleiss ICC(2,1)^37^) - to quantify absolute agreement. Pearson correlation coefficient (r) was computed to characterize the strength of the linear correlation between methods independently of systematic offset; r and ICC are complementary, the gap between them indicating the contribution of directional bias to disagreement. Bias and limits of agreement (LoA, defined as bias ± 1.96 × SD of the paired differences) were derived from Bland-Altman analysis. Bias was defined as the mean of (Nyx − manual) differences, such that a positive bias indicates overestimation by the automatic method. The Wilcoxon signed-rank test was used to test the null hypothesis of zero median paired difference. As a sensitivity check for within-animal clustering, ICCs were additionally computed at the animal level by averaging values across recordings within each animal prior to estimation; concordance with per-recording estimates was treated as evidence that conclusions are not artifacts of pseudoreplication. ICC values were interpreted following Koo & Li (2016)^38^ as poor (<0.5), moderate (0.5-0.75), good (0.75-0.9), or excellent (>0.9). For the real-time implementation, the proportion of long REM bouts (> 30 s) between baseline and stimulation conditions was compared using a one-sided Fisher’s exact test. The 2×2 contingency table was constructed by classifying each REM bout in each condition as either long (> 30 s) or short (≤ 30 s). To compare REM entry frequency between baseline and stimulation conditions, a two-sided Poisson means test was used, with bout counts and recording durations as inputs, under the assumption that REM entries follow a Poisson process within each condition. Analyses were performed in Python (v3.12) using the pingouin (v0.6) and scipy.stats libraries.

### Generative AI disclosure

Generative AI tools were used in a limited capacity during preparation of this manuscript: for code assistance (debugging and refactoring), for editorial assistance with text (drafting and revision suggestions, with all scientific content authored and verified by the authors), and for the generation of one illustrative visual asset (the rat-on-shaker schematic in Fig. 5, generated with Google Imagen 3 via the Gemini platform).

## Data availability

Public datasets are available at their original sources: DOD-H and DOD-O at https://github.com/Dreem-Organization/dreem-learning-evaluation; MESA, CCSHS, and CHAT via the National Sleep Research Resource (https://sleepdata.org); ANPHY at https://osf.io/r26fh/overview; the Oxford mouse benchmark at https://zenodo.org/records/10200482; the Ellen Dash dataset at https://zenodo.org/records/5227351; the SIESTA dataset at https://zenodo.org/records/15322394; the Sippel Morris dataset at https://search.kg.ebrains.eu/instances/Dataset/830f33f1-cb6e-45ac-9203-11f23e97c273; and the Gulledge dataset at https://sleepdata.org/datasets/gulledge-2025.

The SleePing dataset was shared by collaborators and is described in Wilhelm et al. (2026), bioRxiv https://doi.org/10.64898/2026.06.26.734777. Jackdaw recordings (van Hasselt et al., 2025), ageing mouse recordings, and tegu lizard recordings were shared by collaborators and are available from the corresponding authors on reasonable request.

In-house rat recordings generated for this study will be made available without restriction upon publication. Manual scoring annotations used as reference throughout the validation are provided with each dataset, with the exception of the tegu lizard recordings.

## Code availability

The Nyx framework will be available as an open-source Python package on GitHub. Documentation, installation instructions, and tutorial notebooks will be provided. The graphical user interface will be distributed with the package. All analysis code used to generate the figures in this manuscript will be available with notebooks corresponding to each main and supplementary figure upon publication of the work.

## Acknowledgements

We thank Lina Okinina for animal care and drive assembly, and Sandra Bryne for manual scoring of the in-house DSI mouse dataset.

This work was supported by the European Research Council (ERC, 101117770), the grants 288478, 105428102, 332004 and 187615/V40 from the Research Council of Norway, the EU MSCA Cofund (Scientia Fellows II GA# 801133), and EMBO ALTF 423-2021.

Data from SleePing nap dataset were generously provided by Bernhard P. Staresina from the University of Oxford, UK. We thank Danqian Liu and Yujing Tian from the Center for Excellence in Brain Science and Intelligence Technology, Chinese Academy of Sciences, China, for sharing the ageing mouse sleep recordings. Data from jackdaws were generously provided by Peter Meerlo from the University of Groningen, Netherlands. Data from the wild rat and tegu lizard were generously provided by Paul-Antoine Libourel from the Center for Functional and Evolutionary Ecology (CEFE, CNRS UMR 5175), Montpellier, France.

The human dataset MESA, CCSHS, and CHAT were obtained through the National Sleep Research Resource which is supported by the U.S. National Institutes of Health, National Heart Lung and Blood Institute (R24 HL114473, 75N92019R002). The Multi-Ethnic Study of Atherosclerosis (MESA) Sleep Ancillary study was funded by NIH-NHLBI Association of Sleep Disorders with Cardiovascular Health Across Ethnic Groups (RO1 HL098433). MESA is supported by NHLBI funded contracts HHSN268201500003I, N01-HC-95159, N01-HC-95160, N01-HC-95161, N01-HC-95162, N01-HC-95163, N01-HC-95164, N01-HC-95165, N01-HC-95166, N01-HC-95167, N01-HC-95168 and N01-HC-95169 from the National Heart, Lung, and Blood Institute, and by cooperative agreements UL1-TR-000040, UL1-TR-001079, and UL1-TR-001420 funded by NCATS. The Childhood Adenotonsillectomy Trial (CHAT) was supported by the National Institutes of Health (HL083075, HL083129, UL1-RR-024134, UL1 RR024989). The Cleveland Children’s Sleep and Health Study (CCSHS) was supported by grants from the National Institutes of Health (RO1HL60957, K23 HL04426, RO1 NR02707, M01 Rrmpd0380-39).

