## Supplementary Information for "Nyx: a flexible framework for sleep scoring across species, lifespan and modalities"

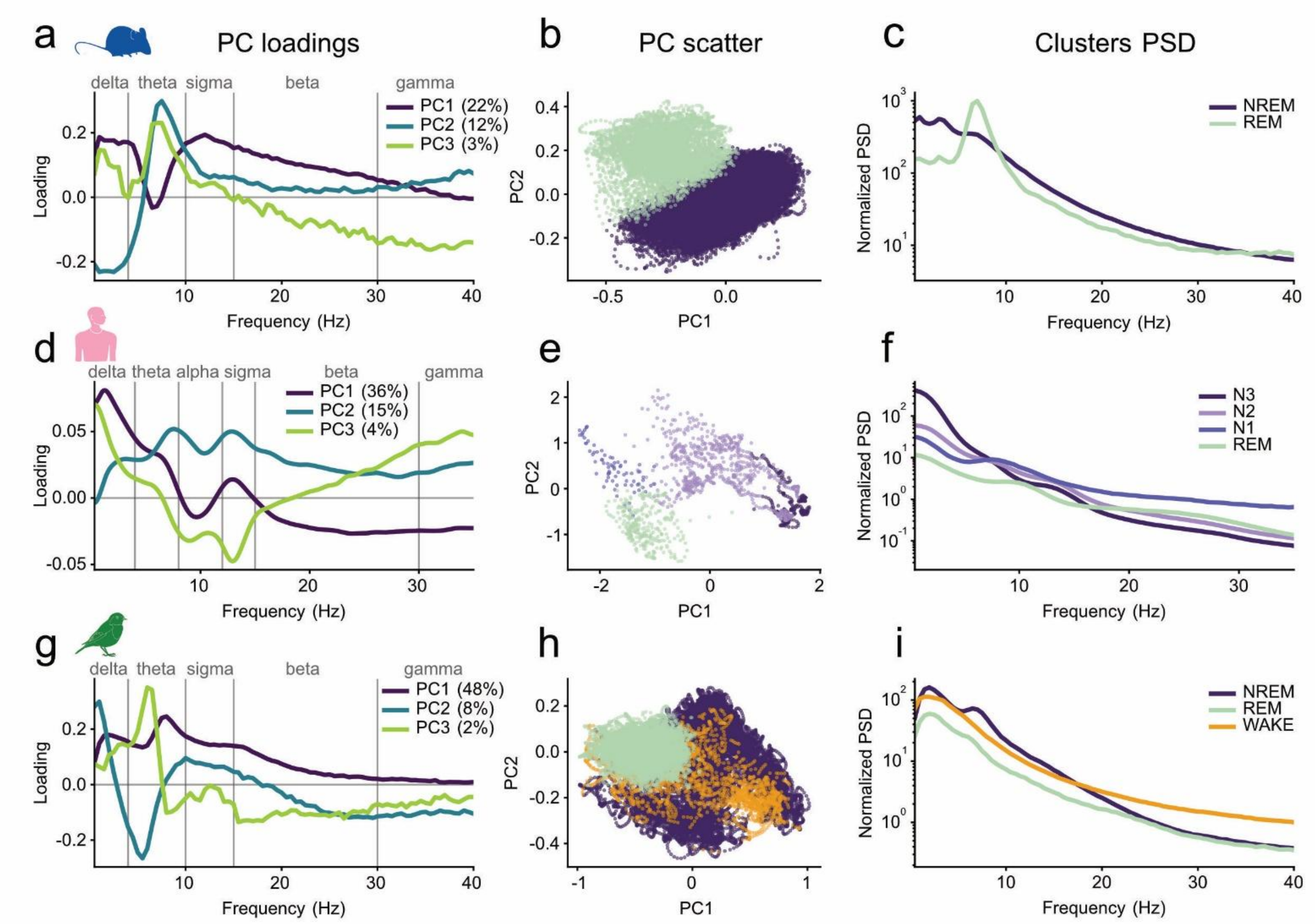


Fig. S1: Anatomy of the PCA space representation across species

Structure of the principal-component decomposition used for automatic sleep scoring in three representative recordings: mouse (a–c), human (d–f) and jackdaw (g–i). For each recording, PCA was computed on the log-power EEG spectrogram over sleep epochs. Panels show the three principal components with ≥2% variance explained as loadings across frequency (a, d, g), the epoch distribution in the PC1–PC2 plane coloured by Nyx cluster/stage assignment (b, e, h), and the median power spectral density per cluster (c, f, i). Across species, PCA recovers physiologically interpretable spectral axes aligned with canonical frequency bands, from which cluster identity emerges without stage labels provided as input. **a**–**c, Mouse** (recording oxford_mouse_benchmark_dataset_test SampleFile_C; 47,640 one-second sleep epochs; NREM 41,386 [86.9%], REM 6,254 [13.1%]; three PCs retained for scoring). **a,** PC loadings across frequency bins (0.5–40 Hz). PC1 is a broadband-power axis, positive across all bands but with a distinct negative notch in theta (~6.5 Hz); PC2 is a delta(−)/theta(+) contrast peaking exactly there (+0.30 at 7.5 Hz). PC3 is positive at theta, negative through beta and gamma. **b,** PC1–PC2 scatter, one point per sleep epoch coloured by cluster assignment (NREM, REM); PC1+PC2 capture 34.0% of the total spectral variance. The components cleanly separate delta-dominant NREM (high PC1, low PC2) from theta-dominant REM (low PC1, high PC2). **c,** Median PSD per cluster (0.5–40 Hz, log-power). NREM power is dominated by low frequencies, REM shows the characteristic theta peak at ~7 Hz. **d**–**f, Human** (recording ANPHY, EPCTL03, C4 EEG, ~7.0 h, 1,289 sleep epochs). **d,** PC loadings across frequency bins (0.5–35 Hz). PC1 is dominated by low-frequency power: strongly positive across the delta band, falling to near zero through theta/alpha and slightly negative across beta–gamma — a slow-wave axis separating high-delta epochs from faster, lower-amplitude activity. PC2 is near zero in delta and positive across the faster bands, with twin peaks at theta (~8 Hz) and sigma (~13 Hz) and a sustained mild-positive tail through beta–gamma. PC3 is positive at both frequency extremes (delta and gamma) but strongly negative through alpha–sigma, with its trough near 13 Hz — a contrast of sigma-band (spindle) power against both slow and high-frequency power. **e,** PC1–PC2 scatter for 1,289 sleep epochs coloured by automatic stage assignment (N1–3, REM); PC1+PC2 explain 51% of variance, and the four stages form a graded trajectory from REM to N3. **f,** Median PSD per stage (0.5–35 Hz, log-power). The stages separate mainly by slow-wave content — N3 has by far the highest delta power and the steepest decrease, N2 and N1 are intermediate, and REM the lowest — while the ordering inverts above ~10 Hz, where N1 carries the most beta/gamma power and N3 the least. **g–i, Jackdaw** (recording L6_06052018_155026_gwOT, day 0, combined-hemisphere EEG). **g,** PC loadings across frequency bins (0.5–40 Hz). PC1 is an all-positive broadband power axis peaking in delta–theta; PC2 contrasts delta/sigma (positive) against theta and higher frequencies (negative); PC3 isolates a narrow ~6 Hz theta peak. **h,** PC1–PC2 scatter of sleep epochs coloured by automatic cluster assignment (NREM, REM, WAKE); PC1+PC2 jointly capture 56% of the variance. **i** Median PSD per cluster (0.5–40 Hz, log-power). NREM shows the highest low-frequency power, WAKE the intermediate spectrum with sustained higher-frequency content, and REM the lowest overall power.

In all loading panels (a, d, g), lines are Gaussian-smoothed and sign-oriented so each PC's largest-magnitude loading is positive. Vertical lines and top labels demarcate canonical frequency bands (delta 0.5–4, theta 4–10, sigma 10–15, beta 15–30, gamma 30–40 Hz; the human panel additionally labels alpha 8–12 Hz). PSDs (c, f, i) are normalised by global mean power per recording (a.u.), preserving relative amplitudes between clusters.

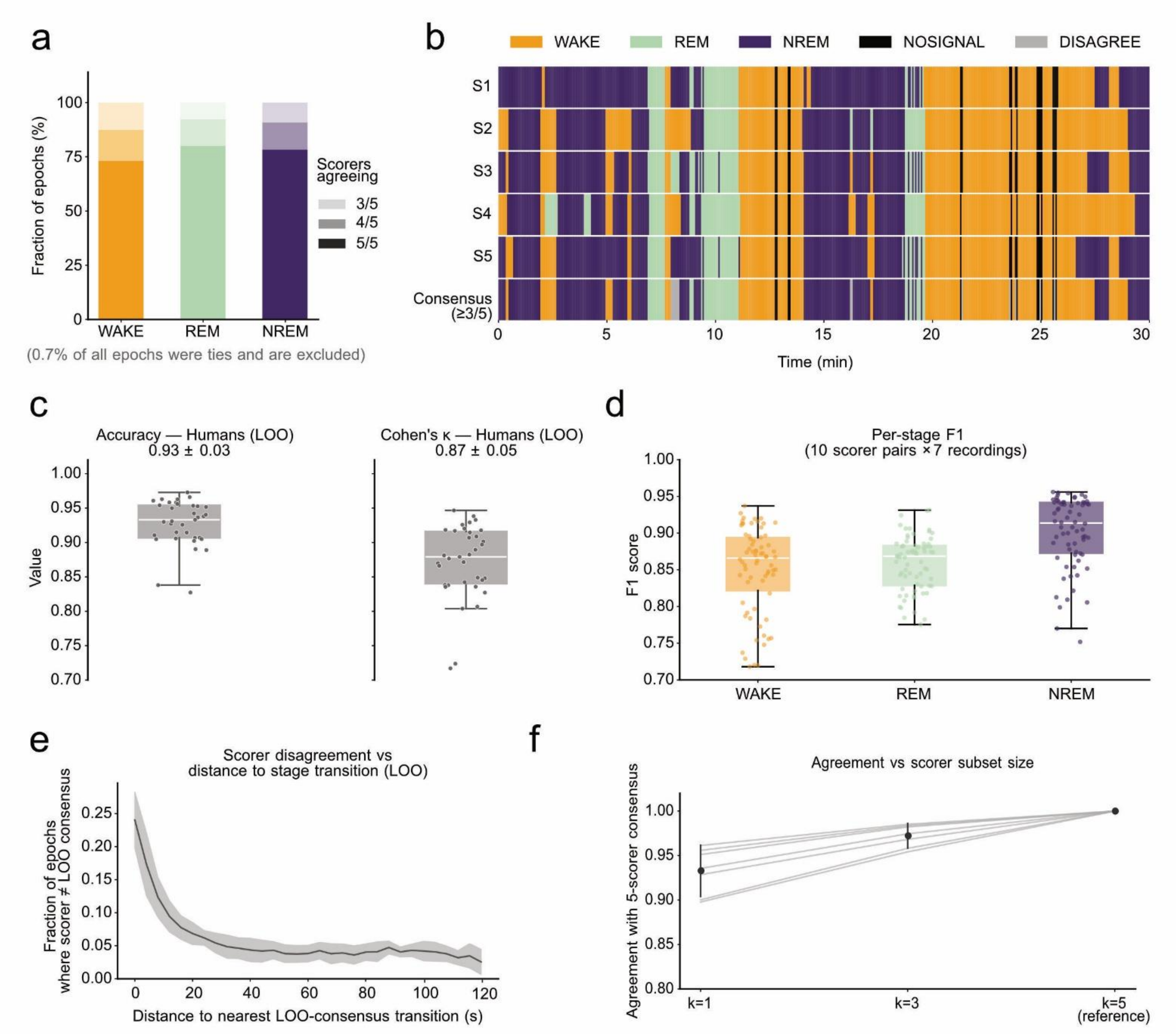


Fig. S2: Inter-scorer agreement on the in-house DSI rodent recordings

**a,** Agreement fraction breakdown across all consensus epochs pooled from the seven DSI rat recordings (n = 40,666 epochs of 4 s). For each stage (WAKE, REM, NREM), the proportion of epochs labelled unanimously by all five scorers, by a 4/5 majority, or by a 3/5 majority. The DISAGREE category (0.7% of epochs) corresponds to epochs receiving no majority label. **b,** Per-scorer hypnograms of a representative 30-minute window from one recording, showing transition-localised disagreement between scorers on otherwise concordant stage sequences. **c,** Per-scorer accuracy (left) and Cohen's κ (right) against a leave-one-out (LOO) consensus of the remaining four scorers, across the 35 scorer–recording combinations (5 scorers × 7 recordings). **d,** Pairwise per-stage F1 across the 10 scorer pairs and 7 recordings (n = 70 per stage). NREM shows the highest pairwise agreement (0.901 ± 0.049); WAKE (0.850 ± 0.059) and REM (0.859 ± 0.037) are comparable at lower values. **e,** Fraction of epochs at which a scorer disagreed with the LOO consensus as a function of distance from the nearest manual stage transition. Disagreement drops from 24% at the transition itself to below 10% within 12 s and plateaus at 4.6% by 120 s. **f,** Agreement with the full five-scorer consensus as a function of subset size k, from k = 1 (n = 35) to k = 4 (n = 35). A three-scorer subset already reaches 0.972 ± 0.015, up from 0.933 ± 0.030 at k = 1. Boxplots show median, IQR, and 1.5 × IQR whiskers. Aggregate values reported as mean ± SD.

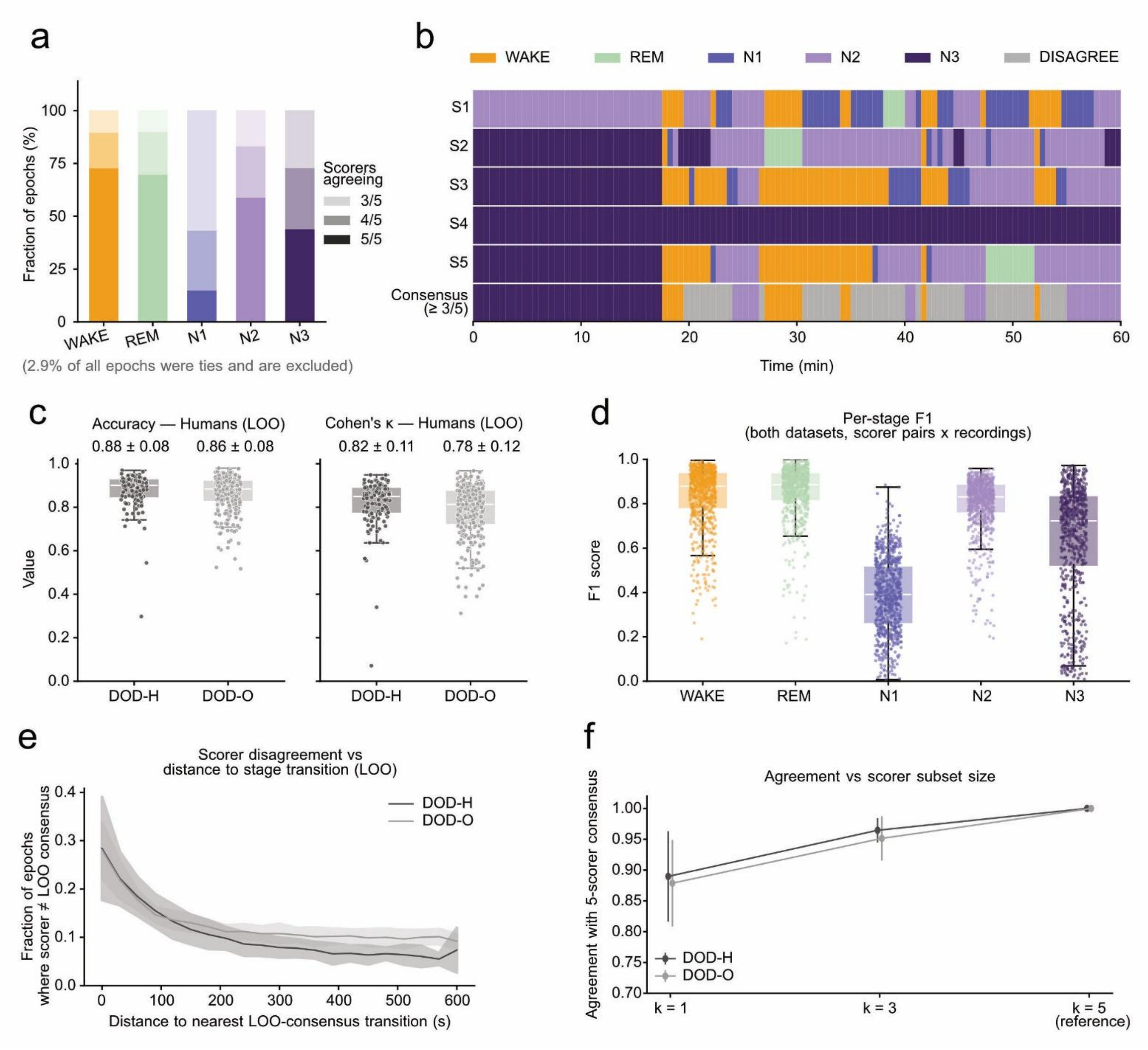


Fig. S3: Inter-scorer agreement on the DOD-H and DOD-O human polysomnography recordings

Same analyses as Fig. S1, applied to the two Dreem Open Datasets — DOD-H (healthy subjects) and DOD-O (subjects with suspected obstructive sleep apnea) — totaling 78,011 valid epochs of 30 s after excluding 2.9% of epochs as ties. **a,** Agreement fraction breakdown across the five AASM stages (WAKE, REM, N1, N2, N3), pooled across both datasets. Unanimous five-scorer agreement reveals a steep gradient: 73% WAKE, 70% REM, 59% N2, 44% N3, and just 15% N1. **b,** Per-scorer hypnograms of a representative recording, showing a higher density of disagreement on REM and N1 epochs than on other stages. **c,** Per-scorer LOO accuracy (left) and Cohen's κ (right), reported separately for DOD-H and DOD-O. Lower than DSI rodent values, as expected for five-stage human scoring. **d,** Pairwise per-stage F1 pooled across both datasets, confirming N1 as the hardest stage for human scorers (median F1 = 0.39); WAKE, REM, and N2 show substantially higher pairwise agreement (median > 0.80); N3 is intermediate (median F1 = 0.72). **e,** Fraction of epochs at which a scorer disagreed with the LOO consensus as a function of distance from the nearest manual stage transition, shown separately for DOD-H and DOD-O. Disagreement declines from ~30% at the transition itself and plateaus below 10% beyond 210 s for DOD-H and 450 s for DOD-O. **f,** Agreement with the full five-scorer consensus as a function of subset size k, rising from ~0.89 at k = 1 to ~0.95 at k = 3 for both datasets.

Boxplots show median, IQR, and 1.5 × IQR whiskers. Aggregate values reported as mean ± SD.

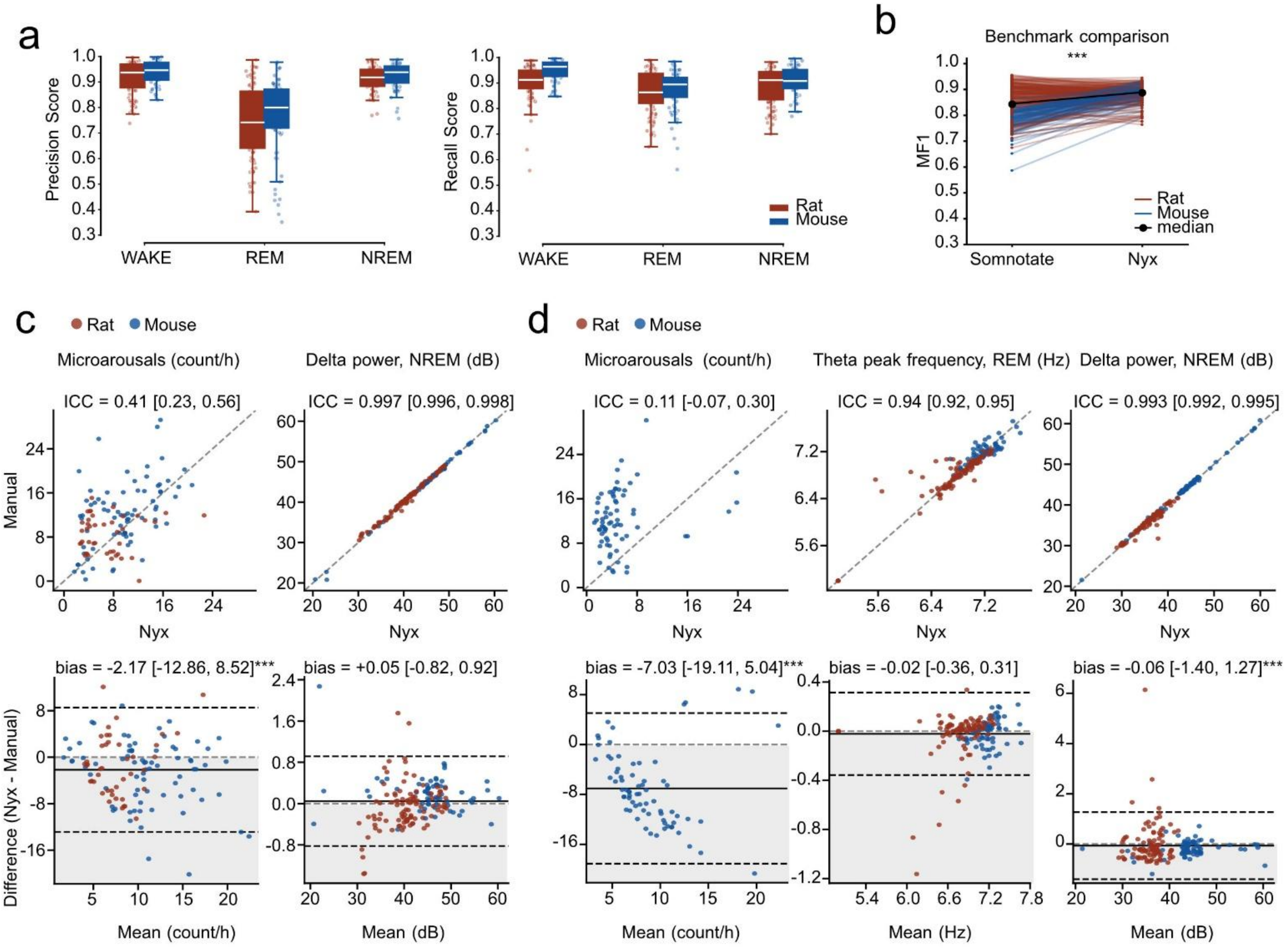


Fig. S4: Per-stage precision and recall, benchmark comparison, and additional sleep variables supporting Fig. 2

**a,** Per-stage precision and recall across recordings, split by species. REM recall exceeds REM precision in both rats and mice, underlying the REM overscoring asymmetry. **b,** Slopegraph of macro-F1 (MF1; unweighted mean of per-stage F1) for each recording scored by Somnotate (left) and Nyx (right) against manual expert scoring, pooled across five datasets (n = 239 recordings from 35 animals; see Table S4 for per-dataset composition and agreement metrics). Each thin line is one recording, coloured by species (blue, mouse; red, rat); thick black line connects medians. Two-sided Wilcoxon signed-rank test on per-animal mean MF1: Nyx vs Somnotate, $p = 1.8 \times 10^{-6}$, rank-biserial $r = 0.84$ (median MF1 0.889 vs 0.805, median difference +0.080; Nyx ≥ Somnotate in 28/35 animals; ***$p < 0.001$). The effect was carried by mice ($p = 1.9 \times 10^{-6}$, $r = 1.00$; Nyx higher in all 20 animals), with no significant difference in rats ($p = 0.60$, $r = 0.17$; median difference +0.001), despite Somnotate having been trained exclusively on mouse recordings. **c,** Agreement on additional summary sleep variables across baseline recordings. Scatter (top) and Bland–Altman (bottom) pairs for, left to right: microarousal rate and delta power in NREM. **d,** Agreement on the same set of variables across manipulation recordings, same layout as c, with the addition of theta peak frequency in REM (left to right: microarousals, theta peak frequency, delta power). Bias convention: Nyx − manual (positive = overestimation by Nyx).

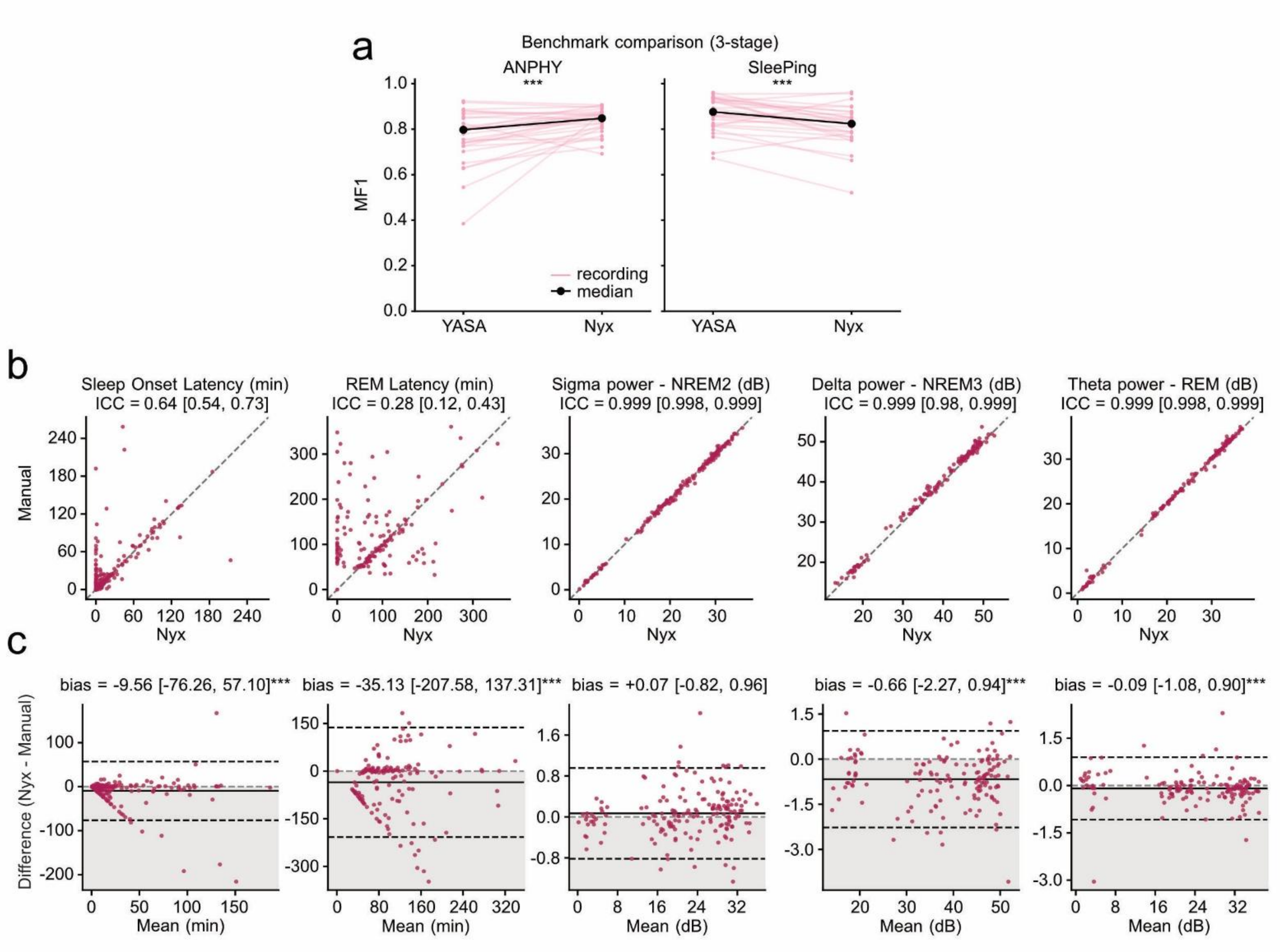


Fig. S5: Benchmark comparison and agreement on additional summary sleep metrics, supporting Fig. 3

**a,** Slopegraph of macro-F1 (MF1) for each recording scored by YASA (left) and Nyx (right) against manual expert scoring on human recordings, split by dataset (ANPHY, left; SleePing, right; n = 29 and 28 recordings respectively). Each thin line is one recording; thick black line connects medians. Two-sided Wilcoxon signed-rank test on per-recording MF1 under the 3-stage AASM scheme: Nyx exceeded YASA on ANPHY (***$p < 0.001$, $r = 0.70$; median MF1 0.848 vs 0.797), YASA exceeded Nyx on SleePing (***$p < 0.001$, $r = 0.69$; median MF1 0.876 vs 0.824), with no significant difference in the pooled comparison ($p = 0.89$; $n = 57$). Results were consistent under 4- and 5-stage schemes (Table S8). YASA was applied without retraining to datasets outside its original training set. **b,** Scatter plots of Nyx (*y*) vs manual (x) values across all recordings for, left to right: sleep onset latency, REM onset latency, sigma power in N2, delta power in N3, and theta power in REM. Identity line shown for reference; ICC(A,1) values inset. **c,** Bland–Altman plots for the same metrics in the same order, showing the difference (Nyx − manual) against the mean of the two measurements. Mean bias and 95% limits of agreement indicated; bias convention: Nyx − manual (positive = overestimation by Nyx). Spectral metrics are near-perfectly recovered (ICC ≥ 0.999, |bias| < 0.7 dB); REM latency shows the poorest agreement (ICC = 0.28), reflecting the emergent nature of both anchor points (first sleep epoch and first REM epoch) under Nyx's unsupervised stage assignment.

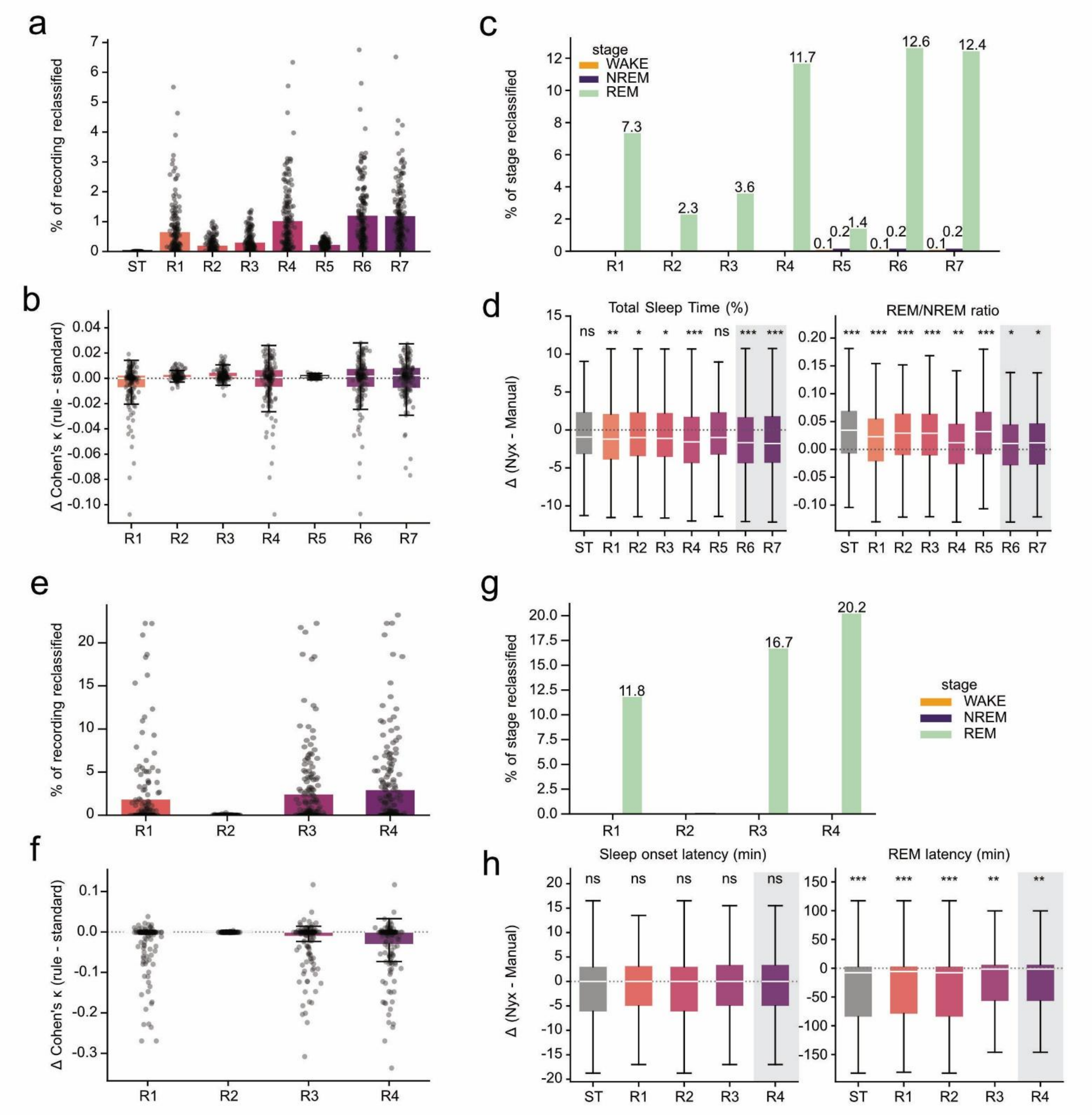


Fig. S6: Post-hoc stage-cleaning rules act almost exclusively on REM, producing targeted shifts in REM-dependent biological metrics

Rules were applied to the automatic hypnogram only. Standard (ST) is the un-ruled baseline. R1 reclassifies any REM segment flanked by WAKE to WAKE; R2 applies the same rule only to short REM segments (<20 s rodents, <30 s humans); R3 reclassifies REM following a long WAKE bout (same thresholds) to WAKE; R4 reclassifies REM following any WAKE bout to WAKE. Rodents only: R5 enforces a 4 s minimum bout length; R6 = R4 then R5; R7 = R5 then R4 (full details in Supplementary Methods). Top rows: rodents (a–d, 179 recordings from 10 datasets, seven rules). Bottom rows: humans (e–h, 179 recordings from five datasets, four rules).

**a,e,** Rules touch only a small fraction of the recording. Percentage of recording reclassified by each rule relative to Standard (bar = mean, dots = individual recordings). Rules alter ≤1.2% of recording time in rodents and ≤2.9% in humans.

**b,f,** Overall Nyx–manual agreement is essentially unchanged, with a small tail on aggressive rules. Paired per-recording change in Cohen's κ relative to Standard. Boxes show median and IQR, whiskers 1.5×IQR, dots individual recordings; dotted line marks no change. Median Δκ is near zero for every rule in both species. In rodents, the negative tail is confined to the aggressive rules: 9.5% of recordings (17/179) lost >0.02 κ under R1, R4, and R6 (6.1% under R7) and 2.2% (4/179; worst −0.108) fell below −0.05, versus 0% under R2, R3, and R5; the tail was species-dependent (19.7% of mouse vs 1.9% of rat recordings under R4/R6). Humans showed a substantially larger tail under REM-adjacent-to-wake rules: under R4, 27.4% lost >0.02 κ and 22.3% >0.05 (worst −0.336), with R1 and R3 comparable (14.0% and 16.8% <−0.05), while R2 left every recording unchanged. The same recordings drove the tail across all aggressive rules, and the effect was dataset-dependent (SleePing improved on average, +0.005 κ; MESA carried the heaviest tail, mean −0.039 κ). Unlike in rodents, restricting REM removal to long preceding WAKE bouts (R3) did not protect against this tail.

**c,g,** The small footprint is concentrated on REM. Reclassifications expressed as a percentage of each baseline stage's total duration in the Standard hypnogram, pooled across recordings. Reclassification is essentially confined to REM (up to ≈13% under R4, R6, R7 in rodents and ≈20% under R4 in humans); WAKE and NREM are left untouched.

**d,h,** REM-dependent biological metrics shift accordingly. Bias of the automated estimate relative to manual (Nyx − manual) for two variables per species. Rodents (D): Total Sleep Time (%, left) and REM/NREM ratio (right); REM-removal rules R6 and R7 highlighted. Humans (H): Sleep onset latency (min, left) and REM latency (min, right); REM-removal rule R4 highlighted. Dotted line marks zero bias. Standard over-estimates the REM/NREM ratio in rodents and under-estimates REM latency in humans; REM-removal rules reduce these biases at the cost of a small negative TST bias in rodents.

Stars: two-sided Wilcoxon signed-rank test of the paired per-recording difference against 0 (*** $p < 0.001$, ** $p < 0.01$, * $p<0.05$)

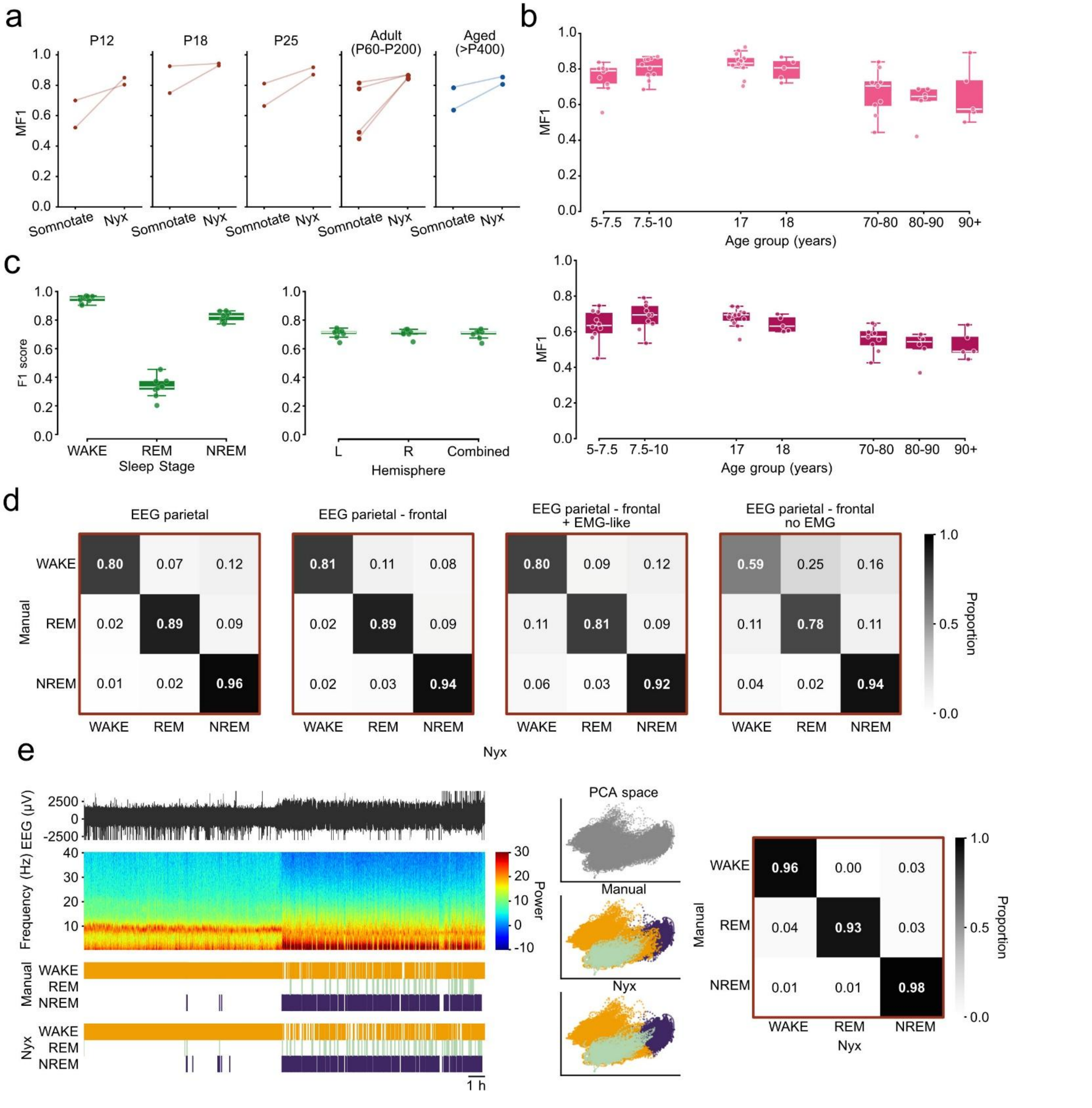


Fig. S7: Per-stage breakdowns, configuration-level controls, and a representative scoring example supporting Fig. 4

**a,** Slopegraph of macro-F1 (MF1) for each recording scored by Somnotate (left) and Nyx (right) against manual expert scoring, across the lifespan cohort (n = 12 recordings from 8 animals; developmental rats at P12, P18, P25 and adult; 14-month aged mice). Each thin line is one recording, coloured by species (red, rat; blue, mouse). Two-sided Wilcoxon signed-rank test on per-animal mean MF1: Nyx vs Somnotate, p = 0.0078 (median MF1 0.863 vs 0.694, median difference +0.171; Nyx > Somnotate in all 8 animals). Per-age-group values shown descriptively (n = 2-4 recordings per group). Somnotate was trained exclusively on its original mouse dataset and applied without retraining (see Table S11 for per-dataset agreement metrics). **b,** Human lifespan performance under the 4- (Left) and 5-stage (Right) AASM schemes, pooled across CHAT, CCSHS, and MESA (n = 60 recordings across 7 age bins). **c,** Jackdaw per-stage and hemisphere breakdowns across the sleep-deprivation protocol (n = 9 birds, 9 recordings per condition). Left: per-stage F1 pooled across conditions, Right: macro F1 for left-hemisphere, right-hemisphere, and combined-channel analyses, showing near-identical performance. **d,** Confusion matrices for the three EMG configurations in n = 4 adult rats scored with the parietal–frontal derivation: separate EMG channel, EMG-like signal derived from EEG and LFP channels, and no EMG. Removing EMG produces a WAKE/REM confusion visible as off-diagonal mass between the two stages, underlying the WAKE F1 collapse (0.47 ± 0.33) reported in Fig. 4f. **e,** Representative scoring from a wild rat recorded with a telemetry PNano system from Manitty, with a clear circadian structure visible in the hypnogram. Left: raw EEG trace, EEG time–frequency spectrogram, manual and Nyx hypnograms, and PC-space projection of epochs coloured by manual and Nyx labels. Right: confusion matrix showing per-stage F1 ≥ 0.93 on the diagonal across all stages.

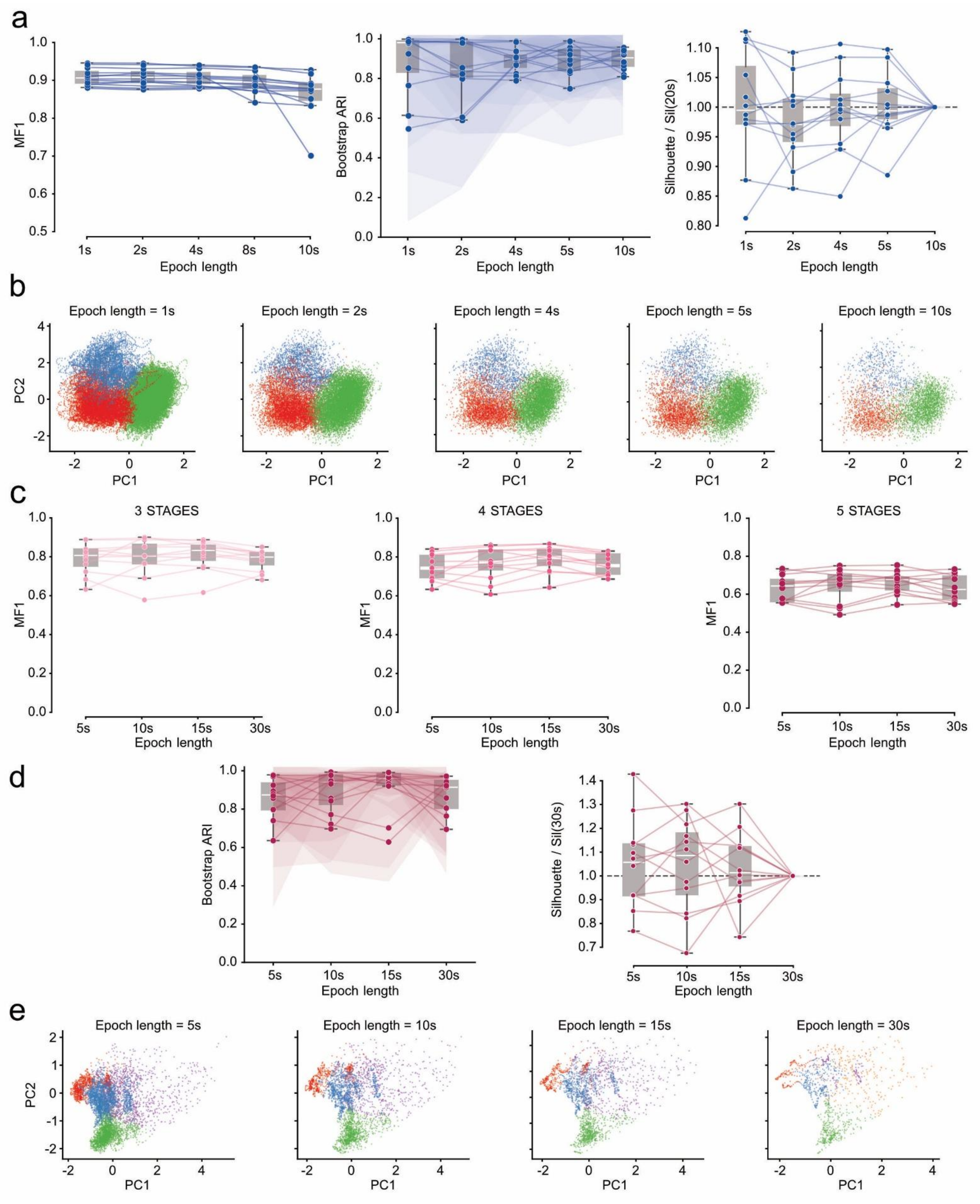


Fig. S8: Nyx classification and cluster structure are stable across epoch lengths in rodents and humans

Rodents **(a–c)** and humans **(d–f)**; Nyx applied to the sleep clustering step in both species. To test whether Nyx depends on the conventional epoch durations used in manual scoring (4 s in rodents, 30 s in humans), we repeated the analysis on the parietal EEG of the twelve Oxford mice at five raw bin sizes (2, 4, 8, 10, 20 s; effective resolutions 1–10 s) and on the twelve DOD-H human recordings at four scored epoch lengths (5, 10, 15, 30 s). Boxplots show median, IQR, and 1.5× IQR whiskers; individual recordings overlaid as points. **a,** Rodent classification agreement is stable across epoch lengths.

Macro F1 relative to manual scoring across rodent epoch durations. Agreement was essentially unchanged across the 2–10 s range (median MF1 = 0.91, 0.91, 0.91, 0.89) and showed only a modest decrement at 20 s (0.87). **b,** Rodent cluster structure is stable across epoch lengths. Left: subsampling bootstrap ARI, quantifying cluster stability independently of manual labels; median ≥ 0.85 at every epoch length. Right: silhouette score under a sample-size-matched, reference-configuration GMM, normalised within recording relative to the longest epoch length; no systematic improvement or degradation across durations. **c,** Cluster geometry is preserved across epoch lengths in rodents. Two-dimensional PC-space projections of one representative rodent recording at each epoch duration, coloured by Nyx cluster assignment. **d,** Human classification agreement is stable across epoch lengths and scoring schemes. Macro F1 across human epoch durations under the 3-stage AASM scheme (left), 4-stage (centre), and 5-stage (right). Performance was flat across the 5–30 s range under all three schemes (5-stage median MF1 = 0.65, 0.67, 0.66, 0.63; 4- and 3-stage schemes equivalently flat). **e,** Human cluster structure is stable across epoch lengths. As in (b), for humans (median ARI = 0.87, 0.94, 0.96, 0.91 at 5, 10, 15, 30 s). Between-recording spread was systematically wider than in rodents at every epoch length, reflecting the harder partitioning problem the human sleep step solves: four overlapping sub-clusters (N1/N2/N3/REM) as opposed to three (W/NREM/REM), a more variable EEG substrate across subjects, and an ambiguous N1/N2 boundary. That the algorithm recovers a stable partition at every epoch length indicates the wider human dispersion is a property of the data. **f,** Cluster geometry is preserved across epoch lengths in humans. As in (c), for one representative human recording at each epoch duration. Full statistical values in Table S15.

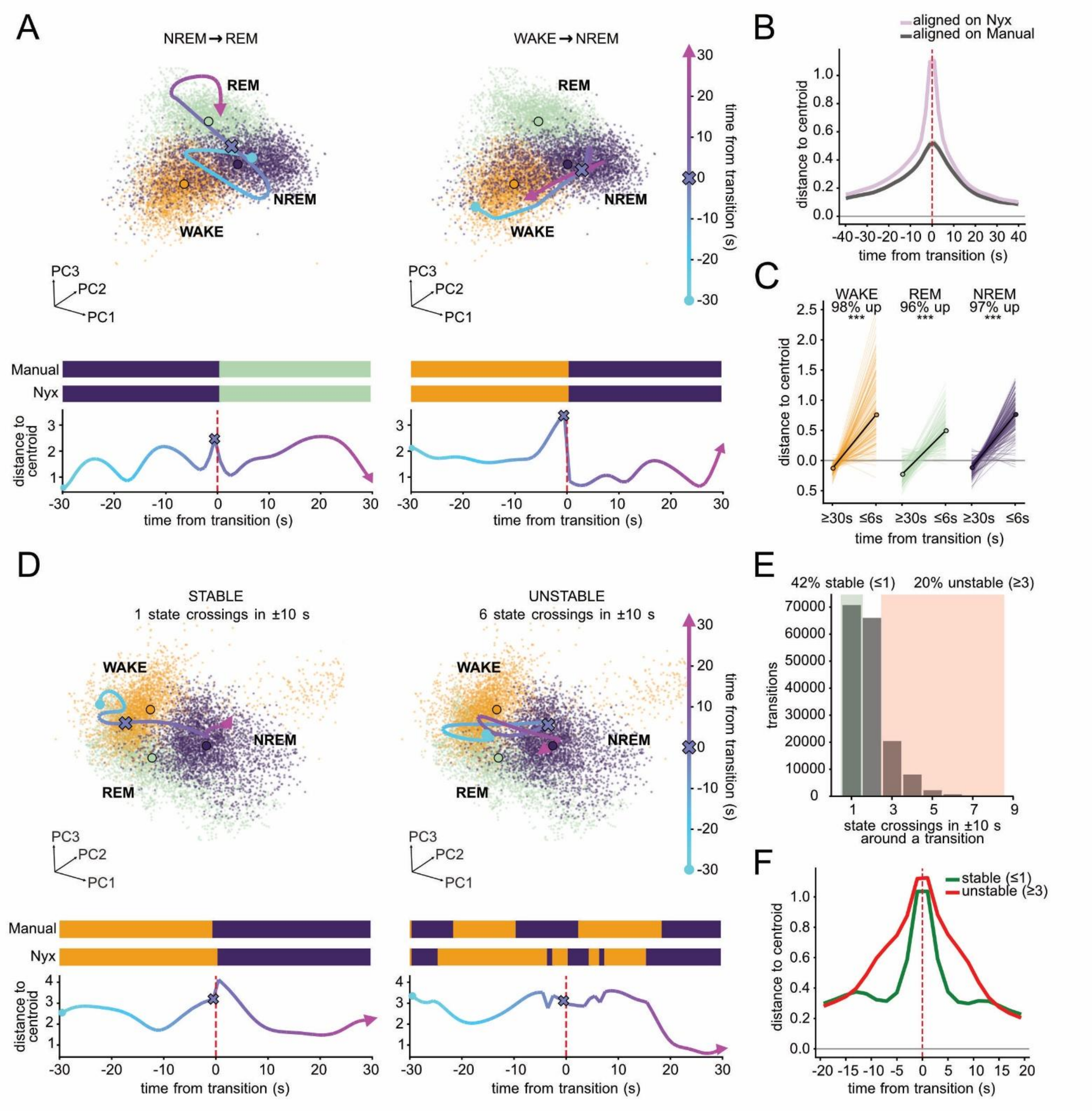


Fig. S9: State transitions are continuous passages through the Nyx PC space

**a,** Single transitions unfold as continuous trajectories. Two representative transitions from one recording (NREM→REM, WAKE→NREM). Top: epoch cloud in PC1–PC2–PC3 coloured by state (WAKE orange, NREM purple, REM green), with the three state centroids labelled and the ±30 s trajectory overlaid, colour-coded by time from the transition (cyan→magenta, colorbar); "o" marks the trajectory start, "X" the transition (t=0), and the arrowhead the end. Middle: Manual and Nyx hypnograms over the same window. Bottom: distance from each epoch to its assigned state centroid (Mahalanobis, a.u.) on the same time-from-transition colour scale. The state travels continuously between centroids while the label switches at a single instant. **b,** The distance signature is consistent across transitions. Event-triggered average of the distance to the assigned state centroid across ±40 s around every transition, pooled over 179 recordings (mean ± s.e.m.), aligned on Nyx (lilac) and Manual (grey). Distances are Mahalanobis and z-scored within recording, so they are comparable across states and across animals. The Nyx alignment confirms the geometry the pipeline uses. The Manual alignment is the informative one: the same peri-transition rise-peak-settle profile emerges around an independently defined transition time, so the continuous geometry is a property of the underlying state change and not of the clustering. **c,** Near-transition epochs sit farther from the centroid than far-from-transition epochs. Within recording and within state, near-transition epochs ($|\Delta t| \leq 6$ s) are farther from the assigned centroid than far-from-transition epochs ($|\Delta t| \geq 30$ s). Thin lines are recordings coloured by state (WAKE, NREM, REM); bold black lines are medians. Distances are Mahalanobis and z-scored within recording, so baselines are equalised across states and across animals. Near > far in ~96–98% of recordings per state (Wilcoxon $p \approx 10^{-30}$). **d,** Stable and unstable transitions have distinct geometric signatures. Two example transitions from one recording. Left: a stable transition (single crossing, direct path between centroids, one quick-settling distance peak). Right: an unstable transition (six crossings, path looping between WAKE and NREM, oscillating and slow-settling distance). The flicker is visible in both Manual and Nyx hypnograms. Panel structure follows (A): top, epoch cloud and trajectory in PC1–PC2–PC3; middle, Manual and Nyx hypnograms; bottom, distance to the assigned centroid (Mahalanobis, a.u.). **e,** Transitions span a continuum of stability. Distribution of Nyx transition crossings within ±10 s of each Nyx transition anchor, pooled over 179 recordings. Crossings ≥1 by definition (the anchor is included). ~42% of transitions are stable (≤1 crossing) and ~20% are unstable (≥3 crossings). **f,** Stable and unstable transitions produce distinct pooled profiles. Event-triggered average of the distance to the assigned state centroid across ±20 s, split by stable (≤1 crossing) and unstable (≥3 crossings) transitions, pooled over 179 recordings (mean ± s.e.m.). Distances are Mahalanobis and z-scored within recording. Stable transitions produce a sharp, quick-settling peak; unstable transitions produce a broad, slow-settling peak.

### Supplementary Tables

#### Supplementary Table 1. Datasets.

| Dataset | Species | Strain / Condition | N individuals | N recordings | N excluded | Setup | Sampling rate | Average duration | Scoring epoch | Source | Ref. | Fig. |
|---|---|---|---|---|---|---|---|---|---|---|---|---|
| ***Rodent datasets*** | | | | | | | | | | | | |
| Boccara lab — DSI | Rat | WT (adult, telemetry) | 3 | 7 | — | EEG (P) + EMG | 1000 Hz | ~7 h | 4 s | In-house | — | 2 |
| Boccara lab — tethered | Rat | WT (P12) | 2 | 2 | — | EEG (P+F) + EMG + LFP | 48 kHz | 30 min – 1 h | 4 s | In-house | — | 4, 6 |
| | Rat | WT (P18) | 2 | 2 | — | EEG (P+F) + EMG + LFP | 48 kHz | 30 min – 1 h | 4 s | In-house | — | 4, 6 |
| | Rat | WT (P25) | 2 | 2 | — | EEG (P+F) + EMG + LFP | 48 kHz | 15 min - 45 min | 4 s | In-house | — | 4, 6 |
| | Rat | WT (adult) | 2 | 4 | — | EEG (P+F) + EMG + LFP | 48 kHz | 15-30 min | 4 s | In-house | — | 4, 6 |
| | Rat | WT (P34) | 1 | 2 | — | EEG (P+F) + EMG | 20 kHz | ~3 h | — | In-house | — | 5 |
| Oxford Mouse test | Mouse | WT (test) | 6 | 6 | — | EEG (P+F) + EMG | 256 Hz | 24 h | 4 s | Open | [1,2] | 2, 4 |
| Oxford Mouse SD | Mouse | SD | 6 | 12 | — | EEG (P+F) + EMG | 256 Hz | 12 h | 4 s | Open | [1–3] | 2, 4 |
| Oxford Mouse opto | Mouse | Optogenetic | 11 | 22 | — | EEG (P+F) + EMG | 256 Hz | 24 h | 4 s | Open | [1,2,4] | 2 |
| Ellen Dash | Rat | WT | 9 | 42 | 6 | EEG (P+F) + EMG | 250 Hz | 24 h | 4 s | Open | [5,6] | 2 |
| SIESTA | Mouse | WT | 10 | 10 | — | EEG (P+F) + EMG | 400 Hz | 24 h | 5 s | Open | [7,8] | 2 |
| | Mouse | SCN1a (circadian) | 6 | 6 | — | EEG (P+F) + EMG | 400 Hz | 24 h | 5 s | Open | [7,8] | 2 |
| | Mouse | APP-PS1 (Alzheimer) | 6 | 6 | 1 | EEG (P+F) + EMG | 400 Hz | 24 h | 5 s | Open | [7,8] | 2 |
| | Mouse | NmsVgat (arrhythmia) | 4 | 4 | — | EEG (P+F) + EMG | 400 Hz | 24 h | 5 s | Open | [7,8] | 2 |
| | Mouse | WT (telemetry) | 3 | 3 | — | EEG(F) + EMG | 250 Hz | 24 h | 5 s | Open | [7,8] | 2 |
| Sippel Morris | Mouse | WT (C57BL/6) | 10 | 40 | — | EEG (O+F) + EMG | 400 Hz | 24 h | 4 s | Open | [9,10] | 2 |

| Dataset | Species | Strain / Condition | N individuals | N recordings | N excluded | Setup | Sampling rate | Average duration | Scoring epoch | Source | Ref. | Fig. |
|---|---|---|---|---|---|---|---|---|---|---|---|---|
| | Mouse | Cyclin D2-/- | 10 | 40 | — | EEG (O+F) + EMG | 400 Hz | 24 h | 4 s | Open | [9,10] | 2 |
| Gulledge | Rat | WT (saline) | 4 | 33 | 3 | EEG (1 bipolar) + EMG | 500 Hz | 24 h | 10 s | Open | [11–13] | 2 |
| | Rat | Oxycodone | 11 | 129 | — | EEG (1 bipolar) + EMG | 500 Hz | 24 h | 10 s | Open | [11–13] | 2 |
| Ageing mice | Mouse | WT (aged, P420) | 2 | 2 | — | EEG (P+F) + EMG | 1000 Hz | 24 h | 5 s | Collaborator | — | 4 |
| ***Human datasets*** | | | | | | | | | | | | |
| ANPHY | Human | Healthy | 29 | 29 | — | 83 EEG + EMG + EOG + ECG | 1000 Hz | Overnight | 30 s | Open | [14,15] | 3 |
| DOD-H | Human | Healthy | 25 | 25 | 3 | 12 EEG + EMG + EOG + ECG | 250 Hz | Overnight | 30 s | Open | [16,17] | 3 |
| DOD-O | Human | OSA suspicion | 55 | 55 | 6 | 8 EEG + EMG + EOG + ECG | 250 Hz | Overnight | 30 s | Open | [16,17] | 3 |
| MESA | Human | Healthy / SDS | 56 | 56 | 5 | 3 EEG + EMG + EOG + ECG | 256 Hz | Overnight | 30 s | Open | [18,19] | 3 |
| | Human | Healthy / SDS (aged, 70+) | 25 | 25 | 5 | 3 EEG + EMG + EOG + ECG | 256 Hz | Overnight | 30 s | Open | [18,19] | 4 |
| SleePing | Human | Healthy (day-time nap) | 29 | 29 | 1 | 58 EEG + 2 EMG + 2 EOG | 100 Hz | Nap (1–2 h) | 30 s | Collaborator | [20] | 3 |
| CCSHS | Human | Healthy (adolescents, 17-18) | 20 | 20 | — | 2 EEG + EMG + EOG + ECG | 128 Hz | Overnight | 30 s | Open | [19,21] | 4 |
| CHAT | Human | OSA (children, 5-10) | 20 | 20 | — | 8 EEG + EMG + EOG + ECG | >200 Hz | Overnight | 30 s | Open | [19,22] | 4 |
| ***Other species*** | | | | | | | | | | | | |
| Jackdaws | Bird | Healthy + SD | 9 | 27 (9×3 days) | — | 32 EEG + accelerometer | 250 Hz | 24 h | 4 s | Collaborator | [23] | 4,6 |
| Ephy animals | Lizard | Aregntine tegu (Salvator merianae) | 1 | 1 | — | 5 DVRA+2 EMG+2 ECG+4 EOG+7 LFP+2 olfactory bulb | 250 Hz | 20 h | — | Collaborator | — | 6 |
| | Rat (wild) | Wild Black Rat (Rattus rattus) (telemetry) | 1 | 1 | — | 4 EEG + 1 EMG + 1 ECG | 256 Hz | 24 h | 4 s | Collaborator | — | S7 |

*P, parietal; F, frontal; O, occipital; WT, wild-type; SD, sleep deprivation; OSA, obstructive sleep apnea; LFP, local field potential; SDS, sleep-disorders suspicion. Source "Open" denotes a publicly available dataset; "In-house" denotes data acquired in the Boccara laboratory. "Scoring epoch" is the epoch length of the manual (ground truth) annotation.*

**Supplementary Table 2. Per-recording classification performance of Nyx relative to manual annotation in rodents.**

| Category | MF1 | Accuracy | Cohen's κ | WAKE F1 | REM F1 | NREM F1 | WAKE Prec | REM Prec | NREM Prec | WAKE Recall | REM Recall | NREM Recall | N |
|---|---|---|---|---|---|---|---|---|---|---|---|---|---|
| ***Baseline recordings*** | | | | | | | | | | | | | |
| **All** | 0.874 ± 0.043 | 0.905 ± 0.034 | 0.832 ± 0.060 | 0.923 ± 0.044 | 0.792 ± 0.094 | 0.906 ± 0.035 | 0.926 ± 0.057 | 0.752 ± 0.152 | 0.919 ± 0.046 | 0.924 ± 0.060 | 0.868 ± 0.079 | 0.899 ± 0.063 | 179 |
| **Rat** | 0.864 ± 0.045 | 0.895 ± 0.036 | 0.816 ± 0.065 | 0.908 ± 0.048 | 0.783 ± 0.093 | 0.900 ± 0.038 | 0.917 ± 0.063 | 0.740 ± 0.155 | 0.914 ± 0.044 | 0.904 ± 0.065 | 0.865 ± 0.081 | 0.891 ± 0.070 | 103 |
| **Mouse** | 0.888 ± 0.037 | 0.920 ± 0.025 | 0.853 ± 0.044 | 0.943 ± 0.026 | 0.804 ± 0.094 | 0.915 ± 0.028 | 0.938 ± 0.045 | 0.767 ± 0.147 | 0.926 ± 0.049 | 0.951 ± 0.040 | 0.872 ± 0.077 | 0.909 ± 0.051 | 76 |
| ***Manipulation recordings*** | | | | | | | | | | | | | |
| **All** | 0.879 ± 0.042 | 0.906 ± 0.036 | 0.833 ± 0.062 | 0.919 ± 0.044 | 0.815 ± 0.086 | 0.904 ± 0.033 | 0.944 ± 0.059 | 0.815 ± 0.150 | 0.888 ± 0.051 | 0.901 ± 0.065 | 0.841 ± 0.080 | 0.927 ± 0.065 | 171 |
| **Rat** | 0.877 ± 0.040 | 0.895 ± 0.034 | 0.818 ± 0.059 | 0.905 ± 0.039 | 0.830 ± 0.081 | 0.897 ± 0.034 | 0.918 ± 0.058 | 0.854 ± 0.139 | 0.895 ± 0.052 | 0.897 ± 0.063 | 0.827 ± 0.077 | 0.906 ± 0.066 | 99 |
| **Mouse** | 0.882 ± 0.045 | 0.922 ± 0.033 | 0.854 ± 0.061 | 0.939 ± 0.044 | 0.795 ± 0.089 | 0.913 ± 0.029 | 0.980 ± 0.037 | 0.760 ± 0.147 | 0.879 ± 0.048 | 0.905 ± 0.067 | 0.861 ± 0.080 | 0.955 ± 0.050 | 72 |

*Values are mean ± standard deviation across recordings. MF1, macro F1 (unweighted mean of per-stage F1 scores). All metrics were computed per recording at the epoch level. N indicates the number of recordings.*

**Supplementary Table 3. Per-dataset classification performance of Nyx in rodents.**

| Dataset | Species | N | MF1 | Accuracy | Cohen's κ | WAKE F1 | REM F1 | NREM F1 | WAKE Prec | REM Prec | NREM Prec | WAKE Recall | REM Recall | NREM Recall |
|---|---|---|---|---|---|---|---|---|---|---|---|---|---|---|
| ***Baseline recordings*** | | | | | | | | | | | | | | |
| **Boccara DSI** | Rat | 7 | 0.854 ± 0.051 | 0.857 ± 0.048 | 0.743 ± 0.077 | 0.802 ± 0.066 | 0.875 ± 0.072 | 0.884 ± 0.037 | 0.873 ± 0.078 | 0.851 ± 0.129 | 0.858 ± 0.066 | 0.753 ± 0.110 | 0.914 ± 0.045 | 0.916 ± 0.048 |
| **Ellen Dash** | Rat | 36 | 0.826 ± 0.046 | 0.870 ± 0.036 | 0.770 ± 0.065 | 0.886 ± 0.038 | 0.715 ± 0.111 | 0.877 ± 0.046 | 0.891 ± 0.079 | 0.605 ± 0.142 | 0.914 ± 0.032 | 0.887 ± 0.043 | 0.909 ± 0.063 | 0.848 ± 0.086 |
| **Gulledge Baseline** | Rat | 30 | 0.887 ± 0.026 | 0.911 ± 0.021 | 0.847 ± 0.036 | 0.930 ± 0.021 | 0.819 ± 0.060 | 0.911 ± 0.023 | 0.930 ± 0.044 | 0.815 ± 0.108 | 0.921 ± 0.044 | 0.933 ± 0.037 | 0.836 ± 0.074 | 0.906 ± 0.053 |
| **Gulledge Saline** | Rat | 30 | 0.888 ± 0.017 | 0.916 ± 0.016 | 0.856 ± 0.028 | 0.937 ± 0.020 | 0.806 ± 0.040 | 0.920 ± 0.019 | 0.944 ± 0.029 | 0.802 ± 0.100 | 0.921 ± 0.045 | 0.931 ± 0.034 | 0.828 ± 0.086 | 0.921 ± 0.034 |
| **Oxford Opto Baseline** | Mouse | 11 | 0.914 ± 0.016 | 0.928 ± 0.019 | 0.869 ± 0.034 | 0.938 ± 0.021 | 0.882 ± 0.035 | 0.921 ± 0.023 | 0.931 ± 0.054 | 0.865 ± 0.078 | 0.940 ± 0.044 | 0.949 ± 0.041 | 0.905 ± 0.029 | 0.909 ± 0.064 |
| **Oxford SD Baseline** | Mouse | 6 | 0.892 ± 0.010 | 0.909 ± 0.010 | 0.835 ± 0.017 | 0.891 ± 0.019 | 0.855 ± 0.033 | 0.929 ± 0.010 | 0.885 ± 0.039 | 0.798 ± 0.075 | 0.945 ± 0.034 | 0.900 ± 0.046 | 0.928 ± 0.039 | 0.915 ± 0.028 |
| **Oxford Test** | Mouse | 6 | 0.928 ± 0.013 | 0.944 ± 0.009 | 0.902 ± 0.016 | 0.957 ± 0.009 | 0.888 ± 0.027 | 0.941 ± 0.011 | 0.979 ± 0.017 | 0.893 ± 0.070 | 0.922 ± 0.035 | 0.936 ± 0.027 | 0.890 ± 0.064 | 0.962 ± 0.031 |
| **SIESTA TSE** | Mouse | 3 | 0.876 ± 0.057 | 0.929 ± 0.033 | 0.870 ± 0.059 | 0.950 ± 0.026 | 0.750 ± 0.116 | 0.929 ± 0.036 | 0.932 ± 0.058 | 0.652 ± 0.170 | 0.966 ± 0.027 | 0.970 ± 0.013 | 0.915 ± 0.033 | 0.897 ± 0.073 |
| **SIESTA WT** | Mouse | 10 | 0.853 ± 0.050 | 0.930 ± 0.028 | 0.863 ± 0.058 | 0.961 ± 0.021 | 0.688 ± 0.107 | 0.911 ± 0.044 | 0.965 ± 0.026 | 0.615 ± 0.192 | 0.932 ± 0.062 | 0.959 ± 0.042 | 0.846 ± 0.090 | 0.895 ± 0.063 |
| **Sippel Morris** | Mouse | 40 | 0.883 ± 0.032 | 0.913 ± 0.025 | 0.840 ± 0.041 | 0.946 ± 0.021 | 0.796 ± 0.078 | 0.908 ± 0.024 | 0.936 ± 0.042 | 0.764 ± 0.128 | 0.915 ± 0.050 | 0.958 ± 0.036 | 0.856 ± 0.085 | 0.905 ± 0.046 |
| ***Manipulation recordings*** | | | | | | | | | | | | | | |
| **Gulledge Oxy** | Rat | 99 | 0.877 ± 0.040 | 0.895 ± 0.034 | 0.818 ± 0.059 | 0.905 ± 0.039 | 0.830 ± 0.081 | 0.897 ± 0.034 | 0.918 ± 0.058 | 0.854 ± 0.139 | 0.895 ± 0.052 | 0.897 ± 0.063 | 0.827 ± 0.077 | 0.906 ± 0.066 |
| **Oxford Opto** | Mouse | 11 | 0.878 ± 0.046 | 0.908 ± 0.038 | 0.831 ± 0.068 | 0.921 ± 0.041 | 0.809 ± 0.094 | 0.903 ± 0.038 | 0.973 ± 0.020 | 0.807 ± 0.165 | 0.858 ± 0.066 | 0.877 ± 0.072 | 0.843 ± 0.082 | 0.957 ± 0.020 |
| **Oxford SD** | Mouse | 6 | 0.920 ± 0.006 | 0.943 ± 0.008 | 0.892 ± 0.016 | 0.964 ± 0.008 | 0.878 ± 0.017 | 0.919 ± 0.013 | 0.985 ± 0.016 | 0.847 ± 0.068 | 0.891 ± 0.027 | 0.943 ± 0.011 | 0.917 ± 0.048 | 0.950 ± 0.031 |
| **SIESTA APP-PS1** | Mouse | 5 | 0.790 ± 0.080 | 0.875 ± 0.075 | 0.756 ± 0.142 | 0.871 ± 0.129 | 0.622 ± 0.103 | 0.878 ± 0.051 | 0.941 ± 0.070 | 0.525 ± 0.144 | 0.883 ± 0.083 | 0.841 ± 0.208 | 0.815 ± 0.114 | 0.885 ± 0.103 |
| **SIESTA NmsVgat** | Mouse | 4 | 0.890 ± 0.033 | 0.934 ± 0.023 | 0.881 ± 0.040 | 0.954 ± 0.019 | 0.782 ± 0.063 | 0.934 ± 0.022 | 0.980 ± 0.020 | 0.702 ± 0.100 | 0.927 ± 0.031 | 0.931 ± 0.035 | 0.891 ± 0.037 | 0.943 ± 0.033 |
| **SIESTA SCN1a** | Mouse | 6 | 0.861 ± 0.033 | 0.911 ± 0.026 | 0.836 ± 0.047 | 0.931 ± 0.030 | 0.748 ± 0.107 | 0.903 ± 0.027 | 0.947 ± 0.069 | 0.682 ± 0.163 | 0.901 ± 0.054 | 0.920 ± 0.047 | 0.853 ± 0.065 | 0.911 ± 0.067 |

| Dataset | Species | N | MF1 | Accuracy | Cohen's κ | WAKE F1 | REM F1 | NREM F1 | WAKE Prec | REM Prec | NREM Prec | WAKE Recall | REM Recall | NREM Recall |
|---|---|---|---|---|---|---|---|---|---|---|---|---|---|---|
| **Sippel Morris KO** | Mouse | 40 | 0.892 ± 0.028 | 0.928 ± 0.020 | 0.867 ± 0.035 | 0.948 ± 0.019 | 0.808 ± 0.061 | 0.919 ± 0.020 | 0.990 ± 0.027 | 0.781 ± 0.122 | 0.874 ± 0.037 | 0.910 ± 0.030 | 0.861 ± 0.081 | 0.972 ± 0.036 |

*Values are mean ± standard deviation across recordings within each dataset. Same column structure as Supplementary Table 2; rows are individual datasets used in the rodent analysis, separated into baseline recordings (used for the main per-dataset MF1 comparison) and manipulation recordings. MF1, macro F1 (unweighted mean of per-stage F1 scores). N indicates the number of recordings per dataset.*

**Supplementary Table 4. Per-dataset agreement of Nyx and Somnotate with manual scoring across the rodent benchmark.**

| Dataset | N | Method | Accuracy (M ± SD) | Cohen’s κ (M ± SD) | MF1 (M ± SD) | WAKE F1 (M ± SD) | NREM F1 (M ± SD) | REM F1 (M ± SD) |
|---|---|---|---|---|---|---|---|---|
| **Gulledge Saline** | 30 | *Nyx* | 0.916 ± 0.016 | 0.856 ± 0.028 | 0.888 ± 0.017 | 0.937 ± 0.020 | 0.920 ± 0.019 | 0.806 ± 0.040 |
| | | *Somnotate* | 0.929 ± 0.021 | 0.877 ± 0.036 | 0.905 ± 0.028 | 0.944 ± 0.019 | 0.930 ± 0.024 | 0.841 ± 0.058 |
| **Gulledge Baseline** | 30 | *Nyx* | 0.911 ± 0.021 | 0.847 ± 0.036 | 0.887 ± 0.026 | 0.930 ± 0.021 | 0.911 ± 0.023 | 0.819 ± 0.060 |
| | | *Somnotate* | 0.930 ± 0.023 | 0.877 ± 0.040 | 0.889 ± 0.044 | 0.950 ± 0.023 | 0.935 ± 0.016 | 0.783 ± 0.101 |
| **Sippel Morris WT** | 40 | *Nyx* | 0.913 ± 0.025 | 0.840 ± 0.041 | 0.883 ± 0.032 | 0.946 ± 0.021 | 0.908 ± 0.024 | 0.796 ± 0.078 |
| | | *Somnotate* | 0.842 ± 0.036 | 0.729 ± 0.057 | 0.757 ± 0.033 | 0.940 ± 0.032 | 0.812 ± 0.042 | 0.518 ± 0.053 |
| **Gulledge Oxy** | 99 | *Nyx* | 0.895 ± 0.034 | 0.818 ± 0.059 | 0.877 ± 0.040 | 0.905 ± 0.039 | 0.897 ± 0.034 | 0.830 ± 0.081 |
| | | *Somnotate* | 0.896 ± 0.040 | 0.819 ± 0.068 | 0.853 ± 0.056 | 0.910 ± 0.042 | 0.909 ± 0.037 | 0.741 ± 0.123 |
| **Sippel Morris KO** | 40 | *Nyx* | 0.928 ± 0.020 | 0.867 ± 0.035 | 0.892 ± 0.028 | 0.948 ± 0.019 | 0.919 ± 0.020 | 0.808 ± 0.061 |
| | | *Somnotate* | 0.871 ± 0.032 | 0.769 ± 0.058 | 0.797 ± 0.040 | 0.948 ± 0.025 | 0.836 ± 0.048 | 0.605 ± 0.058 |
| **Overall (pooled)** | 239 | *Nyx* | 0.908 ± 0.030 | 0.838 ± 0.050 | 0.883 ± 0.033 | 0.926 ± 0.035 | 0.907 ± 0.029 | 0.816 ± 0.071 |
| | | *Somnotate* | 0.891 ± 0.045 | 0.810 ± 0.077 | 0.839 ± 0.067 | 0.931 ± 0.038 | 0.887 ± 0.058 | 0.699 ± 0.142 |

*Values are mean ± standard deviation across recordings of accuracy, Cohen’s κ, macro-F1 (MF1) and per-stage F1 (WAKE, NREM, REM) for Nyx and Somnotate relative to manual expert scoring, per dataset and pooled across all five datasets (bottom). Somnotate was trained exclusively on its original mouse dataset (six recordings from the Oxford Mouse Benchmark) and applied without retraining. N, number of paired recordings per dataset.*

**Supplementary Table 5. Agreement between manual and Nyx automated scoring for derived sleep variables in rodents.**

| Variable | Manual (M ± SD) | Automatic (M ± SD) | Bias (M ± SD) | LoA [lo, hi] | Wilcoxon p | N_rec (N_ani) | ICC_rec [95% CI] | ICC_ani [95% CI] | Pearson r |
|---|---|---|---|---|---|---|---|---|---|
| ***Baseline recordings*** | | | | | | | | | |
| **Total sleep time (%)** | 51.33 ± 7.74 | 50.95 ± 8.66 | -0.38 ± 4.07 | [-8.36, 7.59] | 0.157 | 179 (67) | 0.88 [0.84, 0.91] | 0.93 [0.89, 0.96] | 0.883 |
| **REM/NREM ratio** | 0.17 ± 0.05 | 0.21 ± 0.06 | +0.04 ± 0.06 | [-0.08, 0.15] | <0.001 | 179 (67) | 0.36 [0.12, 0.54] | 0.38 [0.05, 0.61] | 0.448 |
| **Microarousals (count/h)** | 10.86 ± 5.56 | 8.70 ± 4.79 | -2.17 ± 5.45 | [-12.86, 8.52] | <0.001 | 119 (53) | 0.41 [0.23, 0.56] | 0.51 [0.28, 0.68] | 0.452 |
| **Theta peak freq. REM (Hz)** | 6.93 ± 0.34 | 6.87 ± 0.38 | -0.06 ± 0.13 | [-0.33, 0.20] | <0.001 | 179 (67) | 0.92 [0.86, 0.95] | 0.93 [0.85, 0.96] | 0.937 |
| **Delta power NREM (dB)** | 42.40 ± 6.41 | 42.44 ± 6.47 | +0.05 ± 0.44 | [-0.82, 0.92] | 0.244 | 165 (57) | 0.997 [0.996, 0.998] | 0.998 [0.997, 0.999] | 0.998 |
| ***Manipulation recordings*** | | | | | | | | | |
| **Total sleep time (%)** | 50.67 ± 8.64 | 52.59 ± 8.54 | +1.92 ± 4.81 | [-7.51, 11.34] | <0.001 | 171 (53) | 0.82 [0.73, 0.88] | 0.81 [0.53, 0.91] | 0.843 |
| **REM/NREM ratio** | 0.19 ± 0.06 | 0.19 ± 0.08 | 0.00 ± 0.07 | [-0.13, 0.14] | 0.135 | 171 (53) | 0.45 [0.33, 0.57] | 0.70 [0.53, 0.81] | 0.479 |
| **Microarousals (count/h)** | 12.32 ± 5.20 | 5.28 ± 4.62 | -7.03 ± 6.16 | [-19.11, 5.04] | <0.001 | 71 (41) | 0.11 [-0.07, 0.30] | 0.16 [-0.09, 0.42] | 0.218 |
| **Theta peak freq. REM (Hz)** | 6.90 ± 0.47 | 6.88 ± 0.50 | -0.02 ± 0.17 | [-0.36, 0.31] | 0.397 | 171 (53) | 0.94 [0.92, 0.95] | 0.95 [0.91, 0.97] | 0.941 |
| **Delta power NREM (dB)** | 39.58 ± 6.15 | 39.51 ± 6.11 | -0.06 ± 0.68 | [-1.40, 1.27] | <0.001 | 169 (51) | 0.993 [0.992, 0.995] | 0.999 [0.998, 0.999] | 0.994 |

*Values are mean ± standard deviation across recordings. Bias is defined as automatic - manual; positive bias indicates that the automatic method overestimates relative to manual. Limits of agreement (LoA) are bias ± 1.96·SD of the paired differences. Wilcoxon signed-rank tests the null of zero median paired difference; values below 0.001 are reported as <0.001. ICCs are ICC(A,1), two-way mixed-effects, absolute-agreement, single measures (Shrout & Fleiss ICC(2,1)); ICC_rec is computed treating recordings as targets (N_rec) and ICC_ani treating animals as targets (N_ani), the latter providing a clustering-conservative comparison. Brackets show 95% confidence intervals. N differs across variables because Microarousals were scored on a subset of recordings and some recordings lacked the channels needed for delta-power estimation.*

**Supplementary Table 6. Per-recording classification performance of Nyx relative to manual annotation in humans, across three scoring schemes.**

| Analysis | MF1 | Accuracy | Cohen's κ | F1 per stage | Precision per stage | Recall per stage | N |
|---|---|---|---|---|---|---|---|
| ***3-stage scoring (WAKE / REM / NREM)*** | | | | | | | |
| **All** | 0.815 ± 0.089 | 0.869 ± 0.066 | 0.737 ± 0.120 | 0.772 / 0.780 / 0.903 | 0.802 / 0.770 / 0.910 | 0.783 / 0.829 / 0.904 | 179 |
| ***4-stage scoring (WAKE / REM / N1+N2 / N3)*** | | | | | | | |
| **All** | 0.738 ± 0.117 | 0.788 ± 0.096 | 0.666 ± 0.138 | 0.772 / 0.780 / 0.794 / 0.661 | 0.802 / 0.770 / 0.841 / 0.624 | 0.783 / 0.829 / 0.766 / 0.856 | 179 |
| ***5-stage scoring (WAKE / REM / N1 / N2 / N3)*** | | | | | | | |
| **All** | 0.621 ± 0.111 | 0.719 ± 0.106 | 0.605 ± 0.135 | 0.772 / 0.780 / 0.271 / 0.735 / 0.661 | 0.802 / 0.770 / 0.245 / 0.827 / 0.624 | 0.783 / 0.829 / 0.368 / 0.689 / 0.856 | 179 |

*Values are mean ± standard deviation across recordings. Each scoring scheme collapses the AASM stages differently: 3-stage merges N1, N2, and N3 into NREM; 4-stage merges N1 and N2 into a single stage; 5-stage retains the full AASM resolution. Per-stage F1, precision, and recall are slash-separated in the stage order shown in each section header. MF1, macro F1 (unweighted mean of per-stage F1 scores). Recordings flagged as poor signal quality were excluded; the number excluded is reported in the Methods. N reflects the recordings retained.*

**Supplementary Table 7. Per-dataset classification performance of Nyx in humans, across three scoring schemes.**

| Dataset | MF1 | Accuracy | Cohen's κ | F1 per stage | Precision per stage | Recall per stage | N |
|---|---|---|---|---|---|---|---|
| *3-stage scoring (WAKE / REM / NREM)* | | | | | | | |
| **ANPHY** | 0.836 ± 0.055 | 0.893 ± 0.033 | 0.770 ± 0.070 | 0.811 / 0.771 / 0.926 | 0.837 / 0.719 / 0.939 | 0.797 / 0.858 / 0.915 | 29 |
| **DOD-H** | 0.809 ± 0.080 | 0.875 ± 0.049 | 0.735 ± 0.111 | 0.708 / 0.807 / 0.913 | 0.653 / 0.835 / 0.932 | 0.815 / 0.798 / 0.898 | 22 |
| **DOD-O** | 0.822 ± 0.070 | 0.879 ± 0.050 | 0.751 ± 0.095 | 0.755 / 0.796 / 0.911 | 0.775 / 0.790 / 0.915 | 0.768 / 0.833 / 0.912 | 49 |
| **MESA** | 0.798 ± 0.119 | 0.851 ± 0.086 | 0.726 ± 0.155 | 0.809 / 0.704 / 0.875 | 0.853 / 0.698 / 0.880 | 0.811 / 0.761 / 0.884 | 51 |
| **SleePing** | 0.815 ± 0.092 | 0.856 ± 0.077 | 0.700 ± 0.129 | 0.742 / 0.743 / 0.909 | 0.836 / 0.740 / 0.910 | 0.722 / 0.801 / 0.916 | 28 |
| *4-stage scoring (WAKE / REM / N1+N2 / N3)* | | | | | | | |
| **ANPHY** | 0.811 ± 0.056 | 0.829 ± 0.050 | 0.738 ± 0.078 | 0.811 / 0.771 / 0.833 / 0.827 | 0.837 / 0.719 / 0.857 / 0.828 | 0.797 / 0.858 / 0.817 / 0.859 | 29 |
| **DOD-H** | 0.775 ± 0.077 | 0.801 ± 0.059 | 0.689 ± 0.103 | 0.708 / 0.807 / 0.814 / 0.768 | 0.653 / 0.835 / 0.855 / 0.774 | 0.815 / 0.798 / 0.786 / 0.829 | 22 |
| **DOD-O** | 0.741 ± 0.091 | 0.780 ± 0.074 | 0.651 ± 0.122 | 0.755 / 0.796 / 0.791 / 0.620 | 0.775 / 0.790 / 0.855 / 0.551 | 0.768 / 0.833 / 0.748 / 0.886 | 49 |
| **MESA** | 0.667 ± 0.135 | 0.764 ± 0.120 | 0.628 ± 0.165 | 0.809 / 0.704 / 0.739 / 0.359 | 0.853 / 0.698 / 0.783 / 0.329 | 0.811 / 0.761 / 0.718 / 0.606 | 51 |
| **SleePing** | 0.760 ± 0.130 | 0.794 ± 0.125 | 0.670 ± 0.156 | 0.742 / 0.743 / 0.812 / 0.693 | 0.836 / 0.740 / 0.865 / 0.636 | 0.722 / 0.801 / 0.787 / 0.877 | 28 |
| *5-stage scoring (WAKE / REM / N1 / N2 / N3)* | | | | | | | |
| **ANPHY** | 0.701 ± 0.060 | 0.770 ± 0.061 | 0.685 ± 0.081 | 0.811 / 0.771 / 0.286 / 0.811 / 0.827 | 0.837 / 0.719 / 0.295 / 0.836 / 0.828 | 0.797 / 0.858 / 0.302 / 0.796 / 0.859 | 29 |
| **DOD-H** | 0.657 ± 0.074 | 0.741 ± 0.062 | 0.637 ± 0.095 | 0.708 / 0.807 / 0.220 / 0.784 / 0.768 | 0.653 / 0.835 / 0.170 / 0.899 / 0.774 | 0.815 / 0.798 / 0.357 / 0.707 / 0.829 | 22 |
| **DOD-O** | 0.621 ± 0.084 | 0.720 ± 0.084 | 0.599 ± 0.117 | 0.755 / 0.796 / 0.198 / 0.743 / 0.620 | 0.775 / 0.790 / 0.168 / 0.867 / 0.551 | 0.768 / 0.833 / 0.302 / 0.666 / 0.886 | 49 |
| **MESA** | 0.534 ± 0.115 | 0.675 ± 0.132 | 0.540 ± 0.155 | 0.809 / 0.704 / 0.147 / 0.638 / 0.359 | 0.853 / 0.698 / 0.138 / 0.705 / 0.329 | 0.811 / 0.761 / 0.210 / 0.628 / 0.606 | 51 |
| **SleePing** | 0.666 ± 0.108 | 0.726 ± 0.127 | 0.623 ± 0.146 | 0.742 / 0.743 / 0.436 / 0.757 / 0.693 | 0.836 / 0.740 / 0.387 / 0.880 / 0.636 | 0.722 / 0.801 / 0.558 / 0.690 / 0.877 | 28 |

*MF1, Accuracy, and Cohen's κ are mean ± standard deviation across recordings. Per-stage F1, precision, and recall are slash-separated means in the stage order shown in each section header (full SDs available on request). Same column structure as Supplementary Table 5; rows are individual human datasets shown under each of the three scoring schemes. MF1, macro F1 (unweighted mean of per-stage F1 scores). Recordings flagged as poor signal quality were excluded. N indicates the number of recordings retained per dataset.*

#### Supplementary Table 8. Per-dataset comparison of Nyx and YASA on human recordings across stage schemes.

##### *8a. Per-dataset agreement of Nyx and YASA with manual scoring on human recordings across stage schemes.*

| Dataset | N | Method | Accuracy (M ± SD) | Cohen's κ (M ± SD) | MF1 (M ± SD) | F1 per stage |
|---|---|---|---|---|---|---|
| ***3-stage scoring (WAKE / REM / NREM)*** | | | | | | |
| ANPHY | *29* | *Nyx* | 0.893 ± 0.033 | 0.770 ± 0.070 | 0.836 ± 0.055 | 0.811 / 0.771 / 0.926 |
| | | *YASA* | 0.814 ± 0.087 | 0.654 ± 0.147 | 0.768 ± 0.118 | 0.655 / 0.781 / 0.868 |
| SleePing | 28 | *Nyx* | 0.856 ± 0.077 | 0.700 ± 0.129 | 0.815 ± 0.092 | 0.742 / 0.743 / 0.909 |
| | | *YASA* | 0.880 ± 0.068 | 0.754 ± 0.135 | 0.867 ± 0.077 | 0.849 / 0.832 / 0.911 |
| ***4-stage scoring (WAKE / REM / N1+N2 / N3)*** | | | | | | |
| ANPHY | *29* | *Nyx* | 0.829 ± 0.050 | 0.738 ± 0.078 | 0.811 ± 0.056 | 0.811 / 0.771 / 0.833 / 0.827 |
| | | *YASA* | 0.772 ± 0.094 | 0.665 ± 0.132 | 0.764 ± 0.105 | 0.655 / 0.781 / 0.787 / 0.834 |
| SleePing | *28* | *Nyx* | 0.794 ± 0.125 | 0.670 ± 0.156 | 0.760 ± 0.130 | 0.742 / 0.743 / 0.812 / 0.693 |
| | | *YASA* | 0.830 ± 0.083 | 0.720 ± 0.133 | 0.825 ± 0.101 | 0.849 / 0.832 / 0.834 / 0.770 |
| ***5-stage scoring (WAKE / REM / N1 / N2 / N3)*** | | | | | | |
| ANPHY | *29* | *Nyx* | 0.770 ± 0.061 | 0.685 ± 0.081 | 0.701 ± 0.060 | 0.811 / 0.771 / 0.286 / 0.811 / 0.827 |
| | | *YASA* | 0.751 ± 0.092 | 0.657 ± 0.122 | 0.650 ± 0.100 | 0.655 / 0.781 / 0.151 / 0.827 / 0.834 |
| SleePing | 28 | *Nyx* | 0.726 ± 0.127 | 0.623 ± 0.146 | 0.666 ± 0.108 | 0.742 / 0.743 / 0.436 / 0.757 / 0.693 |
| | | *YASA* | 0.792 ± 0.099 | 0.701 ± 0.136 | 0.734 ± 0.120 | 0.849 / 0.832 / 0.478 / 0.826 / 0.770 |

*Values are Mean ± standard deviation across recordings of accuracy, Cohen's κ, macro-F1 (MF1) and per-stage F1 for Nyx and YASA relative to manual expert scoring, on ANPHY and SleePing, evaluated under 3-, 4- and 5-stage AASM schemes. Under the 3-stage scheme, NREM is a single stage; under the 4-stage scheme, N1 and N2 are merged. YASA was applied without retraining; neither ANPHY nor SleePing was part of YASA's original training data. N, number of paired recordings per dataset.*

*8b. Statistical comparison of Nyx and YASA on human recordings across stage schemes.*

| Dataset | N | Median MF1 Nyx | Median MF1 YASA | Median diff (Nyx − YASA) | W | p | r | Recordings Nyx > YASA |
|---|---|---|---|---|---|---|---|---|
| ***3-stage scoring (WAKE / REM / NREM)*** | | | | | | | | |
| ANPHY | 29 | 0.848 | 0.797 | +0.064 | 65.0 | **<0.001** | +0.70 | 22 / 29 |
| SleePing | 28 | 0.824 | 0.876 | −0.046 | 64.0 | **<0.001** | −0.69 | 7 / 28 |
| *Pooled* | 57 | 0.843 | 0.826 | +0.001 | 809.0 | 0.889 | +0.02 | 29 / 57 |
| ***4-stage scoring (WAKE / REM / N1+N2 / N3)*** | | | | | | | | |
| ANPHY | 29 | 0.824 | 0.778 | +0.028 | 97.0 | **0.008** | +0.55 | 21 / 29 |
| SleePing | 28 | 0.788 | 0.859 | −0.055 | 57.0 | **<0.001** | −0.72 | 7 / 28 |
| *Pooled* | 57 | 0.796 | 0.820 | −0.003 | 738.0 | 0.482 | −0.11 | 28 / 57 |
| ***5-stage scoring (WAKE / REM / N1 / N2 / N3)*** | | | | | | | | |
| ANPHY | 29 | 0.697 | 0.658 | +0.047 | 99.0 | **0.009** | +0.55 | 19 / 29 |
| SleePing | 28 | 0.683 | 0.757 | −0.061 | 89.0 | **0.008** | −0.56 | 9 / 28 |
| *Pooled* | 57 | 0.692 | 0.712 | −0.009 | 772.0 | 0.665 | −0.07 | 28 / 57 |

*Per-recording macro-F1 (MF1) values for Nyx and YASA relative to manual expert scoring, on ANPHY (n = 29 recordings) and SleePing (n = 28 recordings), under 3-, 4-, and 5-stage AASM schemes. Median difference reported as Nyx − YASA (positive = Nyx higher). Comparisons made with two-sided Wilcoxon signed-rank tests on per-recording MF1; effect size reported as matched-pairs rank-biserial correlation (r; positive = Nyx higher). Significant p-values ($p < 0.05$) are shown in bold. The final column indicates the number of recordings on which Nyx exceeded YASA out of the total N. YASA was applied without retraining to datasets outside its original training set. Neither ANPHY nor SleePing was part of YASA's training data.*

**Supplementary Table 9. Agreement between manual and Nyx automated scoring for derived sleep variables in humans.**

| Variable | Manual (M ± SD) | Automatic (M ± SD) | Bias (M ± SD) | LoA [lo, hi] | Wilcoxon p | N_rec | ICC_rec [95% CI] | Pearson r |
|---|---|---|---|---|---|---|---|---|
| **Total sleep time (%)** | 74.93 ± 15.16 | 76.40 ± 13.77 | +1.47 ± 8.40 | [-15.00, 17.94] | 0.075 | 177 | 0.83 [0.77, 0.87] | 0.835 |
| **REM/NREM ratio** | 0.21 ± 0.12 | 0.23 ± 0.13 | +0.03 ± 0.11 | [-0.18, 0.23] | 0.001 | 177 | 0.65 [0.55, 0.73] | 0.665 |
| **Sleep Onset Latency (min)** | 35.10 ± 43.89 | 25.53 ± 38.47 | -9.58 ± 34.02 | [-76.26, 57.10] | <0.001 | 177 | 0.64 [0.54, 0.73] | 0.666 |
| **WASO (min)** | 74.71 ± 69.08 | 79.51 ± 65.33 | +4.80 ± 46.17 | [-85.69, 95.28] | 0.018 | 177 | 0.76 [0.69, 0.82] | 0.765 |
| **REM Latency (min)** | 120.66 ± 75.42 | 85.53 ± 74.67 | -35.13 ± 87.98 | [-207.58, 137.31] | <0.001 | 156 | 0.28 [0.12, 0.43] | 0.313 |
| **Sigma power N2 (dB)** | 21.65 ± 9.35 | 21.72 ± 9.42 | +0.07 ± 0.45 | [-0.82, 0.96] | 0.059 | 172 | 0.999 [0.998, 0.999] | 0.999 |
| **Delta power N3 (dB)** | 38.17 ± 11.32 | 37.51 ± 11.27 | -0.66 ± 0.82 | [-2.27, 0.94] | <0.001 | 148 | 0.999 [0.98, 0.999] | 0.997 |
| **Theta power REM (dB)** | 23.66 ± 10.89 | 23.57 ± 10.84 | -0.09 ± 0.51 | [-1.08, 0.90] | <0.001 | 153 | 0.999 [0.998, 0.999] | 0.999 |

*Values are mean ± standard deviation across recordings. Bias is defined as automatic - manual; positive bias indicates that the automatic method overestimates relative to manual. Limits of agreement (LoA) are bias ± 1.96·SD of the paired differences. Wilcoxon signed-rank tests the null of zero median paired difference; values below 0.001 are reported as <0.001. ICCs are ICC(A,1), two-way mixed-effects, absolute-agreement, single measures (Shrout & Fleiss ICC(2,1)); ICC_rec is computed treating recordings as targets (N_rec). Brackets show 95% confidence intervals. N differs across variables some recordings lacked specific stages for variable estimation.*

**Supplementary Table 10. Lifespan and cross-species generalization of Nyx.**

| Species | Group | Scheme | N individuals | N recordings | MF1 | F1 per stage | Precision per stage | Recall per stage |
|---|---|---|---|---|---|---|---|---|
| ***Rodents lifespan (WAKE / REM / NREM)*** | | | | | | | | |
| **Rat** | P12 | 3 stages | 2 | 2 | 0.83 ± 0.03 | 0.70/0.88/0.91 | 0.61/0.93/0.94 | 0.82/0.83/0.88 |
| **Rat** | P18 | 3 stages | 2 | 2 | 0.94 ± 0.01 | 0.91/0.95/0.95 | 0.88/0.94/0.97 | 0.95/0.96/0.93 |
| **Rat** | P25 | 3 stages | 2 | 2 | 0.89 ± 0.03 | 0.84/0.89/0.95 | 0.77/0.88/0.98 | 0.92/0.90/0.93 |
| **Rat** | Adult (P60-P200) | 3 stages | 2 | 4 | 0.86 ± 0.01 | 0.80/0.82/0.95 | 0.90/0.79/0.96 | 0.75/0.87/0.94 |
| **Mouse** | Aged (~P420) | 3 stages | 2 | 2 | 0.83 ± 0.03 | 0.87/0.86/0.76 | 0.97/0.82/0.66 | 0.79/0.91/0.92 |
| ***Humans lifespan*** | | | | | | | | |
| ***3 stages scoring (WAKE / REM / NREM)*** | | | | | | | | |
| **Human** | 5-7.5 y | 3 stages | 10 | 10 | 0.81 ± 0.07 | 0.74/0.77/0.92 | 0.75/0.72/0.96 | 0.80/0.86/0.88 |
| **Human** | 7.5-10 y | 3 stages | 10 | 10 | 0.82 ± 0.07 | 0.78/0.78/0.91 | 0.75/0.71/0.95 | 0.87/0.89/0.87 |
| **Human** | 17 y | 3 stages | 15 | 15 | 0.87 ± 0.03 | 0.87/0.83/0.91 | 0.84/0.82/0.94 | 0.92/0.85/0.88 |
| **Human** | 18 y | 3 stages | 5 | 5 | 0.86 ± 0.04 | 0.88/0.79/0.90 | 0.88/0.74/0.92 | 0.89/0.86/0.89 |
| **Human** | 70-80 y | 3 stages | 9 | 9 | 0.80 ± 0.14 | 0.75/0.76/0.90 | 0.87/0.80/0.86 | 0.70/0.74/0.94 |
| **Human** | 80-90 y | 3 stages | 6 | 6 | 0.77 ± 0.11 | 0.86/0.70/0.87 | 0.91/0.79/0.82 | 0.82/0.64/0.93 |
| **Human** | 90+ y | 3 stages | 5 | 5 | 0.76 ± 0.14 | 0.83/0.60/0.84 | 0.93/0.58/0.81 | 0.76/0.69/0.88 |
| ***4 stages scoring (WAKE / REM / N1 + N2 / N3)*** | | | | | | | | |
| **Human** | 5-7.5 y | 4 stages | 10 | 10 | 0.75 ± 0.08 | 0.74/0.77/0.74/0.77 | 0.75/0.72/0.78/0.85 | 0.80/0.86/0.73/0.75 |
| **Human** | 7.5-10 y | 4 stages | 10 | 10 | 0.80 ± 0.06 | 0.78/0.78/0.78/0.86 | 0.75/0.71/0.82/0.93 | 0.87/0.89/0.76/0.81 |
| **Human** | 17 y | 4 stages | 15 | 15 | 0.83 ± 0.06 | 0.87/0.83/0.80/0.83 | 0.84/0.82/0.84/0.85 | 0.92/0.85/0.77/0.85 |
| **Human** | 18 y | 4 stages | 5 | 5 | 0.80 ± 0.06 | 0.88/0.79/0.76/0.76 | 0.88/0.74/0.76/0.91 | 0.89/0.86/0.79/0.69 |
| **Human** | 70-80 y | 4 stages | 9 | 9 | 0.67 ± 0.13 | 0.75/0.76/0.78/0.43 | 0.87/0.80/0.80/0.37 | 0.70/0.74/0.76/0.88 |
| **Human** | 80-90 y | 4 stages | 6 | 6 | 0.62 ± 0.10 | 0.86/0.70/0.78/0.41 | 0.91/0.79/0.77/0.28 | 0.82/0.64/0.84/0.89 |
| **Human** | 90+ y | 4 stages | 5 | 5 | 0.65 ± 0.16 | 0.83/0.60/0.76/0.40 | 0.93/0.58/0.75/0.32 | 0.76/0.69/0.76/0.71 |
| ***5 stages scoring (WAKE / REM / N1 / N2 / N3)*** | | | | | | | | |
| **Human** | 5-7.5 y | 5 stages | 10 | 10 | 0.63 ± 0.09 | 0.74/0.77/0.21/0.71/0.77 | 0.75/0.72/0.20/0.80/0.85 | 0.80/0.86/0.29/0.66/0.75 |
| **Human** | 7.5-10 y | 5 stages | 10 | 10 | 0.69 ± 0.08 | 0.78/0.78/0.24/0.77/0.86 | 0.75/0.71/0.22/0.86/0.93 | 0.87/0.89/0.33/0.70/0.81 |
| **Human** | 17 y | 5 stages | 15 | 15 | 0.68 ± 0.05 | 0.87/0.83/0.13/0.78/0.83 | 0.84/0.82/0.09/0.90/0.85 | 0.92/0.85/0.29/0.71/0.85 |
| **Human** | 18 y | 5 stages | 5 | 5 | 0.64 ± 0.04 | 0.88/0.79/0.07/0.74/0.76 | 0.88/0.74/0.04/0.78/0.91 | 0.89/0.86/0.30/0.72/0.69 |

| Species | Group | Scheme | N individuals | N recordings | MF1 | F1 per stage | Precision per stage | Recall per stage |
|---|---|---|---|---|---|---|---|---|
| **Human** | 70-80 y | 5 stages | 9 | 9 | 0.56 ± 0.07 | 0.75/0.76/0.25/0.66/0.43 | 0.87/0.80/0.23/0.74/0.37 | 0.70/0.74/0.29/0.62/0.88 |
| **Human** | 80-90 y | 5 stages | 6 | 6 | 0.52 ± 0.08 | 0.86/0.70/0.18/0.75/0.41 | 0.91/0.79/0.17/0.79/0.28 | 0.82/0.64/0.28/0.73/0.89 |
| **Human** | 90+ y | 5 stages | 5 | 5 | 0.53 ± 0.08 | 0.83/0.60/0.16/0.67/0.40 | 0.93/0.58/0.13/0.78/0.32 | 0.76/0.69/0.24/0.61/0.71 |
| ***Birds (jackdaw)*** *(WAKE / REM / NREM)* | | | | | | | | |
| ***Per condition*** | | | | | | | | |
| **Jackdaw** | Baseline | Bilateral | 9 | 9 | 0.73 ± 0.04 | 0.95/0.43/0.82 | 0.92/0.41/0.88 | 0.98/0.49/0.77 |
| **Jackdaw** | SD | Bilateral | 9 | 9 | 0.63 ± 0.05 | 0.96/0.11/0.82 | 0.94/0.09/0.86 | 0.98/0.14/0.80 |
| **Jackdaw** | Recovery | Bilateral | 9 | 9 | 0.75 ± 0.04 | 0.94/0.48/0.81 | 0.91/0.43/0.89 | 0.98/0.56/0.76 |
| ***Per hemisphere*** | | | | | | | | |
| **Jackdaw** | All conditions | Left | 9 | 27 | 0.70 ± 0.07 | 0.95/0.34/0.82 | 0.92/0.30/0.88 | 0.98/0.42/0.77 |
| **Jackdaw** | All conditions | Right | 9 | 27 | 0.71 ± 0.07 | 0.95/0.35/0.82 | 0.93/0.31/0.87 | 0.98/0.42/0.77 |
| **Jackdaw** | All conditions | Combined | 9 | 27 | 0.70 ± 0.07 | 0.95/0.33/0.82 | 0.92/0.32/0.87 | 0.98/0.36/0.78 |

*Macro F1 (MF1), per-stage F1, precision, and recall are reported per group, scheme, and montage. MF1 is given as mean ± standard deviation across recordings; per-stage F1, precision, and recall are given as mean only. Per-stage values are reported in the stage order indicated by each sub-section header.*

##### Supplementary Table 11. Per-dataset agreement of Nyx and Somnotate with manual scoring across the lifespan.

| Dataset | N | Method | Accuracy (M ± SD) | Cohen's κ (M ± SD) | MF1 (M ± SD) | WAKE F1 (M ± SD) | NREM F1 (M ± SD) | REM F1 (M ± SD) |
|---|---|---|---|---|---|---|---|---|
| Development (rat; P12–adult) | 10 | *Nyx* | 0.908 ± 0.037 | 0.824 ± 0.055 | 0.874 ± 0.043 | 0.809 ± 0.086 | 0.940 ± 0.028 | 0.874 ± 0.062 |
| | | *Somnotate* | 0.700 ± 0.158 | 0.540 ± 0.221 | 0.691 ± 0.157 | 0.586 ± 0.232 | 0.750 ± 0.143 | 0.738 ± 0.155 |
| Aged (mouse; 14 months) | 2 | *Nyx* | 0.835 ± 0.028 | 0.683 ± 0.084 | 0.833 ± 0.034 | 0.872 ± 0.012 | 0.764 ± 0.072 | 0.864 ± 0.016 |
| | | *Somnotate* | 0.726 ± 0.068 | 0.504 ± 0.160 | 0.713 ± 0.104 | 0.779 ± 0.025 | 0.633 ± 0.145 | 0.728 ± 0.141 |

*Values are mean ± standard deviation across recordings of accuracy, Cohen's κ, macro-F1 (MF1) and per-stage F1 (WAKE, NREM, REM) for Nyx and Somnotate relative to manual expert scoring on developmental (rat; postnatal days P12–adult) and aged (mouse; 14 months) recordings. Somnotate was trained exclusively on its original mouse dataset (six recordings from the Oxford Mouse Benchmark) and applied without retraining. N, number of paired recordings per dataset.*

##### Supplementary Table 12. Region-specific comparison of Nyx performance across cortical electrodes.

| Metric | n pairs | Frontal median | Parietal median | W | p (raw) | p (Holm) | Effect size (r) |
|---|---|---|---|---|---|---|---|
| MF1 | 14 | 0.886 | 0.902 | 3.0 | <0.001 | **0.004** | -0.943 |
| Accuracy | 14 | 0.927 | 0.927 | 13.0 | 0.011 | 0.032 | -0.752 |
| Cohen's κ | 14 | 0.844 | 0.849 | 10.0 | 0.005 | 0.021 | -0.81 |
| F1 REM | 14 | 0.83 | 0.867 | 5.0 | 0.001 | **0.006** | -0.905 |
| F1 WAKE | 14 | 0.919 | 0.899 | 12.0 | 0.214 | 0.214 | -0.467 |
| F1 NREM | 14 | 0.938 | 0.941 | 14.0 | 0.013 | 0.032 | -0.733 |

*Parietal vs frontal EEG channel performance (Wilcoxon signed-rank test). Metrics are reported as median values across n = 14 paired recordings (7 DSI mice, both channels). Effect size r = Z / √n. Significant p (Holm) values (< 0.05) are bolded. W, Wilcoxon test statistic; κ, Cohen's kappa.*

##### Supplementary Table 13. Per-recording classification performance of Nyx across electrode derivations on n = 4 in-house rat recordings with paired modalities.

| Derivation | N recordings | MF1 | Accuracy | Cohen's κ | Precision (macro) | Recall (macro) |
|---|---|---|---|---|---|---|
| EEG frontal | 4 | 0.863 ± 0.028 | 0.919 ± 0.021 | 0.814 ± 0.050 | 0.890 | 0.855 |
| EEG parietal | 4 | 0.876 ± 0.025 | 0.927 ± 0.015 | 0.833 ± 0.022 | 0.901 | 0.864 |
| LFP hippocampus | 4 | 0.835 ± 0.048 | 0.890 ± 0.026 | 0.742 ± 0.080 | 0.868 | 0.818 |
| PAR–PFC | 4 | 0.858 ± 0.012 | 0.912 ± 0.027 | 0.806 ± 0.019 | 0.881 | 0.853 |
| PAR–PFC (EMG-like) | 4 | 0.813 ± 0.058 | 0.884 ± 0.022 | 0.744 ± 0.067 | 0.811 | 0.846 |
| PAR–PFC (no EMG) | 4 | 0.712 ± 0.101 | 0.855 ± 0.036 | 0.679 ± 0.054 | 0.708 | 0.744 |

*MF1, macro F1. Values are mean ± standard deviation across recordings; precision and recall are macro-averaged across stages. Derivations compared: surface EEG (frontal, parietal), LFP (hippocampus), bipolar PAR–PFC with and without an EMG-like channel.*

#### Supplementary Table 14. Nyx scoring performance across intracortical LFP and EEG recordings.

##### *14a. Descriptive statistics per modality*

| Modality | N recordings | N channels | Pooled F1 (median) | Pooled F1 (mean ± SD) | Per-recording F1 median [IQR] | Per-recording SD of F median [IQR] |
|---|---|---|---|---|---|---|
| Intracortical | 6 | 30 | 0.811 | 0.769 ± 0.115 | 0.776 [0.686, 0.864] | 0.055 [0.031, 0.066] |
| EEG | 6 | 12 | 0.884 | 0.870 ± 0.032 | 0.865 [0.861, 0.871] | 0.040 [0.019, 0.050] |

##### *14b. Paired and marginal comparisons*

| Effect | Comparison | Contrast (sign) | Δ | 95% CI | p | |
|---|---|---|---|---|---|---|
| Level (median F1) | Paired within-recording | EEG − Intracortical (+ = EEG higher) | 0.093 | [0.026, 0.160] | 0.008 | ** |
| Level (median F1) | Marginal pooled | EEG – Intracortical (+ = EEG higher) | 0.073 | [0.011, 0.187] | 0.009 | ** |
| Spread (SD of F1) | Paired within-recording | Intracortical − EEG (+ = intra more variable) | 0.028 | [−0.003, 0.067] | 0.103 | n.s. |
| Spread (SD of F1) | Marginal pooled | Intracortical − EEG (+ = intra more variable) | 0.083 | [0.040, 0.102] | 0.012 | * |

*(A) Per-modality descriptive statistics. Pooled F1 aggregates all epochs across channels within each modality; per-recording summaries first compute median or SD of F1 across channels within each recording, then report the median and interquartile range across recordings. (B) Paired within-recording (primary) and marginal pooled (confirmatory) comparisons between modalities. Δ denotes the observed effect under the stated sign convention; 95% CIs are bootstrap intervals; two-sided p-values are reported without correction across the four tests. ** $p < 0.01$; * $p < 0.05$; n.s. not significant.*

**Supplementary Table 15. Classification accuracy, cluster quality and cluster-number selection across epoch lengths and exploratory non-mammalian recordings (Fig. S8).**

***15a. Rodent epoch-length sensitivity — classification metrics (n = 12 Oxford recordings).***

| Epoch (s) | n | Accuracy | Cohen's κ | Macro F1 | F1 per stage (W / REM / NREM) |
|---|---|---|---|---|---|
| 1 | 12 | 0.928 ± 0.020 | 0.868 ± 0.038 | 0.910 ± 0.022 | 0.922 / 0.873 / 0.936 |
| 2 | 12 | 0.927 ± 0.021 | 0.867 ± 0.040 | 0.910 ± 0.022 | 0.920 / 0.875 / 0.936 |
| 4 | 12 | 0.926 ± 0.021 | 0.864 ± 0.040 | 0.908 ± 0.021 | 0.920 / 0.869 / 0.934 |
| 5 | 12 | 0.915 ± 0.030 | 0.845 ± 0.054 | 0.896 ± 0.028 | 0.908 / 0.855 / 0.925 |
| 10 | 12 | 0.894 ± 0.043 | 0.811 ± 0.068 | 0.864 ± 0.059 | 0.888 / 0.785 / 0.918 |

***15b. Rodent epoch-length sensitivity — cluster-quality metrics.***

| Epoch (s) | n | Bootstrap ARI | Silhouette (norm.) | Davies–Bouldin (Δ) | Calinski–Harabasz (norm.) |
|---|---|---|---|---|---|
| 1 | 12 | 0.886 ± 0.159 | 1.002 ± 0.094 | -0.008 ± 0.150 | 1.031 ± 0.108 |
| 2 | 12 | 0.855 ± 0.143 | 0.976 ± 0.066 | -0.006 ± 0.039 | 1.012 ± 0.075 |
| 4 | 12 | 0.891 ± 0.062 | 0.996 ± 0.069 | -0.007 ± 0.066 | 1.004 ± 0.080 |
| 5 | 12 | 0.896 ± 0.071 | 1.003 ± 0.056 | -0.000 ± 0.041 | 1.015 ± 0.074 |
| 10 | 12 | 0.896 ± 0.051 | 1.000 ± 0.000 | 0.000 ± 0.000 | 1.000 ± 0.000 |

***15c. Human epoch-length sensitivity — 5-stage classification metrics (n = 12 DOD-H recordings).***

| Epoch (s) | n | Accuracy | Cohen's κ | Macro F1 | F1 per stage (W / REM / N1 / N2 / N3) |
|---|---|---|---|---|---|
| 5 | 12 | 0.724 ± 0.068 | 0.620 ± 0.097 | 0.632 ± 0.068 | 0.717 / 0.761 / 0.170 / 0.777 / 0.751 |
| 10 | 12 | 0.725 ± 0.078 | 0.629 ± 0.104 | 0.646 ± 0.084 | 0.702 / 0.800 / 0.192 / 0.773 / 0.770 |
| 15 | 12 | 0.750 ± 0.057 | 0.657 ± 0.078 | 0.661 ± 0.058 | 0.715 / 0.806 / 0.178 / 0.801 / 0.813 |
| 30 | 12 | 0.716 ± 0.077 | 0.618 ± 0.097 | 0.634 ± 0.067 | 0.653 / 0.810 / 0.157 / 0.770 / 0.796 |

***15d. Human epoch-length sensitivity — 4-stage classification metrics (N1 excluded).***

| Epoch (s) | n | Accuracy | Cohen's κ | Macro F1 | F1 per stage (W / REM / N2 / N3) |
|---|---|---|---|---|---|
| 5 | 12 | 0.759 ± 0.066 | 0.641 ± 0.099 | 0.747 ± 0.067 | 0.717 / 0.761 / 0.764 / 0.751 |
| 10 | 12 | 0.778 ± 0.062 | 0.671 ± 0.097 | 0.768 ± 0.083 | 0.702 / 0.800 / 0.795 / 0.770 |
| 15 | 12 | 0.806 ± 0.054 | 0.705 ± 0.093 | 0.789 ± 0.067 | 0.715 / 0.806 / 0.823 / 0.813 |
| 30 | 12 | 0.776 ± 0.055 | 0.667 ± 0.082 | 0.762 ± 0.052 | 0.653 / 0.810 / 0.792 / 0.796 |

##### *15e. Human epoch-length sensitivity — 3-stage classification metrics.*

| Epoch (s) | n | Accuracy | Cohen's κ | Macro F1 | F1 per stage (W / REM / NREM) |
|---|---|---|---|---|---|
| 5 | 12 | 0.845 ± 0.055 | 0.699 ± 0.110 | 0.790 ± 0.078 | 0.717 / 0.761 / 0.891 |
| 10 | 12 | 0.852 ± 0.063 | 0.719 ± 0.118 | 0.800 ± 0.094 | 0.702 / 0.800 / 0.897 |
| 15 | 12 | 0.868 ± 0.040 | 0.742 ± 0.089 | 0.810 ± 0.077 | 0.715 / 0.806 / 0.910 |
| 30 | 12 | 0.838 ± 0.044 | 0.693 ± 0.080 | 0.783 ± 0.053 | 0.653 / 0.810 / 0.885 |

##### *15f. Human epoch-length sensitivity — cluster-quality metrics.*

| Epoch (s) | n | Bootstrap ARI | Silhouette (norm.) | Davies–Bouldin (Δ) | Calinski–Harabasz (norm.) |
|---|---|---|---|---|---|
| 5 | 12 | 0.860 ± 0.104 | 1.046 ± 0.186 | -0.034 ± 0.219 | 0.966 ± 0.174 |
| 10 | 12 | 0.888 ± 0.108 | 1.045 ± 0.196 | 0.040 ± 0.194 | 1.027 ± 0.209 |
| 15 | 12 | 0.913 ± 0.120 | 1.034 ± 0.151 | 0.050 ± 0.125 | 1.023 ± 0.131 |
| 30 | 12 | 0.877 ± 0.091 | 1.000 ± 0.000 | 0.000 ± 0.000 | 1.000 ± 0.000 |

*Aggregate metrics reported as mean ± standard deviation across recordings. Per-stage F1 columns show mean values only (stage order: W / REM / NREM, with NREM split where applicable). Silhouette score is normalised within recording relative to the longest epoch length (20 s rodent, 30 s human) as a ratio (value / value_ref), so that higher values indicate better clustering for all three indices.*

#### Supplementary Table 16. Cluster-number selection for non-mammalian exploratory recordings (Fig. 6).

##### Jackdaw

| K | Bootstrap ARI (mean ± SD) | Gap statistic (value ± SE) |
|---|---|---|
| 2 | 0.999 ± 0.001 | 1.214 ± 0.021 |
| 3 | 0.982 ± 0.040 | 1.085 ± 0.016 |
| 4 | 0.862 ± 0.144 | 1.103 ± 0.015 |
| **5** | **0.991 ± 0.003** | **1.177 ± 0.004** |
| 6 | 0.846 ± 0.128 | 1.177 ± 0.011 |
| 7 | 0.912 ± 0.121 | 1.182 ± 0.008 |
| 8 | 0.811 ± 0.159 | 1.147 ± 0.002 |

##### Tegu lizard

| K | Bootstrap ARI (mean ± SD) | Gap statistic (value ± SE) |
|---|---|---|
| 2 | 0.957 ± 0.026 | 1.460 ± 0.020 |
| 3 | 0.628 ± 0.356 | 1.423 ± 0.008 |
| **4** | **0.983 ± 0.006** | **1.486 ± 0.019** |
| 5 | 0.983 ± 0.004 | 1.498 ± 0.006 |
| 6 | 0.967 ± 0.008 | 1.499 ± 0.008 |
| 7 | 0.854 ± 0.142 | 1.511 ± 0.006 |
| 8 | 0.710 ± 0.108 | 1.504 ± 0.010 |

*Aggregate metrics reported as mean ± standard deviation across recordings. Bootstrap ARI computed from 190 pairwise comparisons across 20 × 80% subsamples (without replacement, fixed per-draw seeds); gap statistic computed against B = 50 uniform reference datasets drawn from the feature-space bounding box. Selected K shown in bold.*

#### Supplementary Methods

##### Spectrogram parameters

Spectrograms were computed using scipy.signal.spectrogram with a Hanning window and the parameters listed below. Temporal smoothing was applied post-hoc by convolving adjacent spectrogram columns with a Gaussian kernel of the specified width.

Spectrogram computation parameters by signal and species.

| Parameter | Human EMG | Human EEG | Rodent EMG | Rodent EEG |
|---|---|---|---|---|
| Frequency band (Hz) | 10-100 | 0.3-35 | 30-100 | 0.5-40 |
| Window length (s) | 30 | 30 | 4 | 2 |
| Overlap ratio | 0.5 | 0.5 | 0.5 | 0.5 |
| Temporal smoothing (s) | 30 | 30 | 4 | 4 |
| Spectral scaling | Density | Density | Density | Density |
| Detrend | Constant | Constant | Constant | Constant |
| Power scale | dB | dB | dB | dB |
| Normalisation | z-score | z-score | Mean subtraction | Mean subtraction |

For human recordings, each frequency bin of the EEG spectrogram was z-scored independently across time (zero mean, unit variance). For rodent recordings, the global mean spectrum was subtracted across time. Both approaches yield equivalent clustering results.

##### Clustering algorithm details

Three clustering algorithms are available as user-selectable options. For all methods, features were standardised to zero mean and unit variance before clustering. Outlier epochs (maximum absolute z-score > 6 across all features) were excluded from clustering and labelled NOSIGNAL.

###### K-means

Standard Lloyd's algorithm. The number of clusters *k* is specified by the user prior to each run. Cluster centroids are computed in the standardised PCA+EMG feature space and used both for visualisation and to sort clusters by their PC1 centroid value, which provides a first heuristic for stage assignment (lower PC1 typically corresponding to slow-wave activity).

###### Gaussian mixture models (GMM)

Full-covariance GMM fitted with the expectation–maximisation (EM) algorithm. The number of mixture components equals the number of clusters specified by the user. GMM is preferred when cluster boundaries are soft or non-spherical in the PCA space, as is common in human sleep data where stage transitions are gradual.

###### HDBSCAN

Hierarchical density-based spatial clustering of applications with noise. Key configurable parameters and their defaults for rodent recordings are: minimum cluster size = 100 epochs, minimum samples = 10. Epochs not assigned to any cluster receive a noise label and are mapped to NOSIGNAL. HDBSCAN does not require a pre-specified number of clusters and is robust to outliers; it is particularly suited to 24-hour rodent recordings in which stage proportions may be highly unequal across the light–dark cycle.

##### Minimum duration and post-processing

Following both the EMG-based WAKE/SLEEP classification and each clustering step, a minimum-duration constraint was applied to suppress spurious single-epoch stage transitions. In rodents transition to a new stage was accepted only if the new stage persisted for at least min_duration = 4 s .Isolated epochs that did not meet this criterion were absorbed into the surrounding stage.

Optional forbidden-transition rules (e.g., REM cannot directly follow WAKE without an intervening NREM epoch) can be applied as a post-processing step (see Post-hoc rules). These rules encode known physiological constraints on sleep architecture and can help resolve ambiguous cluster boundaries near stage transitions. Forbidden-transitions were not applied in the primary analyses reported in this study.

##### Evaluation metrics

Epoch-level agreement between automatically and manually scored hypnograms was computed after aligning both scorings to a common time grid at the spectrogram resolution. Epochs labelled NOSIGNAL in either scoring were excluded from all metric calculations.

Overall accuracy was defined as the proportion of epochs for which the automatic and manual labels agreed. Cohen's kappa ($\kappa$) was computed to quantify agreement beyond chance, accounting for all classes simultaneously:

$$\kappa = \frac{p_0 - p_e}{1 - p_e}$$

where $p_0 = \sum_k \frac{n_{kk}}{N}$ is the observed proportion agreement ($n_{kk}$= diagonal entries of the confusion matrix, *N* = total epochs), and $p_e = \sum_k \frac{\mathrm{row}_k/N}{\mathrm{col}_k/N}$is the expected chance agreement, with $\mathrm{row}_k$ and $\mathrm{col}_k$ denoting the marginal totals for stage $k$.

Per-class metrics for stage $k$ were computed from the confusion matrix as:

$$\text{Precision}_k = \frac{\mathrm{TP}_k}{\mathrm{TP}_k + \mathrm{FP}_k}$$

$$\text{Recall}_k = \frac{\mathrm{TP}_k}{\mathrm{TP}_k + \mathrm{FN}_k}$$

$$\text{F1}_k = \frac{2 \cdot \text{Precision}_k \cdot \text{Recall}_k}{\text{Precision}_k + \text{Recall}_k}$$

where $\mathrm{TP}_k$, $\mathrm{FP}_k$, $\mathrm{FN}_k$ are true positives, false positives, and false negatives for stage $k$, respectively. The macro-averaged F1 (MF1) is the unweighted mean of per-stage F1 scores. Unlike accuracy or weighted-averaged F1, which are dominated by the most prevalent stages (NREM in rodents, N2 in humans), MF1 weights each stage equally and is therefore sensitive to performance on rare stages such as REM and N1.

Confusion matrices (rows = true label, columns = predicted label) are presented as row-normalised percentages (recall per stage).

##### Inter-scorer agreement

Five analyses were performed on the in-house Boccara DSI telemetry dataset, following Brodersen et al. (2024) (Fig. S2). First, the fraction of epochs receiving each level of inter-scorer agreement (unanimous, majority, or no majority) was computed per consensus stage, providing a basic measure of how often scorers reached consensus. Second, per-scorer accuracy and Cohen's κ were computed against a leave-one-out (LOO) majority consensus of the remaining four scorers, providing a per-scorer summary of agreement with the group. Third, per-stage F1 score was computed for each of the ten scorer pairs, providing a stage-resolved view of pairwise agreement. Fourth, the rate at which each scorer disagreed with the LOO consensus was computed as a function of distance to the nearest stage transition, characterising whether disagreement concentrates around stage boundaries. Fifth, the agreement between subset-based consensuses and the full five-scorer consensus was computed as a function of subset size, assessing how many scorers are sufficient to approximate a stable ground truth. The automated scoring algorithm (Nyx) was benchmarked against this manual reference on the same recordings, on accuracy, Cohen's κ, and the transition-proximity disagreement profile. Full computational details, including consensus construction and exclusion rules, are given in Supplementary Methods.

Individual scorer hypnograms for DOD-H and DOD-O were obtained from the accompanying evaluation repository (https://github.com/Dreem-Organization/dreem-learning-evaluation). For the inter-scorer analysis, a 5-scorer majority-vote consensus was constructed independently for each dataset following the same procedure as for the DSI dataset (ties labelled as disagreement and excluded), rather than using the top-4 soft-agreement consensuses distributed with the original datasets. All five analyses described above were computed per dataset separately, with the exception of per-stage F1 scores, which were pooled across DOD-H and DOD-O (Fig. S3). The consensuses distributed on Zenodo were used as ground truth for all algorithm-vs-manual comparisons.

**Benchmarking against a supervised algorithm (rodents)**

Comparing supervised and unsupervised sleep-scoring frameworks on equal terms is not straightforward: any supervised model trained on the same labelled recordings it is later evaluated on has direct access to the ground truth that Nyx, by design, never sees. To avoid this asymmetry, we benchmarked Nyx against Somnotate (Brodersen et al., 2024) — an LDA/HMM supervised scorer — trained exclusively on its original published dataset (six mouse recordings from the Oxford Mouse Benchmark; Zenodo DOI:10.5281/zenodo.10200481) and applied without retraining or fine-tuning to a subset of our rodent recordings. This matches Nyx's no-label-access condition and tests both frameworks' ability to generalise to data outside their fitting distribution. Somnotate was compared to manual expert scoring on 239 recordings from 35 animals across five non-developmental datasets (two rat, three mouse), and on 12 recordings from 8 animals for the lifespan analysis (developmental rats at P12/P18/P25/adult; 14-month aged mice). Agreement metrics were computed and Nyx and Somnotate were compared with a two-sided Wilcoxon signed-rank test on paired per-animal mean macro-F1 to avoid pseudoreplication, with effect size quantified as the matched-pairs rank-biserial correlation.

**Benchmarking against a supervised algorithm (humans)**

For human recordings, Nyx was benchmarked against YASA (Vallat & Walker, 2021), a widely used supervised sleep-scoring tool distributed as a pretrained model. As with the rodent benchmark, evaluation on data already seen during training would give the supervised comparator direct access to labels that Nyx never sees. YASA was therefore applied without retraining to ANPHY (n = 29 recordings) and SleePing (n = 28 recordings), neither of which was part of YASA's original training set. Both datasets were scored under 3-, 4-, and 5-stage AASM schemes, and Nyx and YASA outputs were compared to manual expert scoring on a duration-resolved timeline as described above. Nyx and YASA were compared with a two-sided Wilcoxon signed-rank test on per-recording macro-F1, tested pooled across datasets and separately within each dataset for each stage scheme, with effect size quantified as the matched-pairs rank-biserial correlation.

**Post-hoc rules**

Rules were evaluated separately in rodents and humans. Rules were applied to the automatic hypnogram only; the manual scoring was never modified. All metrics were recomputed from the ruled hypnograms at 1 s resolution in rodents and 15 s in humans. The rodent evaluation used 179 baseline recordings from 10 datasets (103 rat, 76 mouse). The human evaluation used 179 whole-night/nap recordings from five datasets (ANPHY, DOD-H, DOD-O, MESA, SleePing). The automatic five-stage output was aggregated to three stages (N1-N3 collapsed to NREM).

R1 reclassifies any REM segment flanked by WAKE on both sides to WAKE. R2 applies the same flank condition only to REM segments shorter than a duration threshold (20 s in rodents, 30 s in humans). R3 reclassifies REM immediately following a WAKE bout longer than the same thresholds to WAKE. R4 reclassifies REM immediately following any WAKE bout to WAKE. Three additional rules were tested in rodents only. R5 enforces a 4 s minimum bout length in the order REM, NREM, WAKE: a short REM or NREM segment is merged into its flanking stage when both neighbours match, otherwise into WAKE; a short WAKE segment is merged into NREM. R6 applies R4 followed by R5. R7 applies R5 then R4.

From the 1 s hypnograms we computed Total sleep time (%) and the REM/NREM ratio in rodents, and Sleep onset latency and REM latency in humans. Metrics undefined for a given recording were excluded pairwise: REM/NREM ratio required manual REM > 0; REM latency required scored REM in both hypnograms (humans). Each ruled hypnogram was compared to Standard (ST) per recording. Reclassified epochs were expressed as a percentage of the whole recording and, per baseline stage, as a percentage of that stage's total duration in the Standard hypnogram, pooled across recordings.

Paired per-recording comparisons used two-sided Wilcoxon signed-rank tests: on Δκ against 0 for agreement, and on automated minus manual against 0 for biological metrics.

#### Epoch-length sensitivity analysis

We explored how Nyx performance is modulated by the adopted time-frequency resolution by comparing the classifications obtained with different spectrogram window size or bin size. For rodents, the parietal EEG recordings of the twelve mice of the Oxford mouse benchmark (test and baseline sleep-deprivation sub-datasets) were reanalysed at five raw bin sizes — 2, 4, 8, 10 and 20 s, giving effective epoch resolutions of 1, 2, 4, 5 and 10 s. For humans, we randomly subsampled twelve good quality DOD-H recordings and reanalysed at four scored epoch lengths (effective 5, 10, 15 and 30 s). Because the human pipeline is hierarchical, the analysis was restricted to its multi-class sleep step — the direct analogue of the single sleep-stage clustering used in rodents — which partitions the SLEEP-epoch PCA space into the N1/N2/N3/REM sub-clusters; the upstream WAKE/SLEEP and WAKE/N1 steps were left at their pipeline defaults. The resulting hypnograms were compared against the manual scoring to compute classification performance across epoch lengths; for humans, this comparison was carried out under all three staging schemes (3-, 4- and 5-stage).

We additionally assessed how the EEG epoch length affects the internal quality and stability of the automatic sleep-stage clustering independently of the manual scoring. Cluster stability was quantified by a subsampling bootstrap and used as the primary, sample-size-unbiased measure. For each recording and epoch length, the clusters were recomputed with the exact algorithm, cluster number and component set that the pipeline used at that epoch length. Twenty subsamples were drawn at 80 % without replacement using fixed seeds, and each subsample was re-clustered with that same recipe. For every pair of subsamples the adjusted Rand index (ARI)32 was computed on the epochs the two subsamples shared, and the mean and standard deviation were taken over all 20 × 19 / 2 = 190 pairs. Higher mean ARI indicates cluster structure that is more reproducible under resampling. Because the bootstrap fixes the sample size by construction, this measure requires no further sample-size matching. The silhouette coefficient[1] was computed on the same PCA(+EMG) space as a supporting, comparability-controlled analysis. Because this index is sensitive to the number of points and to within-cluster spread, two controls were imposed. First, to isolate the effect of epoch length from differences in the clustering recipe, the reference recording (here the coarser at 20 s epoch length) configuration — its number of principal components and its number of clusters K — was held fixed across all epoch lengths and the clusters were obtained with a Gaussian mixture model of K components. Second, sample-size matching was applied: at every epoch length shorter than the reference the epochs were randomly subsampled (fixed seed) down to the reference epoch count before the indices were computed, so that all epoch lengths were evaluated on the same number of points. The number of principal components was asserted to be identical across epoch lengths within each recording before the index was computed. The index was additionally expressed as ratios to its 20 s value to summarise whether finer epochs improved or degraded the underlying structure. All measures were reported per recording (one trajectory per recording across epoch lengths). Analyses used scikit-learn (KMeans/GaussianMixture, HDBSCAN via the hdbscan package, and the cluster-validation metrics above).

#### Trajectory and distance analysis of state transitions in the Nyx PC space

Analyses use 179 baseline rodent EEG/EMG recordings (103 rat, 76 mouse). EMG band power (30–100 Hz; 4 s windows, 50% overlap, dB, 4 s smoothing) provided the wake/sleep gate where applicable. EEG spectrograms (0.5–40 Hz; 2 s windows, 50% overlap, dB, 4 s smoothing) were sampled at 1 Hz, giving one PC sample per 1 s epoch. PCA was fit on sleep epochs (5–6 components). The selected PCs (with EMG where applicable) were z-scored per recording to form the clustering substrate, and epochs with any feature $|z|$>threshold were removed. Clustering used each recording's saved recipe on this substrate. Where sleep had already been separated by an EMG threshold, WAKE bouts were projected into the sleep PC space post hoc; otherwise WAKE, NREM, and REM were clustered jointly and WAKE occupied the shared PC space. Geometric clusters (K=3–7) were collapsed to WAKE, NREM, or REM via each recording's saved cluster_to_stage map. Distance was computed as the Mahalanobis distance from each epoch to the centroid of its assigned state in this space; Mahalanobis distance accounts for the shape and spread of each state cloud and puts distances on a common scale across WAKE, NREM, and REM. For analyses pooled across recordings (panels B, C, F), the Mahalanobis distance was z-scored within recording to remove recording-level offsets and scale differences (raw Mahalanobis levels grow with the clustering-space dimensionality, which varies across recordings), so the shape of the peri-transition excursion is comparable across animals. Transitions were aligned on Nyx, with Manual shown alongside for comparison. For each transition, the distance trace across ±40 s was extracted and averaged across all transitions (event-triggered average) to give the peri-transition profile. Near-transition epochs are defined by $|\Delta t| \leq 6$ s and far-from-transition epochs by $|\Delta t| \geq 30$ s. Transition crossings were counted as the number of Nyx transitions within ±10 s of the anchor (the anchor is included, so crossings ≥1); stable transitions have ≤1 crossing and unstable transitions have ≥3.
